# A hybrid *in silico* approach reveals novel inhibitors of multiple SARS-CoV-2 variants

**DOI:** 10.1101/2021.06.04.447130

**Authors:** Sankalp Jain, Daniel C. Talley, Bolormaa Baljinnyam, Jun Choe, Quinlin Hanson, Wei Zhu, Miao Xu, Catherine Z. Chen, Wei Zheng, Xin Hu, Min Shen, Ganesha Rai, Matthew D. Hall, Anton Simeonov, Alexey V. Zakharov

**Affiliations:** National Center for Advancing Translational Sciences (NCATS), National Institutes of Health, 9800 Medical Center Drive, Rockville, MD, 20850, United States

**Keywords:** COVID-19, SARS-CoV-2, virtual screening, machine learning, pharmacophore modeling

## Abstract

The National Center for Advancing Translational Sciences (NCATS) has been actively generating SARS-CoV-2 high-throughput screening data and disseminates it through the OpenData Portal (https://opendata.ncats.nih.gov/covid19/). Here, we provide a hybrid approach that utilizes NCATS screening data from the SARS-CoV-2 cytophatic effect reduction assay to build predictive models, using both machine learning and pharmacophore-based modeling. Optimized models were used to perform two iterative rounds of virtual screening to predict small molecules active against SARS-CoV-2. Experimental testing with live virus provided 100 (~16% of predicted hits) active compounds (Efficacy > 30%, IC_50_ ≤ 15 μM). Systematic clustering analysis of active compounds revealed three promising chemotypes which have not been previously identified as inhibitors of SARS-CoV-2 infection. Further analysis identified allosteric binders to host receptor angiotensin-converting enzyme 2, which were able to inhibit the entry of pseudoparticles bearing spike protein of wild type SARS-CoV-2 as well as South African B.1.351 and UK B.1.1.7 variants.

---

In December 2019, a novel coronavirus strain SARS-CoV-2 began to spread in Wuhan, China^1^ and eventually led to an alarming global pandemic. As of May 2021, the pandemic has reached over 154 million cases and the resulting complications have caused more than 30 million deaths worldwide^2^. Numerous strategies have been employed to find a reliable COVID-19 therapy including vaccine development, drug repurposing, and developing novel small molecule SARS-CoV-2 inhibitors^3–6^. The FDA has now issued emergency use authorization for multiple vaccines; however, the outbreak is far from under control, especially due to the emergence of several SARS-CoV-2 variants. As per a recent CDC report, there are 13 variants, 5 of which are classified as variants of concern^7,8^.

At the beginning of the pandemic, the National Center for Advancing Translational Sciences (NCATS) started a COVID-19 drug repurposing campaign and created the OpenData Portal to make all data generated utilizing SARS-CoV-2 related assays publicly accessible^9^. The COVID-19 targeted high-throughput screening (HTS) campaigns at NCATS apply a wide range of biochemical and cell-based assays, including the cytopathic effect assay (CPE) of live SARS-CoV-2 in Vero-E6 cells^10^. More recently, NCATS included testing of therapeutics against different SARS-CoV-2 variants (https://opendata.ncats.nih.gov/variant/assays).

Drug discovery is a time- and resource-intensive process; virtual screening to identify small molecule protein modulators offers significant advantages, especially when used to complement traditional HTS methodology^11,12^. Multiple *in silico* studies have been reported which employ virtual screening of small molecule databases^13–18^. However, in the majority of published COVID-19-related virtual screening communications, hit compounds were not experimentally validated in SARS-CoV-2 assays or were not counterscreened for cytotoxicity, rendering the results inconclusive.

While many efforts are focused on repurposing existing drugs^19,20^, we performed a hybrid virtual screening of two in-house libraries (~140k compounds) in an effort to identify new chemotypes with antiviral activity and limited cytotoxicity, utilizing the NCATS publicly available screening data. This hybrid approach integrates quantitative structure-activity relationship (QSAR) and ligand-based pharmacophore modeling, followed by experimental testing of predicted hits in the CPE and cytotoxicity assays. We executed two iterative rounds of virtual screening; hit compounds identified in the first round were experimentally tested and these data were utilized to enrich the training dataset for the proposed hybrid approach used in the second round (**Fig. 1**).

**Fig. 1:**
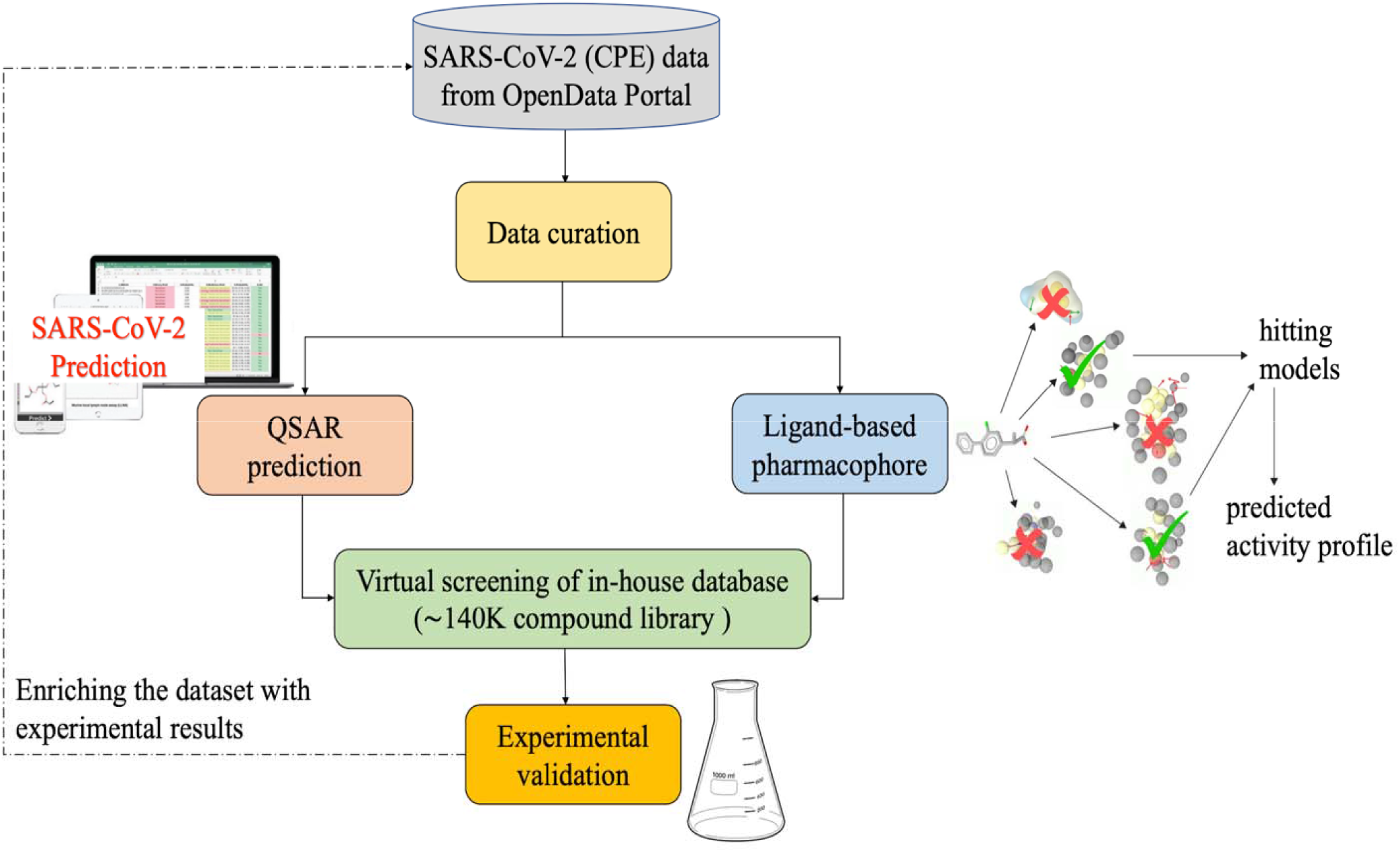
Flowchart of the virtual screening strategy adopted in this study.

These efforts resulted in a total of 100 compounds (out of 640 virtual screening hits; hit-rate ~16%) which showed inhibition (half-maximum inhibitory concentration, IC_50_ ≤ 15 μM; Efficacy > 30%) in the CPE assay and minimal cytotoxicity (IC_50_ > 30 μM), where 68 of them had an efficacy greater than 70%. Interestingly, three novel antiviral chemotypes emerged with multiple (≥3) active structural analogs in each cluster. Some preliminary structure-activity relationships (SARs) were identified within each cluster, which further validates these chemotypes as candidates for further medicinal chemistry optimization as novel SARS-CoV-2 inhibitors.

In an effort to elucidate the mechanism of action, hit compounds/chemotypes were tested across several viral targets. Six novel SARS-CoV-2 CPE inhibitors, identified as allosteric ACE2 binders (using microscale thermophoresis), blocked viral entry as assessed by a pseudoparticle entry assay (PP assay). In most cases, ACE2 binding showed a direct correlation to activity in the PP assay. We further validated these six novel inhibitors in PP assays using both the South African and the UK SARS-CoV-2 variants: two compounds showing submicromolar activity against both variants.

To the best of our knowledge, this is the first study that reveals novel inhibitors of multiple SARS-CoV-2 variants with elucidated mechanism of action. The curated dataset and optimized prediction models are publicly available via github (https://github.com/ncats/covid19_pred) as well as NCATS Predictor website (https://predictor.ncats.io/).

## Results

### Hybrid approach for *in silico* screening

Combination of ligand- and structure-based methods into consensus approach has been used earlier to discover novel actives against various targets^21–23^, since it improves the precision and reduces false positives^24^. In this study, we combined QSAR modeling with pharmacophore-based screening to identify novel chemotypes active against SARS-CoV-2. We used a consensus of the predictions based on the two approaches (**Fig. 1**) to select compounds for experimental validation.

#### Model Performance - Stratified Bagging

The performance of QSAR models based on different combinations of descriptors and stratified bagging (SB) approach is provided in Supplementary Table 1. It was measured by different metrics, including operating characteristic area under the curve (ROC AUC). All developed models showed ROC AUC values > 0.75. In the 1^st^ round of modeling, the consensus of descriptors (RDKit, Morgan and Avalon) provided the best performance with ROC AUC = 0.80 on the test set. SB models generated in the 1^st^ round of screening are referred to as SB-1. After obtaining the experimental results from the 1^st^ round of virtual screening, we updated our SB model (referred to as SB-2). In the 2^nd^ round of modeling, the consensus of descriptors (RDKit, Morgan and Avalon) showed improved results with ROC AUC = 0.84 on the test set.

#### Ligand-based Pharmacophore Modeling

For the 1^st^ round of ligand-based pharmacophore (LBP) modeling, we used the 48 active compounds: clustering based on pharmacophore-based similarity (cluster distance of 0.4, 0.6, 0.7, and 0.8), followed by generation of ligand-based hypotheses led to a total of 44 pharmacophore hypotheses (merged-features pharmacophore (MFP) and shared-features pharmacophore (SFP)). Taking the computational constraints into account, 15 pharmacophore models that hit the majority (> 20%) of active versus inactive (Supplementary Table 2) were selected for virtual screening. The ligand-based pharmacophore models generated in the first round of screening are referred to as LBP-1. For the 2^nd^ round, referred to as LBP-2, we considered 53 actives and followed the same protocol as above. This resulted in 55 pharmacophore hypotheses (MFP and SFP). Pharmacophore models (20) were then selected for virtual screening. All pharmacophore hypotheses generated in this study are presented in the supporting information (Supplementary Table 3). In general, the pharmacophoric sites such as hydrogen bond acceptor (HBA), hydrogen bond donor (HBD), aromatic ring, hydrophobic sites, and positive ionizable groups were prudently characterized.

#### Experimental Testing of 1^st^ Round *in silico* Screening Hits

The 320 compounds selected from the 1^st^ round of *in silico* screening (see Methods for details) were tested in the CPE assay in a 5-point dilution series, with concentrations ranging from 20 μM to 124 nM. To exclude compounds with cytotoxic effects, the compounds were counter-screened in a cell viability assay. Out of the 320 compounds tested, 46 compounds showed a SARS-CoV-2 CPE inhibiting activity with a maximum response (MaxResponse) greater than 30% and IC_50_ values of 3-15 μM. Of these 46, 42 compounds did not show any cytotoxicity or modest toxicity with an efficacy < 25% (Supplementary Table 4). This gives us a positive predicted value (PPV) (i.e., the fraction of model predicted positives that are experimentally confirmed) of 13%, which is 3-fold higher than the PPV calculated from the training set (PPV= 4%).

#### Experimental Testing of 2^nd^ Round *in silico* Screening Hits and Validation

The selected 320 compounds from the 2^nd^ round of *in silico* screening (see Methods for details) were also tested in the CPE assay in a 5-point dilution series, with concentrations ranging from 20 μM to 124 nM. For the 2^nd^ round testing, out of the 320 compounds, 65 compounds were identified with anti-SARS-CoV2-2 activity having a MaxResponse > 30% and IC_50_ values of 3-15 μM. Of these 65 compounds, 58 did not show any cytotoxicity or minimal toxicity (efficacy < 30%; Supplementary Table 5). Moreover, 27 of these 58 compounds exhibited an IC_50_ ≤ 10 μM in the CPE reduction assay. This further improved the PPV value to 18%.

For validation, 69 compounds were ‘cherry-picked’ and retested in the CPE reduction assay in duplicate as an 8-point dilution series, with concentrations ranging from 20 μM to 78 nM. 53 of the retested compounds were confirmed to have CPE inhibitory activity with an efficacy > 30% and IC_50_ of 5-25 μM, and five of them exhibited an IC_50_ ≤ 10 μM with no notable cytotoxicity (Supplementary Table 6). The five most potent and efficacious compounds (**1-5**) from the follow-up CPE assay are shown in **Fig. 2a**.

**Fig. 2:**
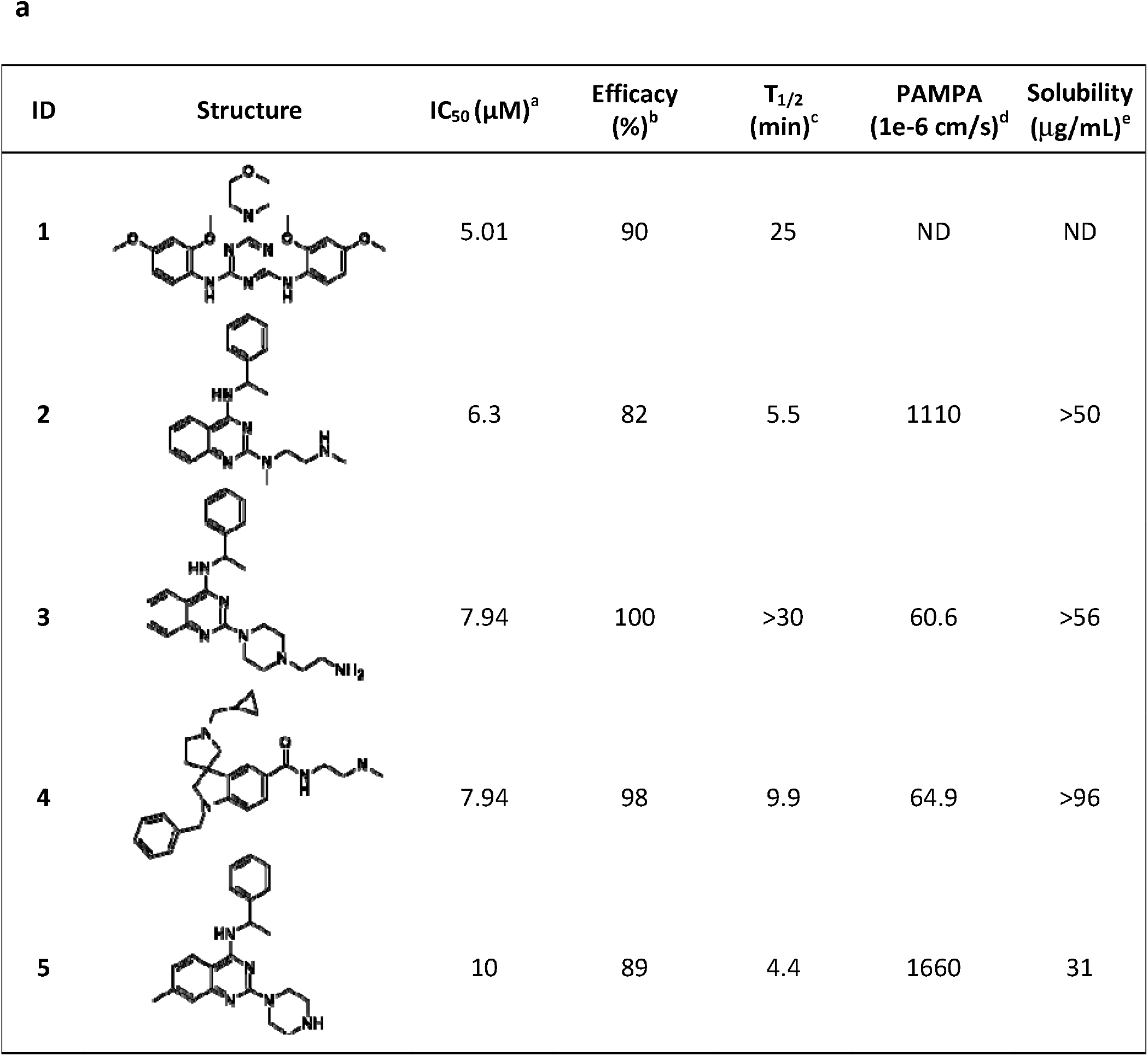
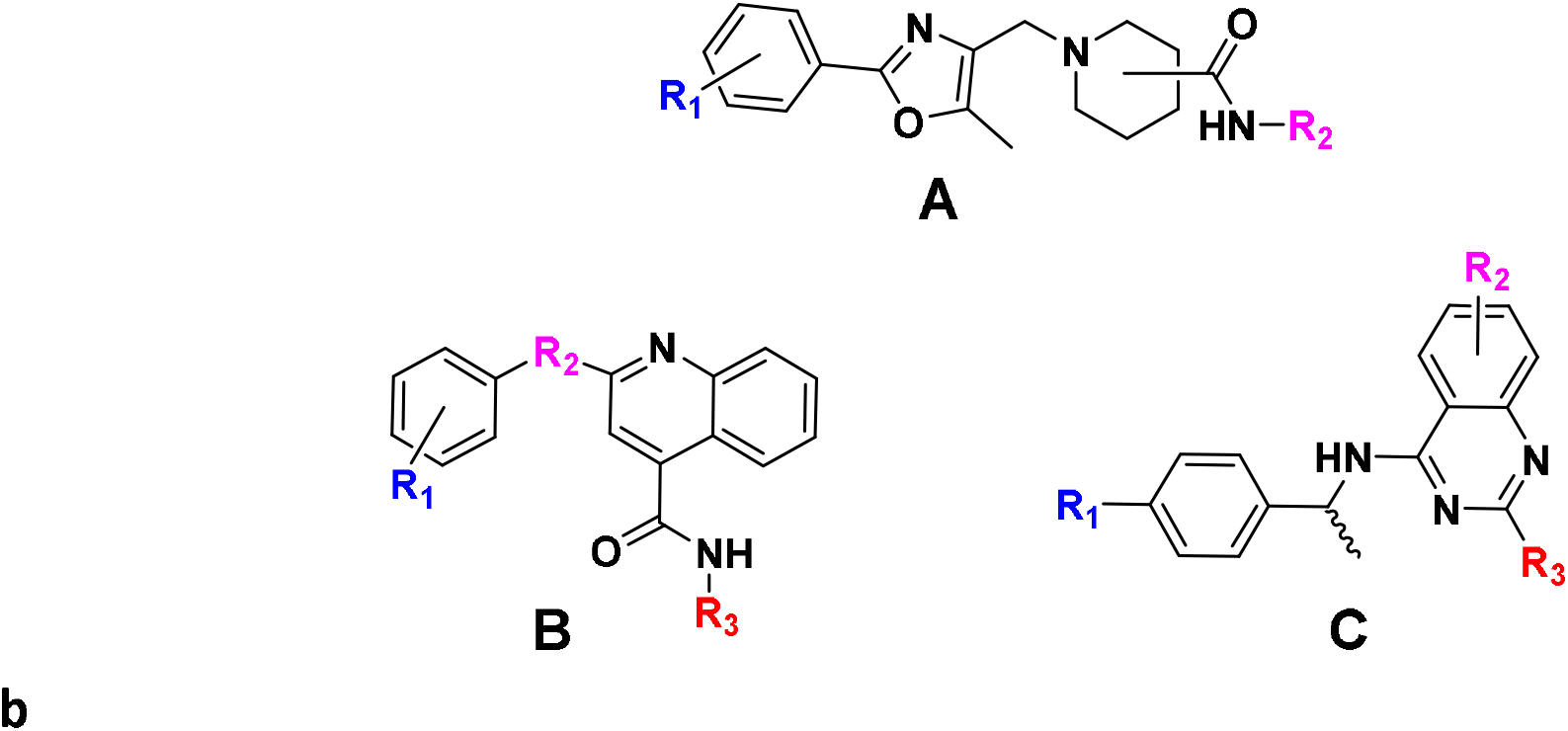
**a**. Five most potent and efficacious compounds identified, along with *in vitro*/physico-chemical ADME data. ^a^IC_50_: half-maximal inhibitory concentration value obtained from the CPE assay in 8-point dose response, measured in duplicate. ^b^Efficacy: maximum inhibitory effect observed in CPE assay. ^c^T_1/2_: metabolic half-life measured in rat liver microsome lysates reported in minutes, with a minimum detectable half-life of 1 minute. ^d^PAMPA (parallel artificial membrane permeation assay) is reported as a metric of the passive permeability of the compounds (1×10^−6^ cm/s). ^e^Solubility – pION μSOL assay for kinetic aqueous solubility determination, pH 7.4. **b.** Three chemotypes (**A-C**) identified as active in the CPE assay.

#### Clustering and Preliminary SAR Analysis

Hierarchical cluster analysis of the 100 CPE reducing compounds, identified in the two rounds of virtual screening, revealed three promising chemotypes (**Fig. 2b**), where three or more active structural analogs were identified (IC_50_ ≤ 15 μM and efficacy ≥ 70 %) and no notable cytotoxicity (IC_50_ ≤ 30 μM). Importantly, some preliminary structure-activity relationships (SAR) could be established for chemotypes where the analogs analyzed (and present in in-house compound libraries) were structurally similar enough for direct comparison. The most promising analogs from each of the three chemotypes, activity in the CPE assay, as well as *in vitro* physicochemical properties are shown in **Fig. 3** and **Table 1**.

**Fig. 3:**
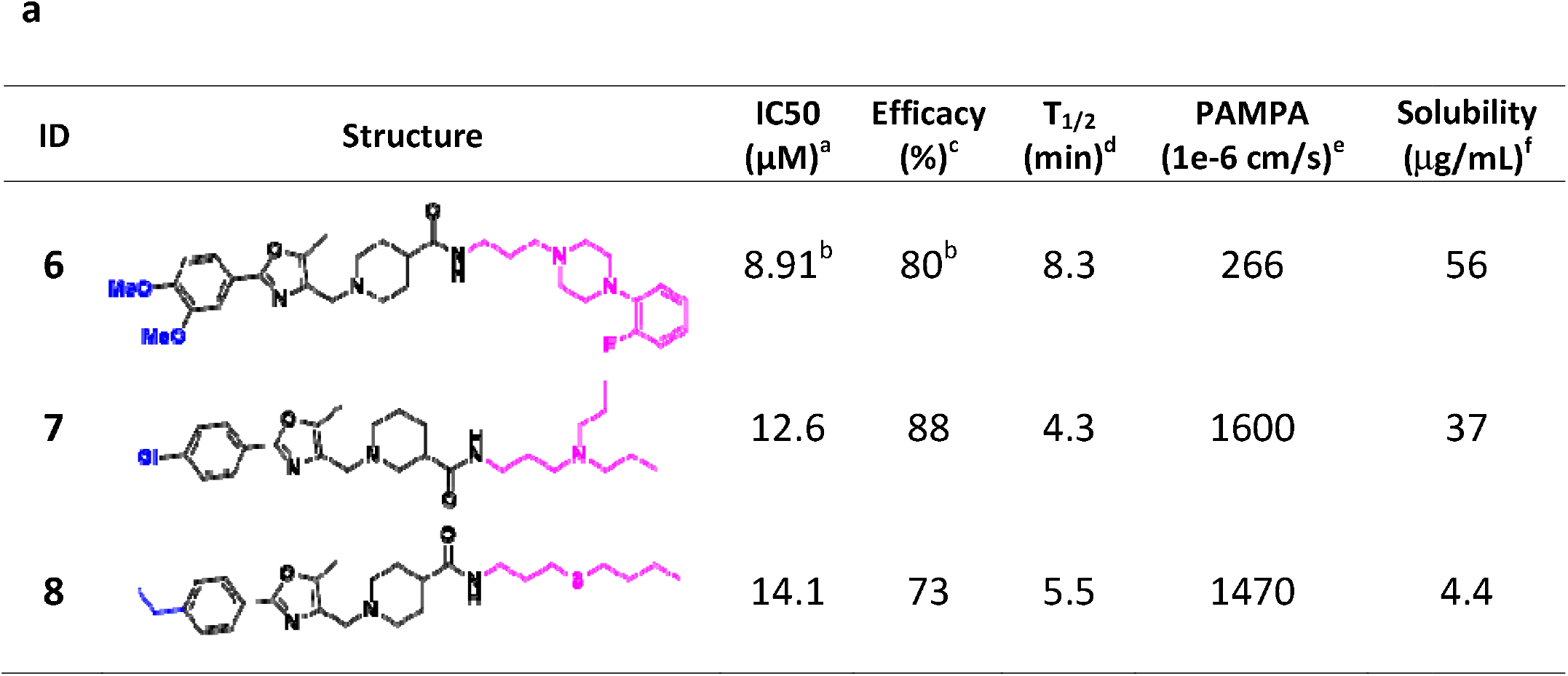
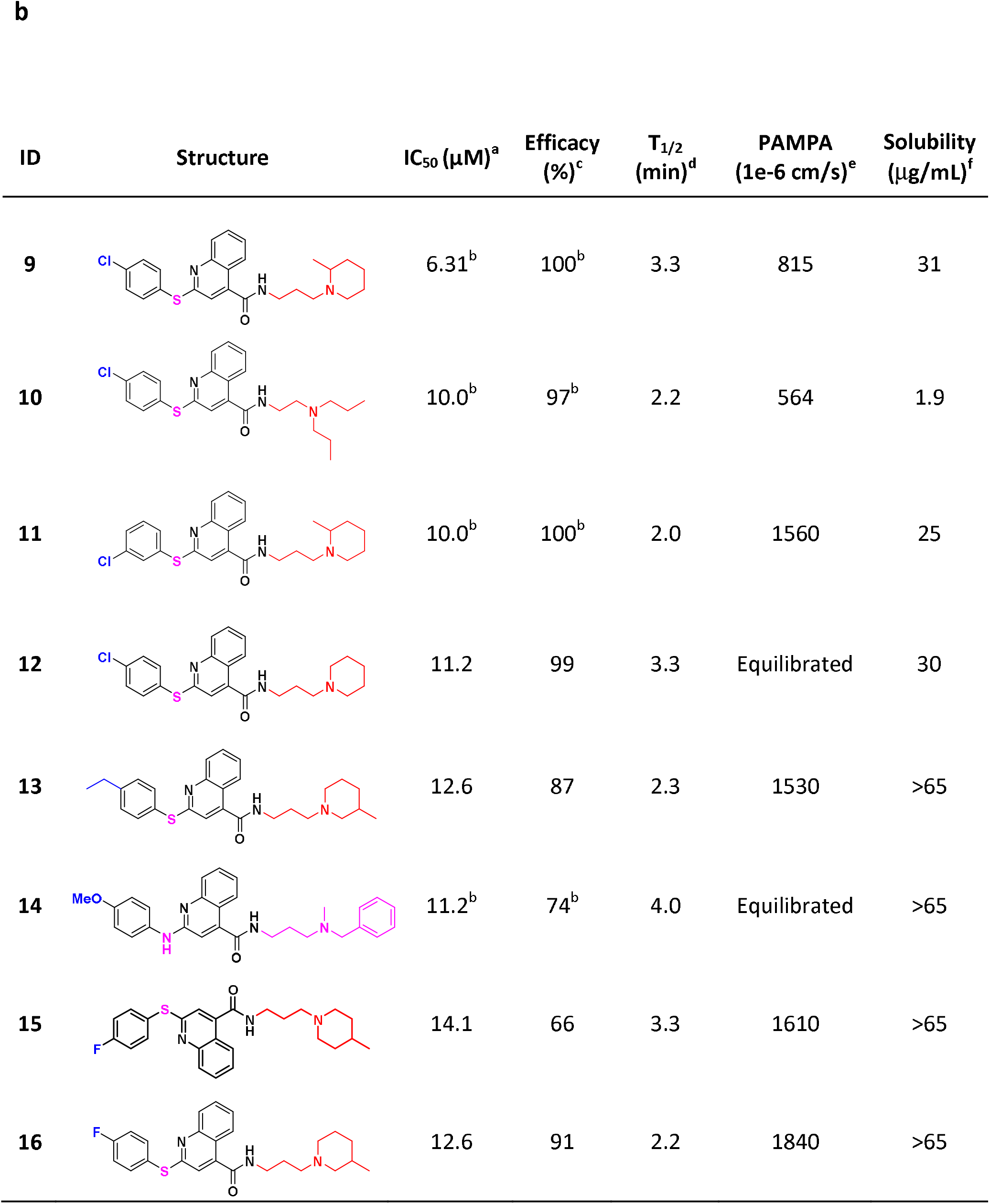
Notably active chemotypes **(a) A** and **(b) B** which show no notable cytotoxicity (IC_50_ ≤ 30 μM). ^a^IC_50_: half-maximal inhibitory concentration values obtained from the CPE assay in 8-point dose response, measured in duplicate. ^b^ Values represent data obtained from 5-point dose response, measured in duplicate. ^c^Efficacy: maximum inhibitory effect observed in CPE assay. ^d^T_1/2_: metabolic half-life measured in rat liver microsome lysates reported in minutes, minimum detectable half-life of 1 minute. ^e^PAMPA (parallel artificial membrane permeation assay) is reported as a metric of the passive permeability of the compounds. ^f^Solubility – pION μSOL assay for kinetic aqueous solubility determination, pH 7.4.

**Table 1.**
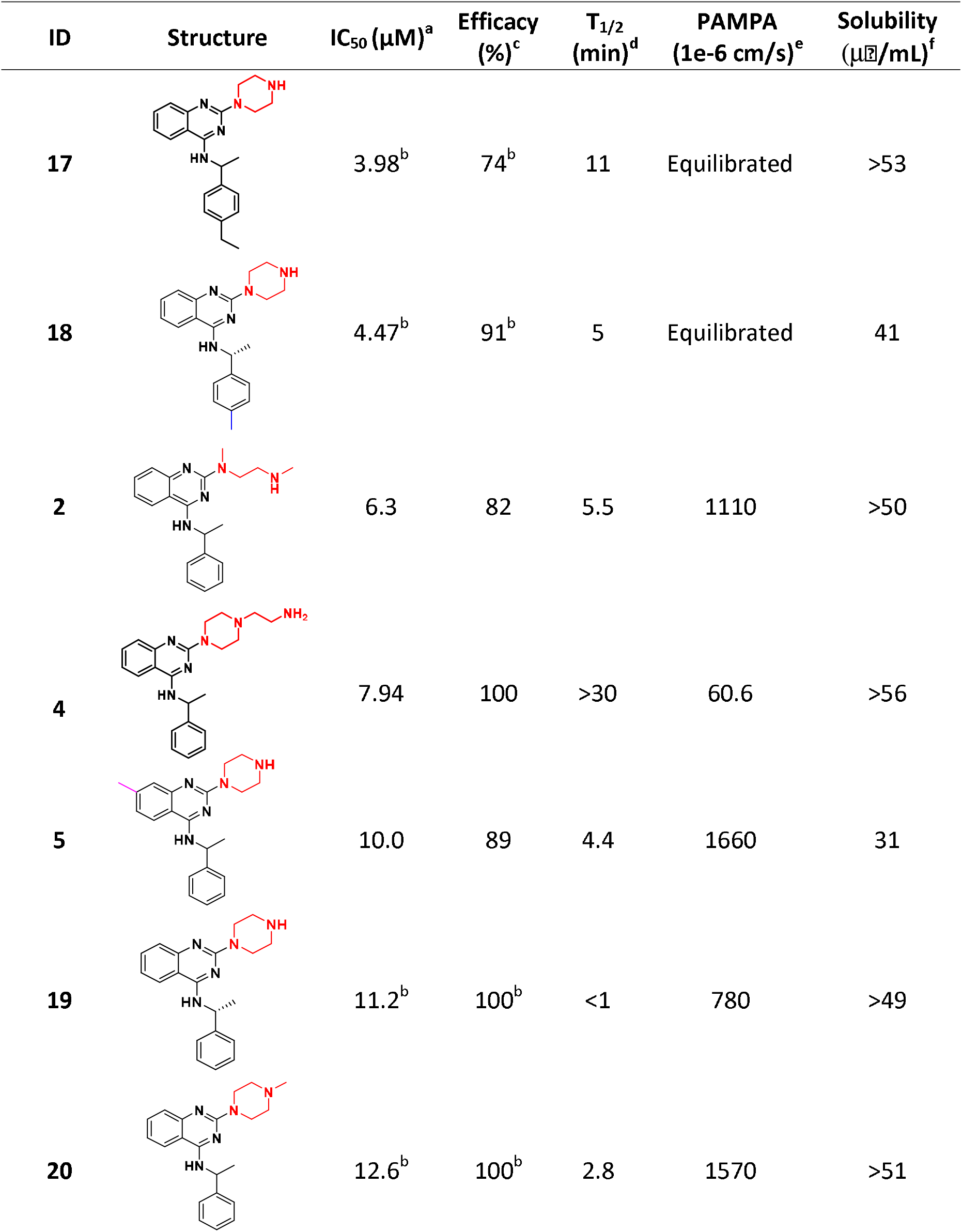
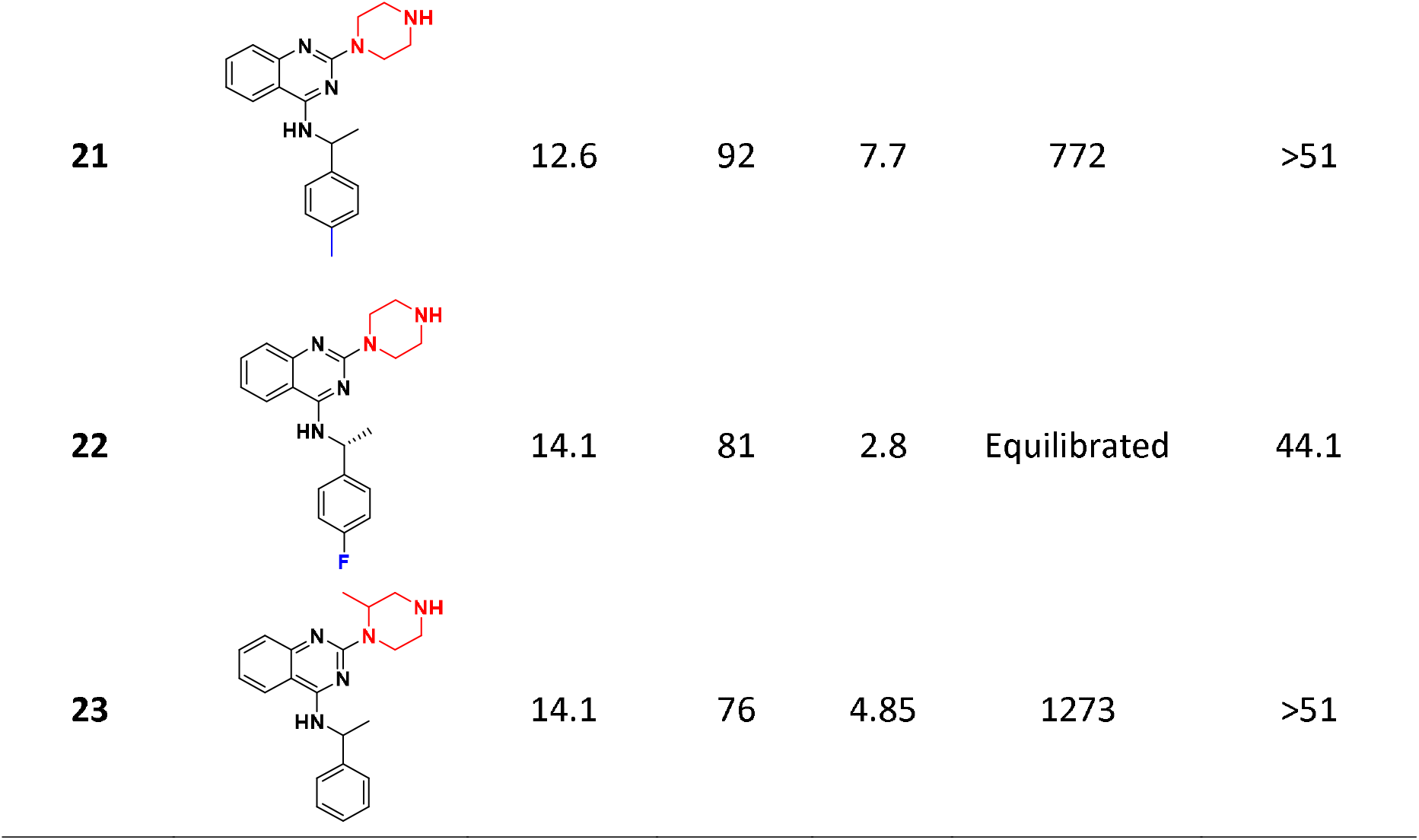
Notably active chemotype **C** which show no notable cytotoxicity (IC_50_ ≤ 30 μM). ^a^IC_50_: half-maximal inhibitory concentration values obtained from the CPE assay in 8-point dose response, measured in duplicate. ^b^ Values represent data obtained from 5-point dose response, measured in duplicate. **c**Efficacy: maximum inhibitory effect observed in CPE assay. ^d^T_1/2_: metabolic half-life measured in rat liver microsome lysates reported in minutes, minimum detectable half-life of 1 minute. ^e^PAMPA (parallel artificial membrane permeation assay) is reported as a metric of the passive permeability of the compounds. ^f^Solubility. pION μSOL assay for kinetic aqueous solubility determination, pH 7.4.

| ID | Structure | IC<sub>50</sub> (μM)<sup>a</sup> | Efficacy (%)<sup>c</sup> | T<sub>1/2</sub> (min)<sup>d</sup> | PAMPA (1e-6 cm/s)<sup>e</sup> | Solubility (μg/mL)<sup>f</sup> |
| --- | --- | --- | --- | --- | --- | --- |
| 17 |  | 3.98<sup>b</sup> | 74<sup>b</sup> | 11 | Equilibrated | >53 |
| 18 |  | 4.47<sup>b</sup> | 91<sup>b</sup> | 5 | Equilibrated | 41 |
| 2 |  | 6.3 | 82 | 5.5 | 1110 | >50 |
| 4 |  | 7.94 | 100 | >30 | 60.6 | >56 |
| 5 |  | 10.0 | 89 | 4.4 | 1660 | 31 |
| 19 |  | 11.2<sup>b</sup> | 100<sup>b</sup> | <1 | 780 | >49 |
| 20 |  | 12.6<sup>b</sup> | 100<sup>b</sup> | 2.8 | 1570 | >51 |
| 21 |  | 12.6 | 92 | 7.7 | 772 | >51 |
| 22 |  | 14.1 | 81 | 2.8 | Equilibrated | 44.1 |
| 23 |  | 14.1 | 76 | 4.85 | 1273 | >51 |

Within chemotype **A**, 26 analogs were tested from internal compound libraries and upon screening, three (compounds **6-8**) have IC_50_ values ranging from 8.9 - 14.1 μM and efficacy ≥ 83% (**Fig. 3a**). Although conclusive SAR trends were limited, the anti-SARS-CoV-2 activity (and cytotoxicity) is sensitive to substitutions on the phenyl ring attached to the oxazole and both the position (3- vs 4-) and structure of the piperidinyl-amide.

Within chemotype **B** were 18 in-house structural analogs, eight of which (compounds **9-16**) have promising IC_50_ and efficacy values (**Fig. 3b**). Analogs of this cluster were too structurally very diverse to analyze for conclusive SAR trends. With the exception of **10** which suffers from poor solubility, CPE-actives from this series have favorable solubility and permeability. Similar to chemotype **A**, all suffer from short metabolic half-time (T_1/2_) in rat liver microsomes.

Quinazoline-containing chemotype C provided 10 promising analogs (compounds **2, 4, 5, 17, 18, 20-23**), including three (**2, 4, 5**) of the most active compounds identified in the study (**Table 1**). Most of the notably active analogs contain variously substituted piperazines at the 2-position of the quinazoline core and 2-methyl-benzylamine at the 4-position. However, the open-chain (vs piperazine) analog (**2**) is also quite active, suggesting a 2-position diamine with an ethylene spacer is perhaps part of the parent pharmacophore. Methyl- and ethyl- substitutions off the 4-position of the benzylamine phenyl ring (**18** and **17**, respectively) were well tolerated, while 4-fluoro (**22**) and 4-phenyl substituents reduced the activity. Similar to chemotypes **A** and **B**, this series has favorable solubility and permeability but suffers from poor metabolic T_1/2_. However, it seems the metabolic liability can be mitigated *via* addition of an *N*-aminoethyl group off the piperazine ring (**4**; T_1/2_ > 30 min).

#### Mechanism of Action Studies

In efforts to elucidate the mechanism of action of virtual screening hits active in the CPE reducing assay, compounds were tested for their activity against some key events necessary for viral entry and replication. The SARS-CoV-2 main protease (M^pro^) represents an attractive target for anti-viral drug development because its inhibition prevents the formation of mature functional viral proteins and thus, viral replication^25^. As such, active compounds were screened in the SARS-CoV-2 M^pro^ enzymatic assay, however no activity was observed.

Compounds were also tested for their ability to interrupt the binding of the SARS-CoV-2 receptor binding domain (RBD) of the spike protein to the host receptor ACE2 using an AlphaLisa proximity assay, in combination with a counter-assay, to identify false positive hits. The compounds showed activity in both RBD-ACE2 AlphaLisa and in the TruHit counterscreen (see Methods for details), rendering to inconclusive results.

Nonetheless, we tested the compounds for their binding to ACE2 using microscale thermophoresis (MST) followed by a SARS-CoV-2 pseudoparticle (PP) entry assay, to explore if a molecule interacting with the viral receptor could interfere with host cell entry. In parallel, the compounds were tested in an ACE2 enzymatic assay. Six compounds were identified as ACE2 binders with an equilibrium dissociation constant (K_d_) ≤ 20 μM, with no inhibitory or agonistic activity in ACE2 enzymatic assay (**Table 2**).

**Table 2.**
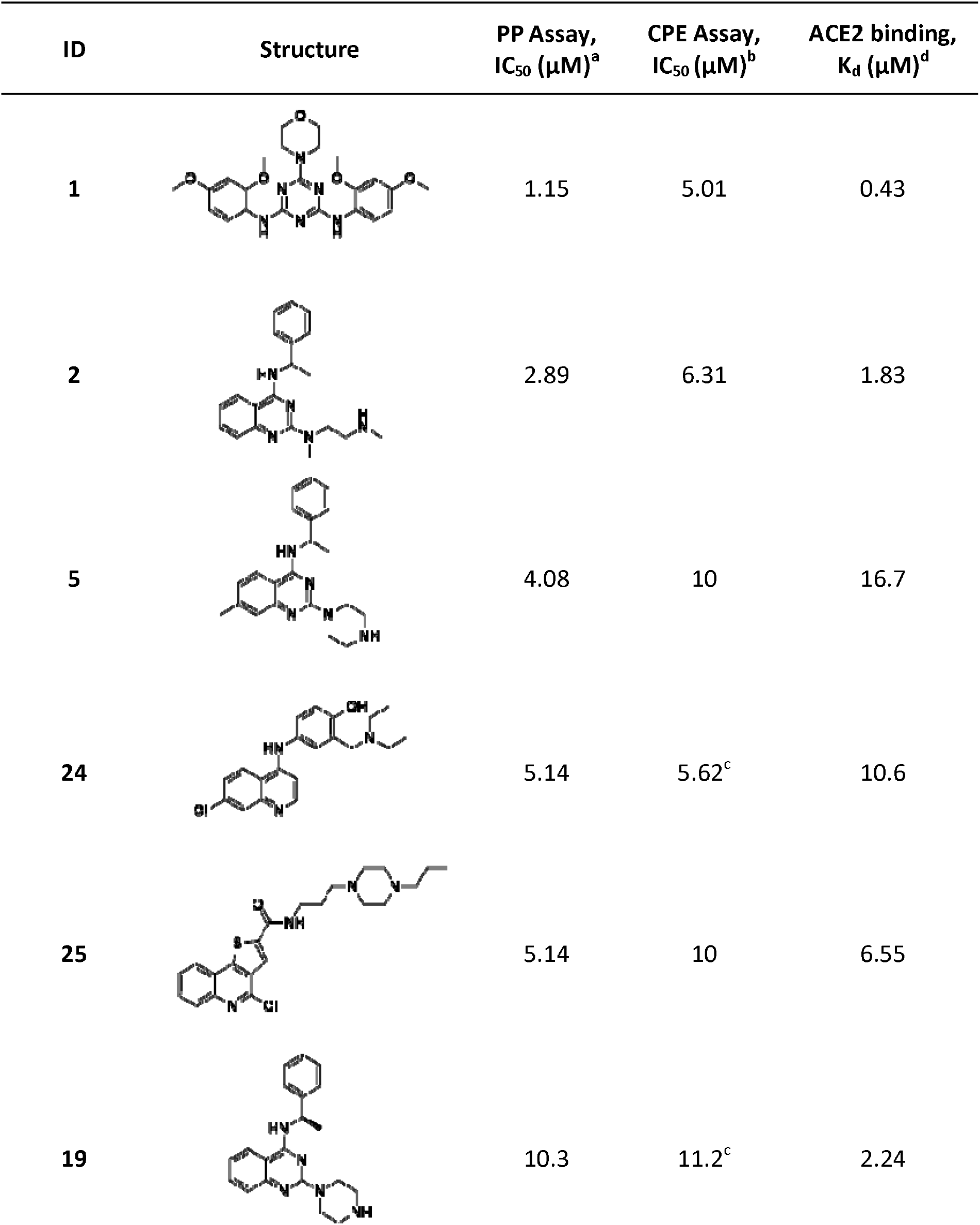
Compounds identified as ACE2 binders and inhibitors of viral entry in PP assay. aActivity in the SARS-CoV-2 PP assay. ^b^IC_50_: half maximal inhibitory concentration values obtained from the CPE assay in 8-point dose response, measured in duplicate. ^c^Values represent data obtained from 5-point dose response, measured in duplicate. ^d^ACE2 binding affinity (K_d_) measured by MST.

| ID | Structure | PP Assay,<br>IC<sub>50</sub> (μM)<sup>a</sup> | CPE Assay,<br>IC<sub>50</sub> (μM)<sup>b</sup> | ACE2 binding,<br>K<sub>d</sub> (μM)<sup>d</sup> |
| --- | --- | --- | --- | --- |
| 1 |  | 1.15 | 5.01 | 0.43 |
| 2 |  | 2.89 | 6.31 | 1.83 |
| 5 |  | 4.08 | 10 | 16.7 |
| 24 |  | 5.14 | 5.62<sup>c</sup> | 10.6 |
| 25 |  | 5.14 | 10 | 6.55 |
| 19 |  | 10.3 | 11.2<sup>c</sup> | 2.24 |

All six compounds were able to inhibit the PP entry into ACE2-overexpressing HEK293 cells, where the molecule with the strongest affinity to ACE2 showed the highest activity in PP entry inhibition (**Table 2**). Since these compounds do not bind S protein, we hypothesized their activity should be independent of S protein sequence and thus, active against different strains of SARS-CoV-2. Therefore, we tested them against other strains of SARS-CoV-2. As result, we found that compounds inhibited the entry of pseudoparticles bearing S proteins of South African B.1.351 and UK B.1.1.7 variants with the same or greater potency compared to the wild type (**Fig. 4**).

**Fig. 4:**
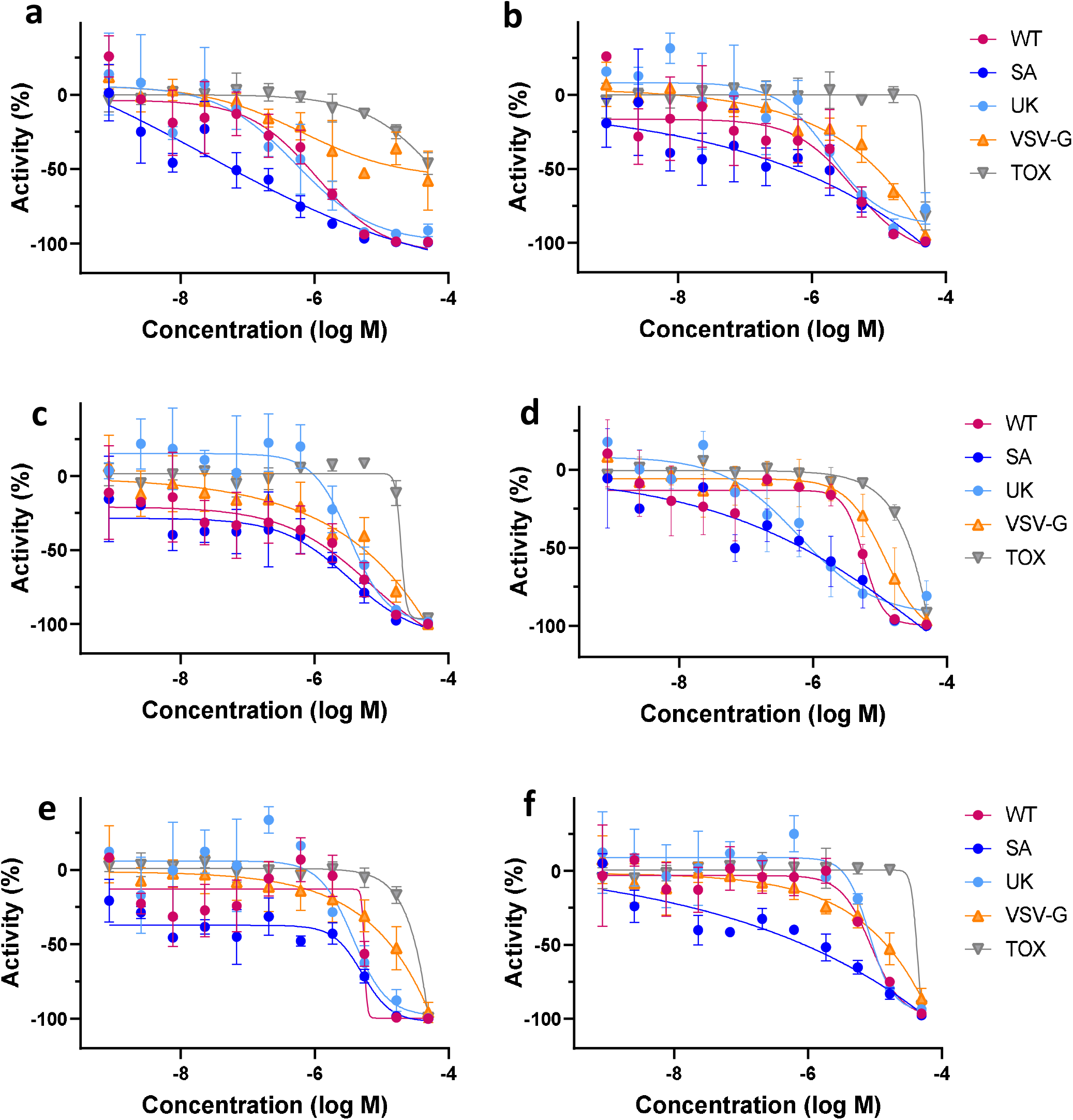
Dose-response curves of the six ACE2 binding compounds in PP and cytotoxicity assays. **a**, compound **1**; **b**, compound **2**; **c**, compound **5**; **d**, compound **24**; **e**, compound **25**; **f**, compound **19**. WT – wild type SARS-CoV-2 variant assay; SA – South African B.1.351 variant assay; UK – UK B.1.1.7 variant assay; VSV-G – PP assay containing the G protein of vesicular stomatitis virus; Tox – cytotoxicity assay.

## Discussion

A traditional QSAR modeling approach relies on the assumption that the biological activity of small molecules is correlated with their physicochemical properties or the so-called structural descriptors^26^, however it does not consider the 3D geometric features of the molecules. This results in an incomplete description of ligand-target interactions. Furthermore, QSAR models are also restricted to their applicability domain, i.e. the chemical space within which the models are originally trained^27^. To overcome these shortcomings, a hybrid approach was developed which combines QSAR models with pharmacophore screening that can retrieve ligands with structurally diverse scaffolds.

Utilization of the hybrid approach led to 4-fold improvement of the hit-rate and revealed multiple novel scaffolds with activity against SARS-CoV-2. More importantly, 44 compounds experimentally confirmed as active in the CPE reduction assay did not show appreciable cytotoxicity.

Further analysis of active analogs revealed some preliminary SAR, although trends were limited due to significant structural differences within the set of analogs. This supports the hypothesis that these compounds are acting on a common target or *via* a shared mechanism to inhibit viral proliferation, thus decreasing the viral cytopathic effect. Overall, the chemotypes identified showed good efficacy and potency as screening hits. These preliminary screening results clearly warrant further investigation of each chemotype *via* medicinal chemistry to thoroughly explore and establish the SAR, as well as optimize physicochemical properties.

In an effort to elucidate the mechanism of action, active compounds were screened against some previously established SARS-CoV-2 targets which have been shown to mediate antiviral activity: SARS-CoV-2 M^pro^ and RBD-ACE2 protein-protein interaction. None of the identified hits exhibited activity against these targets.

Interestingly, we identified six CPE-active compounds which are ACE2 binders and inhibitors of viral entry. We assume these molecules are allosteric binders to ACE2, as they did not inhibit ACE2 enzymatic activity, meaning they did not at least interfere with the substrate binding. Although these compounds do not interrupt the RDB-ACE2 interaction, they were able to inhibit the entry of pseudoparticles resembling wild-type as well as other variants of SARS-CoV-2. These compounds might interfere with the conformational change of S protein bound to ACE2 and/or influence the endosome environment, such as the pH decrease in the endosomal lumen, which triggers the conformational change of S protein. The compounds showed some reduced inhibitory activity, compared to SARS-CoV-2 PP, in the counterscreen experiments with PP containing the G protein of vesicular stomatitis virus (VSV-G). VSV-G do not utilize ACE2 for host cell binding but require low endosomal pH for a conformational change to induce membrane fusion. ACE2-overexpressing HEK293 cells were used for all PP assays, consequently these compounds could bind to ACE2, trap within the endosomes and affect the VSV-G PP entry. Further experimentation is required to determine the exact mechanism by which these compounds hinder viral entry.

To accelerate further research on finding of small molecules active against SARS-CoV-2 we provided the best developed prediction models and modeling sets via github (https://github.com/ncats/covid19_pred) and through the NCATS Predictor website (https://predictor.ncats.io/).

## Supplementary Information

Supplementary Table 1-8

Supplementary Pharmacophore models

## Acknowledgement

This research was supported in part by the Intramural Research Program of the National Center for Advancing Translational Sciences (NCATS), National Institutes of Health (NIH).

