## Supplementary material for "A hybrid *in silico* approach reveals novel inhibitors of multiple SARS-CoV-2 variants": Methods

The data used in this study was obtained from the single agent screening by SARS-CoV-2 cytopathic effect (CPE) and its corresponding counter SARS-CoV-2 cytopathic effect (host tox counterscreen) assays available at the NCATS OpenData Portal (https://opendata.ncats.nih.gov/covid19/). As NCATS has been unceasingly performing screening campaigns, we combined additional *in-house* quantitative HTS data to this, which resulted in a dataset of 9,046 compounds.

**Dataset Curation**

The initial data set was curated in the following steps: (i) removal of inorganic compounds according to the chemical formula in MOE 2014.09.31; (ii) removal of salts and compounds containing metals and/or rare or special atoms; (iii) standardization of chemical structures using Francis Atkinson’s standardiser (https://github.com/flatkinson/standardiser) (iv) removal of duplicates and permanently charged compounds using MOE 2014.09.31.

**Compound Labeling**

Compounds having an IC_50_ < 30 μM, curve class in the range of 1 to 3^28^ and a maximum response (MaxResponse) > 30 % were considered active for the CPE reduction assay, whereas others were labeled as inactive. For the cytotoxicity counter-screen, compounds with an IC_50_ < 30 μM, curve class in the range of 1 to -3, and MaxResponse < -30 % were considered active and others as inactive. In the combined data set, compounds active in the CPE reduction assay and inactive in the counter-screen assay were considered as active. All others were labeled as inactive. While merging the data from multiple protocols, the compound with contradictory results in different experimental runs was removed from the study. In the 1^st^ round of modeling, the data set was comprised of 8474 compounds (319 active and 8155 inactive). Enriching this dataset with compounds identified in the 1^st^ round of virtual screening and experimentally tested in the 1^st^ round of screening resulted in a dataset of 9046 compounds (456 active and 8590 inactive).

### **Descriptor Calculation**

Three different sets of descriptors were calculated for all datasets using RDKit (https://www.rdkit.org/).

1. RDKit descriptors based on the two-dimensional structure (119 descriptors in total).

2. Morgan fingerprints (1024 bits).

3. Avalon fingerprints (1024 bits).

### **Training and Test Set Selection**

From each class (active, inactive), 70 % of the data was randomly selected and used as a training set. The remaining 30 % of compounds were considered as the test set. Five-fold external cross-validation was omitted since pharmacophore modeling is computationally expensive and the selection of the best consensus model combining pharmacophore and machine learning approaches requires the established training and the test set. We emphasize that the selected consensus model was used in virtual screening and thus prospectively validated. The composition of the resulting datasets is shown in **Table 3**.

|  | **Total Compounds** | **Active** | **Inactive** | **Imbalance Ratio (inactive/active)** |
| --- | --- | --- | --- | --- |
| **1^st^ Round** |  |  |  |  |
| Full dataset | 8474 | 319 | 8155 | 26:1 |
| Training set | 5931 | 223 | 5708 | 26:1 |
| Test set | 2543 | 96 | 2447 | 25:1 |
| **2^nd^ Round** |  |  |  |  |
| Full dataset | 9046 | 456 | 8590 | 19:1 |
| Training set | 6332 | 319 | 6013 | 19:1 |
| Test set | 2714 | 137 | 2577 | 19:1 |

**Table 3**. An overview of the datasets used in this study.

**Virtual Screening Libraries**

To discover compounds active against SARS-CoV-2, we performed virtual screening using two of our internal libraries (approximately 140K compounds). These libraries contain a diverse collection of small molecules with an emphasis on medicinal chemistry-tractable scaffolds. The compound libraries were curated using the same protocol described in the *Dataset* [*Curation*](https://pubs.acs.org/doi/10.1021/acs.molpharmaceut.5b00594#sec2_1) section. These compounds were screened against the model and rank-ordered based on the predicted activity score, which roughly corresponds to their probability of being active against SARS-CoV-2.

**Machine Learning: Stratified Bagging**

Considering the high degree of data imbalance (1:26), we used under-sampling stratified bagging (SB). This method has been proven to be the best performing method for dealing with imbalanced datasets^29^ . SB is a machine learning technique based on an ensemble of models developed using multiple training datasets sampled from the original training set. It uses a traditional bagging approach (resampling with replacement) to create the training set of positive samples and randomly selects the same number of samples from the majority class. Thus, the total bagging training set size is double the minority class. Several models are then built, and predictions are averaged to produce a final ensemble model output. Because of random sampling, about 37% of the compounds are left out in each run. These samples form "out-of-the-bag" sets, which are used to test the final model. Although a small set of samples are selected each time, most compounds contribute to the overall bagging procedure since datasets were generated randomly. Random Forest (RF) was used as a base-classifier^30^. The number of trees was arbitrarily set to 100 (default) since it has been shown that the optimal number of trees is usually 64-128, while further increasing the number of trees does not necessarily improve the model's performance^31^.

As consensus QSAR modeling is another highly recommended approach that has been reported to outperform simple QSAR models^32,33^. In this study, we considered a consensus approach that takes the consensus of the predictions from the three different descriptors to predict anti-SARS-CoV-2 activity.

**Pharmacophore-based Screening**

A pharmacophore describes the spatial arrangement of essential interactions of a drug with its respective receptor binding site. Pharmacophore modeling and subsequent virtual screening (VS) is a well-established method in the early drug discovery process^21,22^. In this study, the generation of ligand-based pharmacophore models, their subsequent refinement, and virtual screening (VS) were performed with LigandScout 4.4 Advanced (Inte:Ligand GmbH). The conformational libraries for both pharmacophore modeling and the VS process were created with i:Con (max. 200 conformations per compound), a conformer generator implemented in LigandScout^34^. To design the ligand-based pharmacophore model, the most potent compounds were selected based on the IC_50_ (< 30 μM) and MaxResponse (> 50.0 %) values. The molecules were clustered based on pharmacophore-based similarity (cluster distance 0.4, 0.6, 0.7, and 0.8, respectively). For each of the clusters obtained from different cluster distance thresholds, merged-features pharmacophore (MFP) and shared-features pharmacophore (SFP) models were generated that incorporate the features of selected compounds per cluster. A good pharmacophore model should not only be able to estimate the activity of active compounds, but also have the ability to identify the active molecules from a database containing a large number of inactive compounds. To select the best models for screening, we applied these models on our complete dataset (training and test set combined) and calculated the percentage of active and inactive that hit these pharmacophore models. The models that hit 20% more active compounds versus inactive compounds were selected for the final virtual screening. The screening was performed using iscreen module, with default settings with the maximum number of omitted features set to 2.

**1st Round of *in silico* Screening**

The complete collection of 138,749 compounds was tested against our stratified bagging models and ranked by the prediction score. This score takes a value between 0 and 1, and the higher the score, the higher the probability of the compound to be active against SARS-CoV-2. In the 1^st^ round of screening, we selected the top 300 predictions from each descriptor combination, thus identified a subset of 890 compounds that were predicted to be in the top 300 by at least two out of the four models. We then retrieved all generated conformations of these 890 compounds prepared from the i:Con^34^ and screened them against our pharmacophore models (LBP-1) (Supplementary Table 2). Finally, we ranked the compounds according to their pharmacophore fit score. The top 320 compounds were selected for experimental validation in the SARS-CoV-2 CPE assay and cytotoxicity counter-screen assay.

**2^nd^ Round of *In Silico* Screening**

After obtaining the results from the 1^st^ round of modeling, followed by the experimental validation, we updated our machine learning (SB) and ligand-based pharmacophore (LBP) model with the new (confirmed) SARS-CoV-2 active and inactive compounds. We then removed the 320 compounds tested in the 1st round of experimental validation from our virtual screening library. The remaining compounds (138,429 compounds) were predicted using our updated stratified bagging models (SB-2) and ranked by the prediction score. In this 2nd round of screening, we selected the top 500 predictions from each descriptor combination. This resulted in a subset of 1325 compounds that were consistently present in the list of top predictions by at least two out of the four models. We screened the 138,429 compounds using the updated ligand-based pharmacophore models (LBP-2) shortlisted for the screening (Supplementary Table 3). This gave us 65,952 compounds. We then ranked these compounds according to their pharmacophore fit and selected 320 compounds that were overlapping with the 1325 compounds for the experimental validation.

**Model Performance Assessment**

Receiver Operating Characteristic Area under the curve (ROC AUC) was used to assess the performance of the models. ROC AUC plots sensitivity (TP/(TP+FN)) against 1-specificity (TN/(TN+FP)). The higher the ROC AUC value, the better the model performs in distinguishing between active and inactive compounds. A ROC AUC value of 1.0 indicates a perfect classification model, whereas a value close to 0.5 indicates that the model provides random predictions. The ROC AUC score was calculated using the ROC curve (javascript) node in KNIME^35^. For the experimental validation results, the model performance was measured by the positive predicted value (PPV=TP/(TP+FP)). Based on the earlier studies, we chose PPV and AUC as our model performance metrics^36,37^.

$$Sensitivity=\frac{TP}{(TP+FN)}$$

$$Specificity=\frac{TN}{(TN+FP)}$$

$$Balanced Accuracy=\frac{1}{2}\left( \frac{(TP)}{(TP+FN)}+\frac{(TN)}{(TN+FP)} \right)$$

$$MCC=\frac{\left\{ \left( TP \times TN \right)-(FP\times FN) \right\}}{\left\{ (TP+FP)\times(TP+FN)\times(TN+FP)\times(TN+FN) \right\}^{1/2}}$$

$$Precicion=\frac{TP}{(TP+FP)}$$

TP: true positives; TN: true negatives; FP: false positives; FN: false negatives;

MCC: Matthews correlation coefficient

**Experimental Testing**

**SARS-CoV-2 Cytopathic Effect Screening**

The cytophatic effect (CPE) of SARS-CoV-2 was measured in Vero E6 cells in a BSL-3 facility as described previously^10^. Cells were harvested, resuspended at 160,000 cells/mL in assay media (MEM, 2% (v/v) heat inactivated FBS, 1% HEPES, 1% Pen/Strep/GlutaMax) and inoculated with SARS-CoV-2 (USA_WA1/2020) at a multiplicity of infection (MOI) of 0.002. 25 µL of cell-virus mixture were dispensed per well of an assay ready 384-well-plate (Greiner, # 781091). The assay ready plates were prepared by adding 5 µL of assay media per well pre-spotted with 60 nL of library compounds at a 5-point serial dilution with concentrations ranging from 10 mM to 62 µM (final assay concentrations ranging from 20 µM to 124 nM) using an acoustic dispenser (Echo550, Labcyte, Inc.). Each plate contained 2 columns with 60 nL DMSO as negative (no inhibitor) control, and 24 wells of cells only (no virus) as positive control. Plates were incubated for 72 h at 37°C, 5% CO2, 90% humidity. The cell viability was assessed by measuring the luminescence signal with Envision plate reader (Perkin Elmer) after addition of 30 µL/well of CellTiter-Glo reagent (Promega, Cat # G7573) and 10 min incubation at room temperature. The signal was normalized against negative (0% response) and positive control (100% response) and the resulting percent of inhibition data were fitted to a sigmoidal dose response curve using four-parameter Hill equation.

**Cytotoxicity Counter-Assay**

The assay was set up in the same way as in the CPE assay, but omitting the addition of virus. DMSO and hyamine at 100 µM final concentration served as negative and positive controls, respectively. The obtained luminescence signal was normalized against negative control (0% response) and positive control (-100% response).

**SARS-CoV-2 M^pro^ Assay**

The ability of the compounds to inhibit the recombinant M^pro^ activity was measured by a biochemical assay, SARS-CoV-2 M^pro^ assay described previously^38^.

**ACE2-RBD AlphaLISA Proximity Assay**

The interaction of SARS-CoV-2 receptor binding domain (RBD) with ACE2 was tested using an AlphaLISA proximity assay in combination with a corresponding TruHits counter-assay as described elsewhere^39^.

**Microscale Thermophoresis Assay**

Binding of the compounds to recombinant human ACE2 (Sino Biological, Cat #: 10108-H08H) was evaluated by microscale thermophoresis (MST). His-tagged ACE2 was labeled with RED-tris-NTA 2^nd^ generation dye (Nanotemper Technologies, Cat #: MO-L018) following manufacturer’s protocol and diluted in MST buffer (10 mM HEPES pH 7.4, 150 mM NaCl, 10 mM CaCl2, 0.01% Tween 20) to a final concentration of 3 nM. 100 nL of compounds in two-fold dilution series were transferred to 384-well compound plate (Greiner, Cat #: 784201-1B) using Echo 650 series acoustic dispenser (Labcyte Inc.), mixed with 10 μL of labeled protein and incubated for 15 min at RT. MST traces were collected using Monolith NT.Automated (Nanotemper Technologies) unit and standard treated capillary chip (Nanotemper Technologies, Cat #: MO AK002) with following setting: 45% excitation power, medium MST power and MST periods of 3s/10s/1s. K_d_ values were calculated by fitting the change in normalized fluorescence signal of the thermograph using MO.Affinity analysis software.

**ACE2 Enzymatic Assay**

ACE2 enzyme activity was monitored in a fluorometric assay. Briefly, 25 nL compounds were transferred to the 1536-well assay plate (Greiner, solid black medium-binding plates) using Echo 650 (Labcyte Inc.) acoustic dispenser. 3μL/well of 0.27 nM ACE2 (0.2 nM final concentration) suspension in assay buffer (PBS, pH 7.4, 0.01% Tween-20) was dispensed into assay plate with Aurora Discovery BioRAPTR Dispenser (FRD; Beckton Dickenson) and incubated 15 min at room temperature (RT). One μL/well of 60 μM ACE2 substrate (AnaSpec, Cat #: AS-60757) was then added. The plate was centrifuged at 1000 rpm for 15 sec and the fluorescence was detected with the PHERAstar plate reader (BMG Labtech) equipped with Module 340/440 at t1=0 min and t2=15 min at RT. Data was normalized to enzyme activity in presence of DMSO, set as 0%, and in presence of 6.2 µM MLN-4760, set as -100% inhibition. The resulting percent of inhibition data were fitted to a sigmoidal dose response curve using four-parameter Hill equation.

**Pseudotyped Particle (PP) Entry Assay**

Expi293F cells with stable expression of human ACE2 (HEK293-ACE2, Codex Biosolutions, Cat #: CB-97100-220) were seeded in white, solid bottom 384-well microplates (Greiner BioOne) at 6000 cells/well in 30 µL/well medium (DMEM, 10% FBS, 1x L-glutamine, 1x Pen/Strep, 1 ug/ml puromycin), and incubated at 37 °C with 5% CO2 overnight (~16 h). 150 nL of compounds at 11-point titration, 1:3 dilution in DMSO were dispensed via Echo 650 (Labcyte Inc.) acoustic dispenser to assay plates. Cells were incubated with the test compounds for 1 h at 37 °C with 5% CO2, before 2 µL/well of SARS-CoV-2-S PPs were added. PPs with following spike variants were used: wild-type (Codex Biosolutions, Cat #: CB-97100-154), South African variant B.1.351 (Codex Biosolutions, Cat #: CB-97100-154) and U.K. variant B.1.1.7 (Codex Biosolutions, Cat #: CB-97100-153). The plates were spinoculated by centrifugation at 1500 rpm (453 x g) for 45 min at RT and incubated for 48 h at 37 °C with 5% CO2 to allow cell entry of PPs and expression of luciferase reporter. After the incubation, the supernatant was removed with gentle centrifugation using a Blue Washer (BlueCat Bio). Four µL/well of Bright-Glo Luciferase detection reagent (Promega) was added to assay plates and incubated for 5 min at RT. The luminescence signal was measured using a PHERAStar plate reader (BMG Labtech). Data was normalized with wells inoculated with SARS-CoV-2-S PPs as 100%, and wells inoculated with bald PPs (no spike protein) as 0%.

An ATP content cytotoxicity assay was done as a counter-assay. HEK293-ACE2 cells were seeded in white, solid bottom 384-well microplates (Greiner BioOne) and treated with compounds under the same experimental conditions as in the PP entry assay omitting the inoculation step. After 48 h incubation, 4 µL/well of ATPLite (PerkinElmer) was added to assay plates and incubated for 15 min at RT. The luminescence signal was measured using a Viewlux plate reader (PerkinElmer). Data was normalized with wells containing cells as 100%, and wells containing media only as 0%.
