## Supplementary material for "A hybrid *in silico* approach reveals novel inhibitors of multiple SARS-CoV-2 variants": SI

**Supplementary Table 1.** Model performance on the training and test dataset.

| Training set performance | | | | | | | |
| --- | --- | --- | --- | --- | --- | --- | --- |
| **Model** | **Descriptor** | **Sensitivity** | **Specificity** | **Balanced Accuracy** | **MCC** | **Precision** | **AUC** |
| **SB-1** | RDKit | 0.66 | 0.77 | 0.72 | 0.19 | 0.10 | 0.79 |
|  | Morgan | 0.47 | 0.85 | 0.66 | 0.17 | 0.11 | 0.81 |
|  | Avalon | 0.48 | 0.83 | 0.66 | 0.16 | 0.10 | 0.76 |
|  | Consensus | 0.58 | 0.84 | 0.71 | 0.20 | 0.12 | 0.80 |
| **SB-2** | RDKit | 0.72 | 0.81 | 0.77 | 0.28 | 0.17 | 0.84 |
|  | Morgan | 0.60 | 0.84 | 0.72 | 0.25 | 0.17 | 0.82 |
|  | Avalon | 0.57 | 0.84 | 0.70 | 0.23 | 0.16 | 0.81 |
|  | Consensus | 0.66 | 0.84 | 0.75 | 0.28 | 0.18 | 0.85 |
| Test set performance | | | | | | | |
| **Model** | **Descriptor** | **Sensitivity** | **Specificity** | **Balanced Accuracy** | **MCC** | **Precision** | **AUC** |
| **SB-1** | RDKit | 0.65 | 0.78 | 0.71 | 0.19 | 0.10 | 0.80 |
|  | Morgan | 0.54 | 0.85 | 0.70 | 0.21 | 0.13 | 0.80 |
|  | Avalon | 0.48 | 0.83 | 0.65 | 0.15 | 0.10 | 0.75 |
|  | Consensus | 0.65 | 0.81 | 0.73 | 0.22 | 0.12 | 0.80 |
| **SB-2** | RDKit | 0.65 | 0.82 | 0.73 | 0.25 | 0.16 | 0.82 |
|  | Morgan | 0.61 | 0.83 | 0.72 | 0.25 | 0.16 | 0.79 |
|  | Avalon | 0.58 | 0.85 | 0.71 | 0.24 | 0.17 | 0.80 |
|  | Consensus | 0.62 | 0.84 | 0.73 | 0.26 | 0.17 | 0.84 |

**Supplementary Table 2.** Ligand-based pharmacophore models generated for the first round of screening (LBP-1).

| Pharmacophore | pharmacophore type | cluster distance |
| --- | --- | --- |
| AR-HBA-HBA-HBA-HBA-HBD-HBD-HBD-XV33 | MFP | 0.4 |
| HBA-HBA-HBA-HBA-XV36 | MFP | 0.6 |
| AR-AR-H-HBA-HBA-HBD-PI-XV28 | MFP | 0.6 |
| H-H-HBA-HBD-XV44 | MFP | 0.6 |
| H-HBA-HBA-HBA-HBD-HBD-XV40 | MFP | 0.6 |
| H-H-HBA-HBA-HBA-XV48 | MFP | 0.6 |
| H-HBA-HBA-HBD-XV25 | MFP | 0.7 |
| H-HBA-HBA-HBA-HBA-HBD-XV42 | MFP | 0.7 |
| H-H-HBA-HBA-HBA-HBA-HBD-HBD-HBD-XV45 | MFP | 0.7 |
| H-HBA-HBA-XV29 | MFP | 0.7 |
| H-H-HBA-XV25 | MFP | 0.7 |
| AR-H-HBA-HBD-XV23 | MFP | 0.7 |
| H-HBA-HBA-XV17 | MFP | 0.7 |
| H-HBA-HBA-HBA-HBD-HBD-HBD-XV34 | MFP | 0.8 |
| H-H-HBA-XV29 | MFP | 0.8 |
| AR-H-HBA-HBA-HBA-HBD-XV45 | MFP | 0.8 |
| H-H-HBA-HBA-XV30 | MFP | 0.8 |
| H-H-HBA-HBD-XV32 | MFP | 0.8 |
| H-HBA-HBD-XV30 | MFP | 0.8 |
| H-HBA-HBA-HBA-XV13 | MFP | 0.8 |
| AR-H-HBA-HBD-XV17 | MFP | 0.8 |
| H-HBA-HBA-XV21 | MFP | 0.8 |
| AR-H-HBA-HBA-HBD-XV36 | SFP | 0.4 |
| H-HBA-HBA-XV35 | SFP | 0.6 |
| H-HBA-HBD-XV28 | SFP | 0.6 |
| H-H-HBA-HBA-XV40 | SFP | 0.6 |
| AR-H-HBA-HBD-XV28 | SFP | 0.6 |
| AR-H-H-HBA-XV29 | SFP | 0.6 |
| AR-H-HBD-XV24 | SFP | 0.7 |
| AR-H-HBA-XV21 | SFP | 0.7 |
| H-H-HBA-HBD-XV38 | SFP | 0.7 |
| H-HBA-HBD-XV27 | SFP | 0.7 |
| H-H-HBA-XV33 | SFP | 0.7 |
| H-HBA-HBD-XV34 | SFP | 0.7 |
| H-HBA-HBA-XV28 | SFP | 0.7 |
| HBA-HBA-HBA-XV25 | SFP | 0.8 |
| H-H-HBA-XV31 | SFP | 0.8 |
| AR-AR-H-HBA-HBA-XV32 | SFP | 0.8 |
| H-H-HBA-XV30 | SFP | 0.8 |
| H-H-HBA-XV27 | SFP | 0.8 |
| H-HBA-HBA-XV38 | SFP | 0.8 |
| H-H-HBA-XV24 | SFP | 0.8 |
| H-HBA-HBD-XV27 | SFP | 0.8 |
| H-H-HBA-XV17 | SFP | 0.8 |

| H | Hydrophobic |
| --- | --- |
| HBA | H-Bond Acceptor |
| HBD | H-Bond Donor |
| PI | Positive Ionizable |
| NI | Negative Ionizable |
| AR | Aromatic Ring |
| XV | Exclusion Volume |
|  | pharmacophore models that hit the majority (>20%) of active vs. inactive and selected for virtual screening |

**Supplementary Table 3.** Ligand-based pharmacophore models generated for the second round of screening (LBP-2).

| Pharmacophore | pharmacophore type | cluster distance |
| --- | --- | --- |
| HBA-HBA-HBA-HBA-HBD-XV36 | MFP | 0.4 |
| AR-AR-AR-H-H-H-HBA-HBA-HBD-HBD-HBD-PI-XV48 | MFP | 0.4 |
| AR-AR-AR-H-H-H-H-HBA-HBA-HBD-HBD-HBD-PI-XBD-XV50 | MFP | 0.4 |
| H-H-HBA-HBA-HBD-HBD-HBD-HBD-XV36 | MFP | 0.4 |
| AR-H-H-HBA-HBA-HBA-HBD-XBD-XV31 | MFP | 0.4 |
| HBA-HBA-HBA-HBD-XV38 | MFP | 0.6 |
| AR-AR-AR-H-H-H-HBA-HBA-HBD-HBD-PI-XV50 | MFP | 0.6 |
| AR-AR-H-HBA-HBA-HBA-HBA-HBD-HBD-XV45 | MFP | 0.6 |
| AR-AR-AR-H-H-H-HBA-HBA-HBD-PI-XV42 | MFP | 0.6 |
| H-H-HBA-HBD-HBD-HBD-XV26 | MFP | 0.6 |
| AR-AR-AR-H-H-H-H-H-HBA-HBA-HBD-HBD-HBD-PI-XBD-XV51 | MFP | 0.6 |
| H-H-HBA-HBA-HBA-HBD-HBD-XV42 | MFP | 0.6 |
| AR-H-HBA-HBA-HBA-HBD-HBD-XV42 | MFP | 0.6 |
| AR-AR-AR-H-H-H-HBA-HBA-HBD-HBD-PI-XV50 | MFP | 0.7 |
| H-HBA-HBA-HBD-HBD-HBD-XV28 | MFP | 0.7 |
| AR-AR-AR-H-H-H-HBA-HBD-HBD-XBD-XV44 | MFP | 0.7 |
| AR-AR-H-H-HBA-HBA-HBA-HBD-PI-XBD-XV36 | MFP | 0.7 |
| H-HBA-HBD-HBD-XV26 | MFP | 0.7 |
| AR-H-H-PI-XV24 | MFP | 0.7 |
| H-H-H-H-HBA-HBA-PI-XV51 | MFP | 0.7 |
| H-HBA-HBA-HBD-HBD-XV28 | MFP | 0.7 |
| AR-H-H-HBA-HBA-HBD-PI-XBD-XV43 | MFP | 0.8 |
| HBA-HBA-HBD-XV12 | MFP | 0.8 |
| AR-AR-H-HBA-HBA-HBD-XV39 | MFP | 0.8 |
| H-HBA-HBD-XV15 | MFP | 0.8 |
| AR-H-HBA-HBA-XV33 | SFP | 0.4 |
| AR-AR-H-H-HBA-HBA-HBD-PI-XV43 | SFP | 0.4 |
| AR-AR-AR-H-H-H-H-HBA-HBD-HBD-XBD-XV45 | SFP | 0.4 |
| H-H-H-HBA-HBA-HBA-HBD-XV50 | SFP | 0.4 |
| AR-H-H-HBA-HBA-XV31 | SFP | 0.4 |
| AR-H-HBA-HBA-XV34 | SFP | 0.6 |
| AR-AR-H-H-H-HBA-HBA-HBD-PI-XV47 | SFP | 0.6 |
| AR-H-HBA-HBA-HBA-HBD-XV47 | SFP | 0.6 |
| AR-AR-H-HBA-HBA-HBD-PI-XV29 | SFP | 0.6 |
| H-HBA-HBA-HBD-XV23 | SFP | 0.6 |
| AR-AR-H-H-H-H-HBA-HBA-HBD-PI-XBD-XV45 | SFP | 0.6 |
| H-H-HBA-HBD-XV36 | SFP | 0.6 |
| H-HBA-HBA-HBD-XV25 | SFP | 0.6 |
| AR-H-HBD-XV28 | SFP | 0.6 |
| H-HBA-HBA-HBD-HBD-XV32 | SFP | 0.6 |
| AR-AR-H-H-H-HBA-HBA-HBD-PI-XV49 | SFP | 0.7 |
| H-HBA-HBD-XV22 | SFP | 0.7 |
| AR-H-HBA-HBA-XV38 | SFP | 0.7 |
| AR-AR-H-H-H-HBA-HBD-XV34 | SFP | 0.7 |
| AR-AR-H-HBA-HBA-HBD-PI-XV26 | SFP | 0.7 |
| H-HBD-HBD-XV23 | SFP | 0.7 |
| AR-H-H-HBA-HBD-XV27 | SFP | 0.7 |
| H-H-PI-XV31 | SFP | 0.7 |
| AR-H-H-HBA-XV38 | SFP | 0.7 |
| AR-H-H-HBA-PI-XBD-XV28 | SFP | 0.8 |
| H-HBA-HBA-HBD-XV14 | SFP | 0.8 |
| AR-HBA-HBA-XV13 | SFP | 0.8 |
| H-HBA-PI-XV32 | SFP | 0.8 |
| H-H-HBD-XV30 | SFP | 0.8 |
| H-HBA-HBD-XV20 | SFP | 0.8 |

| H | Hydrophobic |
| --- | --- |
| HBA | H-Bond Acceptor |
| HBD | H-Bond Donor |
| PI | Positive Ionizable |
| NI | Negative Ionizable |
| AR | Aromatic Ring |
| XV | Exclusion Volume |
|  | pharmacophore models that hit the majority (>20%) of active vs. inactive and selected for virtual screening |

**Supplementary Table 4.** Experimental results for the SARS-CoV-2 actives from the 1st round of testing in CPE and cytotoxicity counterscreen assay (5-point dilution series).

| **Sample ID** | **Smiles** | **AC50 (uM)** | **CC-v2** | **Efficacy** | **Max response** | **AC50 (uM)_counter-screen** | **CC-v2_counter-screen** | **Efficacy_counter-screen** | **Max response_counter-screen** |
| --- | --- | --- | --- | --- | --- | --- | --- | --- | --- |
| NCGC00127046-01 | CCOc1ccccc1CNCc1cccn1-c1nnc(N2CCN(c3ccccc3)CC2)s1 | 3.162278 | 1.2 | 42.383 | 40.209 | 14.125375 | 4 | -25.051 | -22.959 |
| NCGC00104471-02 | CCN(CC)CCNC(=O)c1cc(-c2ccc(Br)s2)nc2ccccc12 | 3.548134 | 1.3 | 84.477 | 71.952 | 14.125375 | 4 | -15.243 | -14.369 |
| NCGC00140192-01 | CCc1ccc(Sc2cc(C(=O)NCCCN3CCC(C)CC3)c3ccccc3n2)cc1 | 3.548134 | 1.2 | 62.383 | 54.142 | 11.220185 | 4 | -18.552 | -21.906 |
| NCGC00140356-01 | Cc1ccc(-c2nc(C[S+]([O-])CC(=O)N3CCN(c4ccc(Cl)cc4)CC3)c(C)o2)cc1 | 3.548134 | 1.2 | 36.906 | 32.849 | 14.125375 | 4 | -16.907 | -16.995 |
| NCGC00104445-01 | CCCCN(C)CCNC(=O)c1cc(-c2ccc(Cl)s2)nc2ccccc12 | 4.466836 | 1.2 | 76.459 | 75.466 | 14.125375 | 4 | -15.091 | -14.659 |
| NCGC00139668-01 | CC1CCCCN1CCCNC(=O)c1cc(Sc2ccc(Cl)cc2)nc2ccccc12 | 6.309573 | 2.1 | 102.706 | 91.731 | NaN | 4 | 0 | -1.679 |
| NCGC00114893-01 | CCc1ccccc1N1CC(c2nc3ccccc3n2CCCCOc2ccccc2OC)CC1=O | 10 | 3 | 39.31 | 35.544 | 15.848932 | 4 | -10.618 | -10.099 |
| NCGC00510147-03 | CC(C)(C)N1CCC(c2ccccc2)(c2ccccc2)CC1 | 10 | 2.2 | 50.396 | 40.663 | NaN | 4 | 0 | -4.245 |
| NCGC00139670-01 | CCCN(CCC)CCNC(=O)c1cc(Sc2ccc(Cl)cc2)nc2ccccc12 | 10 | 2.1 | 97.171 | 86.854 | NaN | 4 | 0 | -3.062 |
| NCGC00126319-01 | Cc1ccc(Cl)cc1N1CCN(CCCNc2ncnc3onc(-c4ccc(F)cc4)c23)CC1 | 11.220185 | 3 | 72.129 | 65.803 | NaN | 4 | 0 | -1.789 |
| NCGC00140413-01 | Cc1ccccc1CN1C(=O)c2ccccc2Sc2ccc(C(=O)NCCCN3CCOCC3)cc21 | 11.220185 | 3 | 68.651 | 65.622 | NaN | 4 | 0 | -9.958 |
| NCGC00140190-01 | CCc1ccc(Sc2cc(C(=O)NCCCN3CCCC(C)C3)c3ccccc3n2)cc1 | 11.220185 | 2.1 | 102.202 | 91.58 | NaN | 4 | 0 | -1.679 |
| NCGC00377203-01 | COc1ccccc1-c1ccc(CN2CCCC(c3nc4ccc(C)cc4c(=O)[nH]3)C2)cc1 | 11.220185 | 2.1 | 93.787 | 89.187 | 12.589254 | -2.4 | -32.682 | -26.818 |
| NCGC00498981-01 | Cc1ccc(-c2cccc(CN3CCCC(NC(=O)c4c(C)noc4C)C3)c2)cc1Cl | 11.220185 | 2.1 | 101.374 | 92.579 | NaN | 4 | 0 | -2.441 |
| NCGC00099741-01 | Cc1cc(OCC(O)CN2CCN(c3cccc(Cl)c3)CC2)ccc1Cl | 12.589254 | 3 | 88.291 | 80.373 | NaN | 4 | 0 | -6.691 |
| NCGC00102225-01 | COc1ccc(Cl)cc1NC(=O)CSc1nc2ccccc2nc1N1CCCCC1 | 12.589254 | 3 | 100.946 | 92.034 | NaN | 4 | 0 | -1.769 |
| NCGC00109669-01 | Cc1oc(-c2cccc(Cl)c2)nc1CN1CCCC(C(=O)NCCc2ccc(Cl)cc2)C1 | 12.589254 | 3 | 49.891 | 43.844 | 12.589254 | 4 | -18.612 | -18.629 |
| NCGC00116825-01 | Oc1ccccc1-c1nc(-c2cccs2)c(-c2cccs2)[nH]1 | 12.589254 | 3 | 52.719 | 47.993 | 14.125375 | 4 | -16.903 | -16.905 |
| NCGC00116815-01 | CN(C)c1ccc(-c2nc(-c3cccs3)c(-c3cccs3)[nH]2)cc1 | 12.589254 | 3 | 45.173 | 41.118 | NaN | 4 | 0 | -1.007 |
| NCGC00120248-01 | CCC1CCCCN1CCCNC(=O)c1sc2nc(C)cc(C)c2c1-n1cccc1 | 12.589254 | 3 | 102.132 | 90.489 | NaN | 4 | 0 | -2 |
| NCGC00126272-01 | CCOc1ccc(NC(=O)c2cnn(-c3ccc(C)c(Cl)c3)c2C2CCNCC2)cc1 | 12.589254 | 3 | 99.715 | 88.854 | NaN | 4 | 0 | -8.174 |
| NCGC00140311-01 | COc1ccc(CCNC(=O)c2cc(Nc3ccccc3)nc3ccccc23)cc1OC | 12.589254 | 3 | 52.14 | 46.358 | NaN | 4 | 0 | -5.267 |
| NCGC00139690-01 | CC1CCCN(CCCNC(=O)c2cc(Sc3ccc(F)cc3)nc3ccccc23)C1 | 12.589254 | 3 | 109.007 | 97.607 | NaN | 4 | 0 | 4.355 |
| NCGC00140464-01 | Cc1cccc(N2CCN(C(=O)C3CCN(Cc4nc(-c5ccc(Cl)cc5)oc4C)CC3)CC2)c1C | 12.589254 | 3 | 118.512 | 108.39 | NaN | 4 | 0 | -2.26 |
| NCGC00426433-01 | COc1cccc(-c2ccccc2NC(=O)C2CCN(Cc3ccccc3F)CC2)c1 | 12.589254 | 3 | 98.98 | 88.339 | NaN | 4 | 0 | -1.679 |
| NCGC00427056-01 | Cn1cc(CN2CCCC(COc3ccccc3C(=O)N3CCCCC3)C2)c2ccccc21 | 12.589254 | 3 | 92.629 | 83.28 | NaN | 4 | 0 | -5.849 |
| NCGC00490405-01 | Cc1ccc(Nc2ccnc(N3CCCC3C(=O)NCCCc3ccccc3)n2)cc1 | 12.589254 | 3 | 39.935 | 36.09 | NaN | 4 | 0 | -7.733 |
| NCGC00493147-01 | COc1ccccc1-c1ccc(CN2CCC(NC(=O)Cc3ccc(F)cc3)C2)cc1 | 12.589254 | 3 | 98.957 | 90.58 | NaN | 4 | 0 | -0.987 |
| NCGC00498641-01 | O=C(Nc1cccc(Cl)c1)C1CCCN(Cc2cccc(-c3ccc(F)cc3F)c2)C1 | 12.589254 | 3 | 47.265 | 42.117 | NaN | 4 | 0 | -9.457 |
| NCGC00498661-01 | O=C(NCc1ccccc1Cl)C1CCCN(Cc2cccc(-c3ccc(F)cc3F)c2)C1 | 12.589254 | 3 | 52.124 | 47.388 | 14.125375 | 4 | -11.422 | -14.108 |
| NCGC00498682-01 | COc1ccccc1-c1cccc(CN2CCCC(C(=O)NCc3ccc(F)cc3)C2)c1 | 12.589254 | 3 | 70.18 | 62.078 | 14.125375 | 4 | -18.663 | -20.734 |
| NCGC00498749-01 | Cc1cc(C)cc(-c2ccc(CN3CCCC3C(=O)NCc3cccc(F)c3)cc2)c1 | 12.589254 | 3 | 87.111 | 78.283 | 14.125375 | 4 | -11.208 | -12.825 |
| NCGC00498687-01 | COc1ccccc1-c1cccc(CN2CCCC(C(=O)NCCc3ccc(F)cc3)C2)c1 | 12.589254 | 3 | 79.852 | 70.286 | 7.943282 | 4 | -12.016 | -14.379 |
| NCGC00498993-01 | COc1ccccc1C(=O)NC1CCCN(Cc2cccc(-c3ccc(C)c(Cl)c3)c2)C1 | 12.589254 | 3 | 75.346 | 65.531 | 15.848932 | 4 | -11.507 | -12.194 |
| NCGC00103820-01 | CCN1CCN(c2cc(C)c3cc(NC(=O)c4cccc(Cl)c4)ccc3n2)CC1 | 14.125375 | 3 | 107.678 | 94.851 | NaN | 4 | 0 | -0.887 |
| NCGC00117038-01 | Cc1c(Cl)cccc1-c1ccc(CNCCSc2nnnn2-c2ccccc2)o1 | 14.125375 | 3 | 43.041 | 37.574 | NaN | 4 | 0 | -1.629 |
| NCGC00399676-01 | CN(C)c1ccc(C(=O)NC[C@H]2C[C@@H]3CCN2C[C@@H]3c2cc(C3CCCCC3)nn2C)cc1 | 14.125375 | 3 | 99.867 | 87.793 | NaN | 4 | 0 | -5.999 |
| NCGC00440956-01 | Clc1ccccc1C1(c2ccnc(-c3ccccc3)n2)CCNCC1 | 14.125375 | 3 | 109.327 | 96.153 | NaN | 4 | 0 | -4.626 |
| NCGC00498636-01 | O=C(NCc1cccs1)C1CCCN(Cc2cccc(-c3ccc(F)cc3F)c2)C1 | 14.125375 | 3 | 82.732 | 72.74 | NaN | 4 | 0 | -10.44 |
| NCGC00493148-01 | COc1ccc(CC(=O)NC2CCN(Cc3ccc(-c4ccccc4OC)cc3)C2)cc1 | 14.125375 | 3 | 101.769 | 87.975 | NaN | 4 | 0 | -4.856 |
| NCGC00411503-01 | Cc1noc2c(NC3CCOCC3)cc(-c3cccc(C(F)(F)F)c3)nc12 | 15.848932 | 3 | 41.334 | 41.723 | 14.125375 | 4 | -21.682 | -22.949 |
| NCGC00416827-01 | COc1cc2c(cc1OCC1CC1)NC(=O)C21CCN(c2cc(NC3CC3)ncn2)CC1 | 15.848932 | 3 | 71.885 | 76.496 | NaN | 4 | 0 | 0.236 |

|  | testing results in the cytophatic effect (CPE) assay |
| --- | --- |
|  | testing results in the cytotoxicity Counter-Assay |

**Supplementary Table 5.** Experimental results for the SARS-CoV-2 actives from the 2nd round of testing in CPE and cytotoxicity counterscreen assay (5-point dilution series).

| **Sample ID** | **Smiles** | **AC50 (uM)** | **CC-v2** | **Efficacy** | **Max response** | **AC50 (uM)_counter-screen** | **CC-v2_counter-screen** | **Efficacy_counter-screen** | **Max response_counter-screen** |
| --- | --- | --- | --- | --- | --- | --- | --- | --- | --- |
| NCGC00522637-01 | CC(C)(C)NCc1ccc(Nc2ccnc3cc(Cl)ccc23)cc1O | 3.162278 | 1.3 | 85.147 | 75.53 | 11.220185 | 4 | -17.765 | -19.507 |
| NCGC00100643-02 | COc1ccc(Nc2nc(Nc3ccc(OC)cc3OC)nc(N3CCOCC3)n2)c(OC)c1 | 3.162278 | 1.1 | 87.52 | 90.175 | NaN | 4 | 0 | -12.885 |
| NCGC00108321-02 | CCCCN(CCCNC(=O)c1cc(Nc2cccc(SC)c2)nc2ccccc12)Cc1ccccc1 | 3.548134 | 1.4 | 37.903 | 33.126 | 6.309573 | 4 | -10.269 | -8.974 |
| NCGC00110841-01 | Cc1ccc2nc(Cl)c3cc(C(=O)NCCCN4CCC(C)CC4)sc3c2c1 | 3.548134 | 1.2 | 70.716 | 61.322 | 10 | 4 | -14.903 | -16.651 |
| NCGC00110541-02 | CCCN1CCN(CCCNC(=O)c2cc3c(Cl)nc4ccccc4c3s2)CC1 | 3.981072 | 1.2 | 81.995 | 78.853 | 5.623413 | 4 | -17.134 | -15.945 |
| NCGC00109633-01 | CCCN(CCC)CCCNC(=O)C1CCCN(Cc2nc(-c3ccc(Cl)cc3)oc2C)C1 | 3.981072 | 1.1 | 87.273 | 85.062 | 11.220185 | 4 | -18.959 | -15.799 |
| NCGC00496752-01 | Cc1ccc2c(NC(C)c3ccccc3)nc(N3CCNCC3)nc2c1 | 3.981072 | 1.1 | 92.845 | 92.549 | NaN | 4 | 0 | -9.458 |
| NCGC00377240-01 | COc1cccc(-c2ccc(CN3CCCC(c4nc5ccc(C)cc5c(=O)[nH]4)C3)cc2)c1 | 4.466836 | 1.1 | 99.414 | 100.256 | NaN | 4 | 0 | 6.157 |
| NCGC00479266-01 | Cc1ccc([C@@H](C)Nc2nc(N3CCNCC3)nc3ccccc23)cc1 | 4.466836 | 1.1 | 91.278 | 89.883 | NaN | 4 | 0 | -9.362 |
| NCGC00390104-01 | Cc1ccc(C(C)Nc2nc(N3CCNCC3)nc3ccccc23)cc1 | 5.011872 | 1.1 | 90.132 | 89.408 | NaN | 4 | 0 | -3.94 |
| NCGC00419475-01 | CN(C)CCNC(=O)c1ccc2c(c1)C1(CCN(CC3CC3)C1)CN2Cc1ccccc1 | 5.011872 | 1.1 | 96.941 | 94.302 | NaN | 4 | 0 | -3.398 |
| NCGC00108196-02 | COc1ccc(Nc2cc(C(=O)NCCCN3CCN(c4ccccc4F)CC3)c3ccccc3n2)c(OC)c1 | 5.623413 | 2.2 | 40.356 | 42.768 | 12.589254 | 4 | -31.476 | -29.566 |
| NCGC00487028-01 | CC(Nc1nc(N2CCN(CCN)CC2)nc2ccccc12)c1ccccc1 | 5.623413 | 2.2 | 52.154 | 56.465 | 14.125375 | 4 | -12.702 | -12.372 |
| NCGC00017063-12 | CCN(CC)Cc1cc(Nc2ccnc3cc(Cl)ccc23)ccc1O | 5.623413 | 2.1 | 83.412 | 89.007 | 11.220185 | 4 | -10.768 | -11.211 |
| NCGC00105953-02 | CCCCN(CCCC)CCCNC(=O)c1cc(-c2ccc(Cl)s2)nc2ccccc12 | 6.309573 | 2.2 | 81.257 | 72.462 | NaN | 4 | 0 | -10.165 |
| NCGC00108262-01 | Cc1ccc(C)c(Nc2cc(C(=O)NCCCN(C)C3CCCCC3)c3ccccc3n2)c1 | 6.309573 | 2.2 | 83.393 | 74.58 | 12.589254 | 4 | -29.828 | -27.794 |
| NCGC00492858-01 | Cc1ccc(Nc2ccnc(N3CCC(C(=O)NCCCc4ccccc4)CC3)n2)cc1 | 7.079458 | 2.2 | 53.961 | 49.963 | 11.220185 | -2.4 | -30.352 | -29.111 |
| NCGC00109487-02 | COc1ccc(-c2nc(CN3CCC(C(=O)NCCCN4CCN(c5ccccc5F)CC4)CC3)c(C)o2)cc1OC | 8.912509 | 2.2 | 79.446 | 73.667 | 11.220185 | 4 | -24.31 | -22.76 |
| NCGC00178968-01 | CCN(CC)Cc1ccc(Nc2ccnc3cc(Cl)ccc23)cc1O | 8.912509 | 2.1 | 97.152 | 92.221 | NaN | 4 | 0 | -9.139 |
| NCGC00389098-01 | CC(Nc1nc(N2CCNCC2C)nc2ccccc12)c1ccccc1 | 8.912509 | 2.1 | 112.168 | 98.393 | NaN | 4 | 0 | -9.081 |
| NCGC00113433-01 | CCCCN(C)CCCNC(=O)c1ccc2nc(N3CCCCC3)sc2c1 | 10 | 3 | 69.91 | 66.472 | NaN | 4 | 0 | -8.877 |
| NCGC00105925-01 | CCCN(CCC)CCNC(=O)c1cc(-c2ccc(Br)s2)nc2ccccc12 | 10 | 2.2 | 77.801 | 67.093 | NaN | 4 | 0 | -5.383 |
| NCGC00108138-01 | CCCCN(CCCNC(=O)c1cc(Nc2ccc(OC)c(OC)c2)nc2ccccc12)Cc1ccccc1 | 10 | 2.2 | 69.245 | 60.044 | NaN | 4 | 0 | -5.237 |
| NCGC00140152-01 | CC1CCCCN1CCCNC(=O)c1cc(Sc2cccc(Cl)c2)nc2ccccc12 | 10 | 2.1 | 108.557 | 97.991 | 12.589254 | 4 | -11.597 | -11.782 |
| NCGC00384291-01 | CNCCN(C)c1nc(NC(C)c2ccccc2)c2ccccc2n1 | 10 | 2.1 | 116.742 | 101.899 | NaN | 4 | 0 | -1.927 |
| NCGC00377241-01 | Cc1ccc(-c2cccc(CN3CCCC(c4nc5ccc(C)cc5c(=O)[nH]4)C3)c2)c(C)c1 | 10 | 2.1 | 111.277 | 100.913 | NaN | 4 | 0 | -8.471 |
| NCGC00479261-01 | C[C@@H](Nc1nc(N2CCNCC2)nc2ccccc12)c1ccc(F)cc1 | 10 | 2.1 | 93.718 | 87.217 | 3.981072 | 4 | -8.942 | -9.952 |
| NCGC00099289-01 | CCOc1ccc(CN2CCN(Cc3ccc4c(c3)c3ccccc3n4CC)CC2)cc1 | 11.220185 | 3 | 68.7 | 65.778 | 14.125375 | 4 | -25.834 | -23.612 |
| NCGC00113431-01 | CCCCN(CCCC)CCCNC(=O)c1ccc2nc(N3CCCCC3)sc2c1 | 11.220185 | 3 | 46.106 | 42.513 | NaN | 4 | 0 | -4.821 |
| NCGC00139692-01 | CC1CCN(CCCNC(=O)c2cc(Sc3ccc(F)cc3)nc3ccccc23)CC1 | 11.220185 | 3 | 97.629 | 92.768 | NaN | 4 | 0 | -9.265 |
| NCGC00140328-01 | COc1ccc(Nc2cc(C(=O)NCCCN(C)Cc3ccccc3)c3ccccc3n2)cc1 | 11.220185 | 3 | 73.855 | 65.888 | 8.912509 | 4 | -9.518 | -14.105 |
| NCGC00140589-02 | COc1ccc(N2CCN(CCCNC(=O)c3cc(Nc4ccc(OC)cc4OC)nc4ccccc34)CC2)cc1 | 11.220185 | 3 | 80.089 | 73.63 | 12.589254 | 4 | -14.696 | -13.631 |
| NCGC00390036-01 | C[C@@H](Nc1nc(N2CCNCC2)nc2ccccc12)c1ccccc1 | 11.220185 | 3 | 101.554 | 97.589 | NaN | 4 | 0 | -2.565 |
| NCGC00480765-01 | CC(C)(Nc1nc(N2CCNCC2)nc2ccccc12)c1ccccc1 | 11.220185 | 3 | 52.546 | 50.584 | 5.623413 | 4 | -10.856 | -4.047 |
| NCGC00139688-01 | O=C(NCCCN1CCCCC1)c1cc(Sc2ccc(Cl)cc2)nc2ccccc12 | 11.220185 | 2.1 | 108.286 | 98.904 | 0.354813 | 4 | -16.703 | -6.002 |
| NCGC00098551-01 | CCn1c2ccccc2c2cc(CN3CCN(Cc4cccc(Cl)c4)CC3)ccc21 | 12.589254 | 3 | 105.909 | 97.042 | NaN | 4 | 0 | 0.058 |
| NCGC00109549-01 | CCCCSCCCNC(=O)C1CCN(Cc2nc(-c3ccc(CC)cc3)oc2C)CC1 | 12.589254 | 3 | 99.013 | 89.847 | 5.623413 | 4 | -8.262 | -5.218 |
| NCGC00109615-01 | CCCCNC(=O)C1CCCN(Cc2nc(-c3ccc(Cl)cc3)oc2C)C1 | 12.589254 | 3 | 51.889 | 48.21 | 7.943282 | 4 | -28.622 | -26.352 |
| NCGC00120066-01 | CCN(CC)CCNC(=O)c1[nH]c2ccccc2c1Sc1ccc(Cl)cc1 | 12.589254 | 3 | 114.77 | 104.529 | NaN | 4 | 0 | -3.175 |
| NCGC00120134-01 | COc1ccc2c(Sc3ccccc3)c(C(=O)NCCCN3CCCCC3)[nH]c2c1 | 12.589254 | 3 | 91.809 | 83.346 | 12.589254 | 4 | -17.112 | -14.26 |
| NCGC00120194-01 | COc1ccc2c(Sc3ccc(Cl)cc3)c(C(=O)NCCCN3CCCCC3)[nH]c2c1 | 12.589254 | 3 | 109.684 | 97.443 | 3.981072 | 4 | 11.507 | -2.004 |
| NCGC00121953-01 | O=C(NCCCN1CCCCCC1)c1ccc2c(c1)sc1nc(-c3ccccc3)cn12 | 12.589254 | 3 | 86.696 | 76.552 | 7.943282 | -2.4 | -33.262 | -26.052 |
| NCGC00132814-01 | CCN(CC)CCCNC(=O)c1ccc(-c2nc(CN3CCc4ccccc43)c(C)o2)cc1 | 12.589254 | 3 | 127.332 | 116.034 | NaN | 4 | 0 | -3.059 |
| NCGC00132818-01 | CCN(CC)CCNC(=O)c1ccc(-c2nc(CN3CCc4ccccc43)c(C)o2)cc1 | 12.589254 | 3 | 108.22 | 98.101 | NaN | 4 | 0 | -2.856 |
| NCGC00137630-01 | COCCCNC(=O)C1CCN(Cc2cc3ccccc3n2Cc2cccc(Cl)c2)CC1 | 12.589254 | 3 | 50.073 | 44.923 | NaN | 4 | 0 | -9.913 |
| NCGC00137604-01 | COc1cccc(CNC(=O)C2CCN(Cc3cc4ccccc4n3Cc3ccccc3)CC2)c1OC | 12.589254 | 3 | 46.828 | 41.782 | 14.125375 | 4 | -10.285 | -11.453 |
| NCGC00377200-01 | Cc1ccc2nc(C3CCCN(Cc4cccc(-c5cccnc5)c4)C3)[nH]c(=O)c2c1 | 12.589254 | 3 | 82.666 | 74.872 | NaN | 4 | 0 | -1.413 |
| NCGC00379304-01 | CC(Nc1nc(N2CCN(C)CC2)nc2ccccc12)c1ccccc1 | 12.589254 | 3 | 119.723 | 109.277 | NaN | 4 | 0 | 0.252 |
| NCGC00140661-02 | COc1ccc(Nc2cc(C(=O)NCCCN3CCN(c4ccc(OC)cc4)CC3)c3ccccc3n2)cc1 | 12.589254 | 3 | 46.812 | 46.457 | 10 | 4 | -20.596 | -21.24 |
| NCGC00400702-01 | O=C(C[C@@H]1CCNC[C@@H]1Cc1cc(-c2ccc(F)cc2)on1)N1CCC(Cc2ccccc2)CC1 | 12.589254 | 3 | 100.317 | 91.709 | NaN | 4 | 0 | -1.559 |
| NCGC00427036-01 | O=C(NCCN1CCCCC1)c1ccc(-c2ccc(N3CCSCC3)nc2)cc1 | 12.589254 | 3 | 57.448 | 52.739 | 14.125375 | 4 | -24.459 | -24.067 |
| NCGC00437792-01 | CN(C)c1cccc(C(=O)NCC2CCN(c3cc(-c4ccc(Cl)cc4)n[nH]3)CC2)c1 | 12.589254 | 3 | 74.363 | 68.736 | 14.125375 | 4 | -27.995 | -25.413 |
| NCGC00496256-01 | O=C(CCc1ccc(CC2CCN(Cc3c[nH]c4ccccc34)CC2)cc1)N1CCCC1 | 12.589254 | 3 | 36.194 | 31.848 | NaN | 4 | 0 | -11.685 |
| NCGC00110901-02 | CCN(CC)CCCNC(=O)CSc1c2c(nc3cc(Cl)ccc13)CCCC2 | 14.125375 | 3 | 95.547 | 84.039 | 2.818383 | 4 | 7.946 | 0.6 |
| NCGC00121825-01 | COc1ccc(-c2cn3c(n2)sc2cc(C(=O)NCCCN4CCCCCC4)ccc23)cc1 | 14.125375 | 3 | 78.545 | 68.517 | 10 | 4 | -28.321 | -25.267 |
| NCGC00419419-01 | CN1CCCC2(C1)CN(Cc1cccnc1)c1ccc(C(=O)NCC3CCCCC3)cc12 | 14.125375 | 3 | 89.028 | 91.673 | 19.952623 | 4 | 18.683 | 10.552 |
| NCGC00449240-01 | Clc1ccc2nc(N3CCC(NCCc4cccnc4)CC3)sc2c1 | 14.125375 | 3 | 101.112 | 89.554 | 8.912509 | 4 | -15.257 | -14.018 |
| NCGC00489837-01 | Cc1ncccc1-c1nc2cc(C(=O)NCCCc3ccccc3)ccc2o1 | 15.848932 | 3 | 45.211 | 48.21 | NaN | 4 | 0 | -10.785 |

|  | testing results in the cytophatic effect (CPE) assay |
| --- | --- |
|  | testing results in the cytotoxicity Counter-Assay |

**Supplementary Table 6.** Experimental results for the SARS-CoV-2 actives from ‘cherry-picked’ compounds retested in CPE and cytotoxicity counterscreen assay (8-point assay and in duplicates).

| **Sample ID** | **Smiles** | **AC50 (uM)** | **CC-v2** | **Efficacy** | **Max response** | **AC50 (uM)_counter-screen** | **CC-v2_counter-screen** | **Efficacy_counter-screen** | **Max response_counter-screen** |
| --- | --- | --- | --- | --- | --- | --- | --- | --- | --- |
| NCGC00100643-02 | COc1ccc(Nc2nc(Nc3ccc(OC)cc3OC)nc(N3CCOCC3)n2)c(OC)c1 | 5.011872 | 1.1 | 89.86 | 86.359 | 12.589254 | 4 | -17.148 | -18.203 |
| NCGC00384291-01 | CNCCN(C)c1nc(NC(C)c2ccccc2)c2ccccc2n1 | 6.309573 | 1.2 | 81.769 | 67.935 | NaN | 4 | 0 | -4.007 |
| NCGC00104445-01 | CCCCN(C)CCNC(=O)c1cc(-c2ccc(Cl)s2)nc2ccccc12 | 7.943282 | 1.2 | 54.555 | 40.96 | 14.125375 | -1.2 | -55.72 | -55.326 |
| NCGC00140192-01 | CCc1ccc(Sc2cc(C(=O)NCCCN3CCC(C)CC3)c3ccccc3n2)cc1 | 7.943282 | 1.1 | 89.226 | 85.496 | 14.125375 | -1.2 | -31.425 | -31.58 |
| NCGC00104471-02 | CCN(CC)CCNC(=O)c1cc(-c2ccc(Br)s2)nc2ccccc12 | 7.943282 | 2.2 | 95.22 | -0.296 | 14.125375 | -1.1 | -86.512 | -73.121 |
| NCGC00419475-01 | CN(C)CCNC(=O)c1ccc2c(c1)C1(CCN(CC3CC3)C1)CN2Cc1ccccc1 | 7.943282 | 1.1 | 98.008 | 91.994 | NaN | 4 | 0 | -7.362 |
| NCGC00487028-01 | CC(Nc1nc(N2CCN(CCN)CC2)nc2ccccc12)c1ccccc1 | 7.943282 | 2.1 | 123.165 | 78.285 | NaN | 4 | 0 | -11.491 |
| NCGC00492858-01 | Cc1ccc(Nc2ccnc(N3CCC(C(=O)NCCCc4ccccc4)CC3)n2)cc1 | 8.912509 | 1.2 | 62.015 | 59.328 | 14.125375 | -2.2 | -33.416 | -28.101 |
| NCGC00110541-02 | CCCN1CCN(CCCNC(=O)c2cc3c(Cl)nc4ccccc4c3s2)CC1 | 10 | 1.2 | 75.143 | 74.075 | 14.125375 | 4 | -29.648 | -21.157 |
| NCGC00496752-01 | Cc1ccc2c(NC(C)c3ccccc3)nc(N3CCNCC3)nc2c1 | 10 | 1.1 | 89.313 | 83.843 | 11.220185 | 4 | -14.377 | -17.361 |
| NCGC00121825-01 | COc1ccc(-c2cn3c(n2)sc2cc(C(=O)NCCCN4CCCCCC4)ccc23)cc1 | 11.220185 | 1.2 | 70.567 | 68.514 | 10 | -1.4 | -32.335 | -26.281 |
| NCGC00105925-01 | CCCN(CCC)CCNC(=O)c1cc(-c2ccc(Br)s2)nc2ccccc12 | 11.220185 | 1.2 | 81.95 | 76.585 | 14.125375 | 4 | -17.703 | -17.47 |
| NCGC00110841-01 | Cc1ccc2nc(Cl)c3cc(C(=O)NCCCN4CCC(C)CC4)sc3c2c1 | 11.220185 | 1.1 | 91.075 | 88.716 | NaN | 4 | 0 | -5.302 |
| NCGC00139688-01 | O=C(NCCCN1CCCCC1)c1cc(Sc2ccc(Cl)cc2)nc2ccccc12 | 11.220185 | 1.1 | 99.262 | 94.328 | NaN | 4 | 0 | -7.176 |
| NCGC00498993-01 | COc1ccccc1C(=O)NC1CCCN(Cc2cccc(-c3ccc(C)c(Cl)c3)c2)C1 | 12.589254 | 1.2 | 37.56 | 41.818 | 31.622777 | 4 | -32.311 | -12.16 |
| NCGC00377240-01 | COc1cccc(-c2ccc(CN3CCCC(c4nc5ccc(C)cc5c(=O)[nH]4)C3)cc2)c1 | 12.589254 | 1.4 | 45.428 | 40.349 | 28.183829 | -2.4 | -51.573 | -11.727 |
| NCGC00132818-01 | CCN(CC)CCNC(=O)c1ccc(-c2nc(CN3CCc4ccccc43)c(C)o2)cc1 | 12.589254 | 2.1 | 82.794 | 91.896 | 14.125375 | 4 | -10.025 | -13.235 |
| NCGC00140190-01 | CCc1ccc(Sc2cc(C(=O)NCCCN3CCCC(C)C3)c3ccccc3n2)cc1 | 12.589254 | 1.1 | 86.56 | 84.131 | 14.125375 | 4 | -18.694 | -20.145 |
| NCGC00109633-01 | CCCN(CCC)CCCNC(=O)C1CCCN(Cc2nc(-c3ccc(Cl)cc3)oc2C)C1 | 12.589254 | 1.2 | 87.64 | 78.705 | 14.125375 | 4 | -17.699 | -16.48 |
| NCGC00139690-01 | CC1CCCN(CCCNC(=O)c2cc(Sc3ccc(F)cc3)nc3ccccc23)C1 | 12.589254 | 1.1 | 90.753 | 92.851 | NaN | 4 | 0 | -1.863 |
| NCGC00120066-01 | CCN(CC)CCNC(=O)c1[nH]c2ccccc2c1Sc1ccc(Cl)cc1 | 12.589254 | 1.1 | 91.961 | 91.174 | NaN | 4 | 0 | -6.95 |
| NCGC00390104-01 | Cc1ccc(C(C)Nc2nc(N3CCNCC3)nc3ccccc23)cc1 | 12.589254 | 1.1 | 92.392 | 90.822 | NaN | 4 | 0 | -11.135 |
| NCGC00377241-01 | Cc1ccc(-c2cccc(CN3CCCC(c4nc5ccc(C)cc5c(=O)[nH]4)C3)c2)c(C)c1 | 12.589254 | 1.1 | 97.954 | 96.745 | NaN | 4 | 0 | -4.117 |
| NCGC00126272-01 | CCOc1ccc(NC(=O)c2cnn(-c3ccc(C)c(Cl)c3)c2C2CCNCC2)cc1 | 14.125375 | 1.2 | 30.062 | 33.913 | 19.952623 | 4 | -11.432 | -9.03 |
| NCGC00140413-01 | Cc1ccccc1CN1C(=O)c2ccccc2Sc2ccc(C(=O)NCCCN3CCOCC3)cc21 | 14.125375 | 1.2 | 32.206 | 40.818 | 14.125375 | 4 | -14.136 | -15.628 |
| NCGC00120248-01 | CCC1CCCCN1CCCNC(=O)c1sc2nc(C)cc(C)c2c1-n1cccc1 | 14.125375 | 2.2 | 35.65 | 53.233 | NaN | 4 | 0 | -8.835 |
| NCGC00498661-01 | O=C(NCc1ccccc1Cl)C1CCCN(Cc2cccc(-c3ccc(F)cc3F)c2)C1 | 14.125375 | 1.2 | 37.261 | 44.175 | 14.125375 | -2.2 | -33.487 | -18.1 |
| NCGC00116815-01 | CN(C)c1ccc(-c2nc(-c3cccs3)c(-c3cccs3)[nH]2)cc1 | 14.125375 | 2.2 | 40.59 | 56.407 | NaN | 4 | 0 | -20.003 |
| NCGC00493148-01 | COc1ccc(CC(=O)NC2CCN(Cc3ccc(-c4ccccc4OC)cc3)C2)cc1 | 14.125375 | 1.2 | 44.222 | 48.262 | 19.952623 | 4 | -11.413 | -7.42 |
| NCGC00140589-02 | COc1ccc(N2CCN(CCCNC(=O)c3cc(Nc4ccc(OC)cc4OC)nc4ccccc34)CC2)cc1 | 14.125375 | 1.2 | 46.734 | 51.725 | 14.125375 | 4 | -22.595 | -22.537 |
| NCGC00522637-01 | CC(C)(C)NCc1ccc(Nc2ccnc3cc(Cl)ccc23)cc1O | 14.125375 | 1.2 | 54.851 | 70.062 | 14.125375 | 4 | -29.465 | -33.984 |
| NCGC00426433-01 | COc1cccc(-c2ccccc2NC(=O)C2CCN(Cc3ccccc3F)CC2)c1 | 14.125375 | 1.2 | 55.79 | 61.319 | NaN | 4 | 0 | -0.611 |
| NCGC00377200-01 | Cc1ccc2nc(C3CCCN(Cc4cccc(-c5cccnc5)c4)C3)[nH]c(=O)c2c1 | 14.125375 | 2.2 | 58.439 | 69.7 | NaN | 4 | 0 | -8.213 |
| NCGC00121953-01 | O=C(NCCCN1CCCCCC1)c1ccc2c(c1)sc1nc(-c3ccccc3)cn12 | 14.125375 | 1.2 | 63.781 | 70.598 | 14.125375 | -1.2 | -30.183 | -32.509 |
| NCGC00105953-02 | CCCCN(CCCC)CCCNC(=O)c1cc(-c2ccc(Cl)s2)nc2ccccc12 | 14.125375 | 1.2 | 63.824 | 75.768 | 17.782794 | 4 | -16.131 | -12.51 |
| NCGC00449240-01 | Clc1ccc2nc(N3CCC(NCCc4cccnc4)CC3)sc2c1 | 14.125375 | 1.2 | 64.034 | 76.234 | 14.125375 | 4 | -18.146 | -17.213 |
| NCGC00139692-01 | CC1CCN(CCCNC(=O)c2cc(Sc3ccc(F)cc3)nc3ccccc23)CC1 | 14.125375 | 1.2 | 65.913 | 84.436 | NaN | 4 | 0 | -8.01 |
| NCGC00498682-01 | COc1ccccc1-c1cccc(CN2CCCC(C(=O)NCc3ccc(F)cc3)C2)c1 | 14.125375 | 1.2 | 67.172 | 74.104 | 12.589254 | 4 | -29.924 | -12.039 |
| NCGC00440956-01 | Clc1ccccc1C1(c2ccnc(-c3ccccc3)n2)CCNCC1 | 14.125375 | 1.2 | 69.584 | 82.291 | 28.183829 | 4 | -16.355 | -10.185 |
| NCGC00400702-01 | O=C(C[C@@H]1CCNC[C@@H]1Cc1cc(-c2ccc(F)cc2)on1)N1CCC(Cc2ccccc2)CC1 | 14.125375 | 1.2 | 72.596 | 85.233 | 3.981072 | 4 | 9.626 | -3.872 |
| NCGC00109549-01 | CCCCSCCCNC(=O)C1CCN(Cc2nc(-c3ccc(CC)cc3)oc2C)CC1 | 14.125375 | 1.2 | 73.005 | 80.038 | 14.125375 | 4 | -21.335 | -14.601 |
| NCGC00120134-01 | COc1ccc2c(Sc3ccccc3)c(C(=O)NCCCN3CCCCC3)[nH]c2c1 | 14.125375 | 1.2 | 73.449 | 76.007 | 15.848932 | 4 | -18.421 | -19.112 |
| NCGC00389098-01 | CC(Nc1nc(N2CCNCC2C)nc2ccccc12)c1ccccc1 | 14.125375 | 1.2 | 75.562 | 84.36 | NaN | 4 | 0 | -10.058 |
| NCGC00098551-01 | CCn1c2ccccc2c2cc(CN3CCN(Cc4cccc(Cl)c4)CC3)ccc21 | 14.125375 | 1.2 | 79.455 | 88.273 | NaN | 4 | 0 | -3.35 |
| NCGC00427036-01 | O=C(NCCN1CCCCC1)c1ccc(-c2ccc(N3CCSCC3)nc2)cc1 | 14.125375 | 1.1 | 81.074 | 83.378 | NaN | 4 | 0 | -5.878 |
| NCGC00479261-01 | C[C@@H](Nc1nc(N2CCNCC2)nc2ccccc12)c1ccc(F)cc1 | 14.125375 | 1.1 | 81.301 | 89.118 | 14.125375 | 4 | -12.942 | -15.443 |
| NCGC00120194-01 | COc1ccc2c(Sc3ccc(Cl)cc3)c(C(=O)NCCCN3CCCCC3)[nH]c2c1 | 14.125375 | 1.1 | 81.531 | 90.118 | 14.125375 | -3 | -75.639 | -1.675 |
| NCGC00132814-01 | CCN(CC)CCCNC(=O)c1ccc(-c2nc(CN3CCc4ccccc43)c(C)o2)cc1 | 14.125375 | 1.1 | 81.819 | 97.248 | NaN | 4 | 0 | -0.173 |
| NCGC00113431-01 | CCCCN(CCCC)CCCNC(=O)c1ccc2nc(N3CCCCC3)sc2c1 | 14.125375 | 1.1 | 83.675 | 86.482 | NaN | 4 | 0 | -7.193 |
| NCGC00109669-01 | Cc1oc(-c2cccc(Cl)c2)nc1CN1CCCC(C(=O)NCCc2ccc(Cl)cc2)C1 | 25.118864 | 3 | 43.391 | 34.045 | 14.125375 | -1.2 | -44.328 | -45.623 |
| NCGC00489837-01 | Cc1ncccc1-c1nc2cc(C(=O)NCCCc3ccccc3)ccc2o1 | 25.118864 | 2.2 | 50.97 | 38.177 | NaN | 4 | 0 | -8.085 |
| NCGC00416827-01 | COc1cc2c(cc1OCC1CC1)NC(=O)C21CCN(c2cc(NC3CC3)ncn2)CC1 | 25.118864 | 2.2 | 73.039 | 57.54 | NaN | 4 | 0 | -10.135 |
| NCGC00419419-01 | CN1CCCC2(C1)CN(Cc1cccnc1)c1ccc(C(=O)NCC3CCCCC3)cc12 | 25.118864 | 2.1 | 124.124 | 97.373 | NaN | 4 | 0 | -3.806 |

|  | testing results in the cytophatic effect (CPE) assay |
| --- | --- |
|  | testing results in the cytotoxicity Counter-Assay |

**Supplementary Table 7.** Experimental results for all virtual screening hits tested in CPE and counter-screen assay (5-point assay).

| **Sample ID** | **Smiles** | AC50 (uM) | CC-v2 | Efficacy | Max response | AC50 (uM)_counter-screen | CC-v2_counter-screen | Efficacy_counter-screen | Max response_counter-screen |
| --- | --- | --- | --- | --- | --- | --- | --- | --- | --- |
| NCGC00099741-01 | Cc1cc(OCC(O)CN2CCN(c3cccc(Cl)c3)CC2)ccc1Cl | 12.589254 | 3 | 88.291 | 80.373 | NaN | 4 | 0 | -6.691 |
| NCGC00102225-01 | COc1ccc(Cl)cc1NC(=O)CSc1nc2ccccc2nc1N1CCCCC1 | 12.589254 | 3 | 100.946 | 92.034 | NaN | 4 | 0 | -1.769 |
| NCGC00104445-01 | CCCCN(C)CCNC(=O)c1cc(-c2ccc(Cl)s2)nc2ccccc12 | 4.466836 | 1.2 | 76.459 | 75.466 | 14.125375 | 4 | -15.091 | -14.659 |
| NCGC00104471-02 | CCN(CC)CCNC(=O)c1cc(-c2ccc(Br)s2)nc2ccccc12 | 3.548134 | 1.3 | 84.477 | 71.952 | 14.125375 | 4 | -15.243 | -14.369 |
| NCGC00103820-01 | CCN1CCN(c2cc(C)c3cc(NC(=O)c4cccc(Cl)c4)ccc3n2)CC1 | 14.125375 | 3 | 107.678 | 94.851 | NaN | 4 | 0 | -0.887 |
| NCGC00109669-01 | Cc1oc(-c2cccc(Cl)c2)nc1CN1CCCC(C(=O)NCCc2ccc(Cl)cc2)C1 | 12.589254 | 3 | 49.891 | 43.844 | 12.589254 | 4 | -18.612 | -18.629 |
| NCGC00116825-01 | Oc1ccccc1-c1nc(-c2cccs2)c(-c2cccs2)[nH]1 | 12.589254 | 3 | 52.719 | 47.993 | 14.125375 | 4 | -16.903 | -16.905 |
| NCGC00116815-01 | CN(C)c1ccc(-c2nc(-c3cccs3)c(-c3cccs3)[nH]2)cc1 | 12.589254 | 3 | 45.173 | 41.118 | NaN | 4 | 0 | -1.007 |
| NCGC00117038-01 | Cc1c(Cl)cccc1-c1ccc(CNCCSc2nnnn2-c2ccccc2)o1 | 14.125375 | 3 | 43.041 | 37.574 | NaN | 4 | 0 | -1.629 |
| NCGC00114893-01 | CCc1ccccc1N1CC(c2nc3ccccc3n2CCCCOc2ccccc2OC)CC1=O | 10 | 3 | 39.31 | 35.544 | 15.848932 | 4 | -10.618 | -10.099 |
| NCGC00120248-01 | CCC1CCCCN1CCCNC(=O)c1sc2nc(C)cc(C)c2c1-n1cccc1 | 12.589254 | 3 | 102.132 | 90.489 | NaN | 4 | 0 | -2 |
| NCGC00126272-01 | CCOc1ccc(NC(=O)c2cnn(-c3ccc(C)c(Cl)c3)c2C2CCNCC2)cc1 | 12.589254 | 3 | 99.715 | 88.854 | NaN | 4 | 0 | -8.174 |
| NCGC00126319-01 | Cc1ccc(Cl)cc1N1CCN(CCCNc2ncnc3onc(-c4ccc(F)cc4)c23)CC1 | 11.220185 | 3 | 72.129 | 65.803 | NaN | 4 | 0 | -1.789 |
| NCGC00127046-01 | CCOc1ccccc1CNCc1cccn1-c1nnc(N2CCN(c3ccccc3)CC2)s1 | 3.162278 | 1.2 | 42.383 | 40.209 | 14.125375 | 4 | -25.051 | -22.959 |
| NCGC00139668-01 | CC1CCCCN1CCCNC(=O)c1cc(Sc2ccc(Cl)cc2)nc2ccccc12 | 6.309573 | 2.1 | 102.706 | 91.731 | NaN | 4 | 0 | -1.679 |
| NCGC00139670-01 | CCCN(CCC)CCNC(=O)c1cc(Sc2ccc(Cl)cc2)nc2ccccc12 | 10 | 2.1 | 97.171 | 86.854 | NaN | 4 | 0 | -3.062 |
| NCGC00140192-01 | CCc1ccc(Sc2cc(C(=O)NCCCN3CCC(C)CC3)c3ccccc3n2)cc1 | 3.548134 | 1.2 | 62.383 | 54.142 | 11.220185 | 4 | -18.552 | -21.906 |
| NCGC00140311-01 | COc1ccc(CCNC(=O)c2cc(Nc3ccccc3)nc3ccccc23)cc1OC | 12.589254 | 3 | 52.14 | 46.358 | NaN | 4 | 0 | -5.267 |
| NCGC00140190-01 | CCc1ccc(Sc2cc(C(=O)NCCCN3CCCC(C)C3)c3ccccc3n2)cc1 | 11.220185 | 2.1 | 102.202 | 91.58 | NaN | 4 | 0 | -1.679 |
| NCGC00139690-01 | CC1CCCN(CCCNC(=O)c2cc(Sc3ccc(F)cc3)nc3ccccc23)C1 | 12.589254 | 3 | 109.007 | 97.607 | NaN | 4 | 0 | 4.355 |
| NCGC00140356-01 | Cc1ccc(-c2nc(C[S+]([O-])CC(=O)N3CCN(c4ccc(Cl)cc4)CC3)c(C)o2)cc1 | 3.548134 | 1.2 | 36.906 | 32.849 | 14.125375 | 4 | -16.907 | -16.995 |
| NCGC00140413-01 | Cc1ccccc1CN1C(=O)c2ccccc2Sc2ccc(C(=O)NCCCN3CCOCC3)cc21 | 11.220185 | 3 | 68.651 | 65.622 | NaN | 4 | 0 | -9.958 |
| NCGC00377203-01 | COc1ccccc1-c1ccc(CN2CCCC(c3nc4ccc(C)cc4c(=O)[nH]3)C2)cc1 | 11.220185 | 2.1 | 93.787 | 89.187 | 12.589254 | -2.4 | -32.682 | -26.818 |
| NCGC00411503-01 | Cc1noc2c(NC3CCOCC3)cc(-c3cccc(C(F)(F)F)c3)nc12 | 15.848932 | 3 | 41.334 | 41.723 | 14.125375 | 4 | -21.682 | -22.949 |
| NCGC00399676-01 | CN(C)c1ccc(C(=O)NC[C@H]2C[C@@H]3CCN2C[C@@H]3c2cc(C3CCCCC3)nn2C)cc1 | 14.125375 | 3 | 99.867 | 87.793 | NaN | 4 | 0 | -5.999 |
| NCGC00140464-01 | Cc1cccc(N2CCN(C(=O)C3CCN(Cc4nc(-c5ccc(Cl)cc5)oc4C)CC3)CC2)c1C | 12.589254 | 3 | 118.512 | 108.39 | NaN | 4 | 0 | -2.26 |
| NCGC00416827-01 | COc1cc2c(cc1OCC1CC1)NC(=O)C21CCN(c2cc(NC3CC3)ncn2)CC1 | 15.848932 | 3 | 71.885 | 76.496 | NaN | 4 | 0 | 0.236 |
| NCGC00426433-01 | COc1cccc(-c2ccccc2NC(=O)C2CCN(Cc3ccccc3F)CC2)c1 | 12.589254 | 3 | 98.98 | 88.339 | NaN | 4 | 0 | -1.679 |
| NCGC00440956-01 | Clc1ccccc1C1(c2ccnc(-c3ccccc3)n2)CCNCC1 | 14.125375 | 3 | 109.327 | 96.153 | NaN | 4 | 0 | -4.626 |
| NCGC00427056-01 | Cn1cc(CN2CCCC(COc3ccccc3C(=O)N3CCCCC3)C2)c2ccccc21 | 12.589254 | 3 | 92.629 | 83.28 | NaN | 4 | 0 | -5.849 |
| NCGC00490405-01 | Cc1ccc(Nc2ccnc(N3CCCC3C(=O)NCCCc3ccccc3)n2)cc1 | 12.589254 | 3 | 39.935 | 36.09 | NaN | 4 | 0 | -7.733 |
| NCGC00493147-01 | COc1ccccc1-c1ccc(CN2CCC(NC(=O)Cc3ccc(F)cc3)C2)cc1 | 12.589254 | 3 | 98.957 | 90.58 | NaN | 4 | 0 | -0.987 |
| NCGC00498636-01 | O=C(NCc1cccs1)C1CCCN(Cc2cccc(-c3ccc(F)cc3F)c2)C1 | 14.125375 | 3 | 82.732 | 72.74 | NaN | 4 | 0 | -10.44 |
| NCGC00498641-01 | O=C(Nc1cccc(Cl)c1)C1CCCN(Cc2cccc(-c3ccc(F)cc3F)c2)C1 | 12.589254 | 3 | 47.265 | 42.117 | NaN | 4 | 0 | -9.457 |
| NCGC00498661-01 | O=C(NCc1ccccc1Cl)C1CCCN(Cc2cccc(-c3ccc(F)cc3F)c2)C1 | 12.589254 | 3 | 52.124 | 47.388 | 14.125375 | 4 | -11.422 | -14.108 |
| NCGC00493148-01 | COc1ccc(CC(=O)NC2CCN(Cc3ccc(-c4ccccc4OC)cc3)C2)cc1 | 14.125375 | 3 | 101.769 | 87.975 | NaN | 4 | 0 | -4.856 |
| NCGC00498682-01 | COc1ccccc1-c1cccc(CN2CCCC(C(=O)NCc3ccc(F)cc3)C2)c1 | 12.589254 | 3 | 70.18 | 62.078 | 14.125375 | 4 | -18.663 | -20.734 |
| NCGC00498749-01 | Cc1cc(C)cc(-c2ccc(CN3CCCC3C(=O)NCc3cccc(F)c3)cc2)c1 | 12.589254 | 3 | 87.111 | 78.283 | 14.125375 | 4 | -11.208 | -12.825 |
| NCGC00498687-01 | COc1ccccc1-c1cccc(CN2CCCC(C(=O)NCCc3ccc(F)cc3)C2)c1 | 12.589254 | 3 | 79.852 | 70.286 | 7.943282 | 4 | -12.016 | -14.379 |
| NCGC00498981-01 | Cc1ccc(-c2cccc(CN3CCCC(NC(=O)c4c(C)noc4C)C3)c2)cc1Cl | 11.220185 | 2.1 | 101.374 | 92.579 | NaN | 4 | 0 | -2.441 |
| NCGC00498993-01 | COc1ccccc1C(=O)NC1CCCN(Cc2cccc(-c3ccc(C)c(Cl)c3)c2)C1 | 12.589254 | 3 | 75.346 | 65.531 | 15.848932 | 4 | -11.507 | -12.194 |
| NCGC00510147-03 | CC(C)(C)N1CCC(c2ccccc2)(c2ccccc2)CC1 | 10 | 2.2 | 50.396 | 40.663 | NaN | 4 | 0 | -4.245 |
| NCGC00100886-01 | Cc1ccc(Sc2c(C)nn(C(=O)COc3ccccc3Br)c2C)cc1 | NaN | 4 | 0 | 3.165 | NaN | 4 | 0 | -5.388 |
| NCGC00100448-01 | Nc1c(C(=O)Nc2ccc(F)c(F)c2)sc2nc(-c3ccncc3)ccc12 | NaN | 4 | 0 | -0.742 | NaN | 4 | 0 | -0.616 |
| NCGC00098388-01 | O=C(NC1CCCCC1)/C(=C/c1ccc(Cl)cc1)NC(=O)c1ccccc1 | NaN | 4 | 0 | -0.106 | NaN | 4 | 0 | 0.165 |
| NCGC00101300-01 | COc1ccc(Cl)cc1NC(=O)CSc1nc(-c2cccs2)cc(C(F)(F)F)n1 | NaN | 4 | 0 | 2.196 | NaN | 4 | 0 | -2.681 |
| NCGC00100234-01 | C#CCOc1ccc(-c2nnc(Nc3cccc(C)c3)c3ccccc23)cc1 | NaN | 4 | 0 | -1.56 | NaN | 4 | 0 | 3.062 |
| NCGC00099930-01 | CC(C)CSc1nnc(-c2ccccc2)c(-c2ccccc2)n1 | NaN | 4 | 0 | 0.924 | NaN | 4 | 0 | -7.172 |
| NCGC00099934-01 | Cc1ccnc(NC(c2cc(C)ccc2O)c2ccc3cccnc3c2O)c1 | 3.548134 | 1.2 | 33.702 | 31.91 | 8.912509 | -2.2 | -63.849 | -58.382 |
| NCGC00100272-01 | CCCCc1nc2ccccc2n1CC(=O)N(COCC)c1c(C)cccc1CC | NaN | 4 | 0 | 1.227 | NaN | 4 | 0 | -6.33 |
| NCGC00099929-01 | CCCCN(CCCC)C1=C(C(=O)c2ccccc2)N(CC)[S+](=O)([O-])c2ccccc21 | NaN | 4 | 0 | -0.045 | 11.220185 | -3 | -36.539 | -36.35 |
| NCGC00100257-01 | CC(C)c1ccccc1NC(=O)C(Cc1c[nH]c2ccccc12)NC(=O)OC(C)(C)C | NaN | 4 | 0 | 1.166 | NaN | 4 | 0 | -2.33 |
| NCGC00100027-01 | CCOC(=O)c1c(Nc2cc(C)cc(C)c2)nnc(-c2ccccc2)c1-c1ccccc1 | NaN | 4 | 0 | 4.407 | NaN | 4 | 0 | -2.391 |
| NCGC00103228-01 | COc1cccc(NC(=O)Nc2cccc(-c3cn4cccnc4n3)c2)c1 | NaN | 4 | 0 | -0.621 | NaN | 4 | 0 | 0.957 |
| NCGC00101881-01 | CCc1ccc(OCC(=O)Nc2c(C(=O)c3ccc(Br)cc3)oc3ccccc23)cc1 | NaN | 4 | 0 | -0.984 | NaN | 4 | 0 | -1.408 |
| NCGC00102171-01 | COc1ccc(NC(=O)CSc2nc3ccccc3nc2N2CCC(C)CC2)cc1 | NaN | 4 | 0 | 6.618 | NaN | 4 | 0 | -0.386 |
| NCGC00104665-01 | COCCNC(=O)CCCn1c(=O)[nH]c2ccsc2c1=O | NaN | 4 | 0 | 1.136 | NaN | 4 | 0 | -1.007 |
| NCGC00103824-02 | CCN1CCN(c2cc(C)c3cc(NC(=O)c4cc(Cl)ccc4OC)ccc3n2)CC1 | 4.466836 | 3 | 78.883 | -4.437 | 12.589254 | -3 | -110.199 | -98.998 |
| NCGC00104579-01 | CSCCC(NC(=O)C1CCCCC1)C(=O)Nc1ccc(F)c(Cl)c1 | NaN | 4 | 0 | 1.863 | NaN | 4 | 0 | 0.466 |
| NCGC00103746-01 | CCOCCCNC(=O)C(c1ccc(OC)cc1)N(Cc1cccs1)C(=O)CNC(C)=O | NaN | 4 | 0 | -1.62 | NaN | 4 | 0 | 2.08 |
| NCGC00105308-01 | Cc1ccc(Sc2ccc(N3CCCC(C(=O)Nc4ccc(F)c(Cl)c4)C3)nn2)cc1 | NaN | 4 | 0 | -1.56 | NaN | 4 | 0 | 2.451 |
| NCGC00106814-01 | Cc1ccc(-c2nc3ccc(Cl)cn3c2Nc2ccccc2C)s1 | NaN | 4 | 0 | -0.167 | NaN | 4 | 0 | -3.483 |
| NCGC00105152-01 | Cc1ccc(C)c(N2CCN(c3nccn4nc(-c5ccc(Br)cc5)cc34)CC2)c1 | NaN | 4 | 0 | 0.5 | NaN | 4 | 0 | -0.787 |
| NCGC00103822-01 | CCCCc1ccc(C(=O)Nc2ccc3nc(N4CCN(CC)CC4)cc(C)c3c2)cc1 | 4.466836 | 3 | 76.862 | 1.499 | NaN | 4 | 0 | 1.017 |
| NCGC00105334-01 | Cc1ccc(Sc2ccc(N3CCCC(C(=O)Nc4ccc(Br)cc4)C3)nn2)cc1 | NaN | 4 | 0 | -1.923 | NaN | 4 | 0 | 0.837 |
| NCGC00104635-01 | CCCCC(CC)CNC(=O)CCCCCn1c(=O)[nH]c2ccsc2c1=O | NaN | 4 | 0 | -1.681 | NaN | 4 | 0 | 2.852 |
| NCGC00106556-01 | Cc1c(NC(=O)Nc2ccc(Cl)c(C(F)(F)F)c2)c(=O)n(-c2ccccc2)n1C | NaN | 4 | 0 | -0.469 | NaN | 4 | 0 | 1.098 |
| NCGC00106963-01 | CCOc1cc2c(cc1OCC)C(c1c(F)cccc1Cl)CC(=O)N2 | NaN | 4 | 0 | -2.378 | NaN | 4 | 0 | -3.403 |
| NCGC00105312-01 | CCOc1ccc(NC(=O)C2CCCN(c3ccc(Sc4ccc(C)cc4)nn3)C2)cc1 | NaN | 4 | 0 | -1.136 | NaN | 4 | 0 | 1.539 |
| NCGC00105336-01 | Cc1ccc(Sc2ccc(N3CCCC(C(=O)Nc4ccc(F)cc4)C3)nn2)cc1 | NaN | 4 | 0 | -2.983 | NaN | 4 | 0 | -3.223 |
| NCGC00105927-01 | CCCCN(C)CCCNC(=O)c1cc(-c2ccc(Br)s2)nc2ccccc12 | 4.466836 | 3 | 104.699 | -4.801 | 12.589254 | -2.1 | -106.322 | -99.599 |
| NCGC00108948-01 | Cc1ccc(-c2nc3ccc(Cl)cn3c2Nc2ccc3c(c2)OCCO3)s1 | NaN | 4 | 0 | 0.833 | NaN | 4 | 0 | -1.118 |
| NCGC00107872-01 | COc1cccc(-c2nc(SCC(=O)NC3CCCC3)c(-c3ccc(C)cc3)[nH]2)c1 | NaN | 4 | 0 | -0.772 | NaN | 4 | 0 | 0.897 |
| NCGC00108963-01 | CCN(CCCNC(=O)CSc1cc(=O)n(C)c2cc(Cl)ccc12)c1ccccc1 | NaN | 4 | 0 | -1.56 | NaN | 4 | 0 | -3.563 |
| NCGC00108911-01 | CCN(CCCNC(=O)CSc1cc(=O)n(C)c2ccccc12)c1ccccc1 | NaN | 4 | 0 | -0.682 | NaN | 4 | 0 | -3.393 |
| NCGC00108134-01 | COc1ccc(Nc2cc(C(=O)NCCc3ccccc3)c3ccccc3n2)cc1OC | NaN | 4 | 0 | 3.226 | NaN | 4 | 0 | 0.316 |
| NCGC00109503-01 | Cc1ccc(-c2nc(C[S+]([O-])CC(=O)N3CCN(c4ccccc4)CC3)c(C)o2)cc1 | 10 | 2.4 | 33.878 | 27.488 | NaN | 4 | 0 | -4.987 |
| NCGC00109613-01 | Cc1oc(-c2ccc(Cl)cc2)nc1CN1CCCC(C(=O)NCCc2ccccc2)C1 | 3.162278 | 4 | 11.631 | 9.193 | 4.466836 | 4 | -15.046 | -19.19 |
| NCGC00109521-02 | Cc1ccc(-c2nc(C[S+]([O-])CC(=O)N3CCN(c4cccc(Cl)c4)CC3)c(C)o2)cc1 | NaN | 4 | 0 | 8.072 | 14.125375 | 4 | -11.532 | -7.944 |
| NCGC00109611-01 | Cc1oc(-c2ccc(Cl)cc2)nc1CN1CCCC(C(=O)NCCc2ccc(Cl)cc2)C1 | 3.548134 | 4 | 14.396 | 10.404 | 14.125375 | 4 | -27.973 | -24.623 |
| NCGC00110909-01 | COc1cccc(CNC(=O)CSc2c3c(nc4cc(Cl)ccc24)CCCC3)c1 | NaN | 4 | 0 | 3.468 | NaN | 4 | 0 | 4.055 |
| NCGC00111780-01 | Cc1ccc(-n2c(=O)c3sc4ccccc4c3n(C)c2=O)cc1F | NaN | 4 | 0 | 3.196 | NaN | 4 | 0 | 2.361 |
| NCGC00110228-01 | CCN(CCCNC(=O)CSc1cc(=O)n(CC)c2ccccc12)c1ccccc1 | NaN | 4 | 0 | 1.772 | NaN | 4 | 0 | -4.977 |
| NCGC00109507-01 | Cc1ccc(Cl)cc1N1CCN(C(=O)C2CCN(Cc3nc(-c4ccc(Cl)cc4)oc3C)CC2)CC1 | 4.466836 | 3 | 59.047 | -4.71 | 14.125375 | -3 | -112.292 | -99.379 |
| NCGC00109671-02 | Cc1ccc(CCNC(=O)C2CCCN(Cc3nc(-c4cccc(Cl)c4)oc3C)C2)cc1 | NaN | 4 | 0 | -0.621 | NaN | 4 | 0 | -4.546 |
| NCGC00111776-01 | COc1cccc(-n2c(=O)c3sc4ccccc4c3n(CC(=O)NC3CCCCC3)c2=O)c1 | NaN | 4 | 0 | 3.559 | NaN | 4 | 0 | -1.659 |
| NCGC00112060-01 | CC1=Nc2cc(C(=O)NCCc3ccccc3)ccc2Sc2c(C)ccc(C)c21 | 4.466836 | 4 | 23.459 | -1.681 | 2.511886 | -1.2 | -30.327 | -35.118 |
| NCGC00112051-01 | COc1ccc(CCNC(=O)c2ccc3c(c2)N(Cc2ccccc2)CCS3)cc1OC | NaN | 4 | 0 | -1.893 | NaN | 4 | 0 | -0.657 |
| NCGC00113553-01 | CCN1CCSc2ccc(C(=O)NCc3cccc(Br)c3)cc21 | NaN | 4 | 0 | 1.136 | NaN | 4 | 0 | -2.631 |
| NCGC00112085-01 | COc1ccc(CNC(=O)c2ccc3c(c2)N(Cc2ccc(C)cc2)CCS3)cc1 | NaN | 4 | 0 | -1.348 | NaN | 4 | 0 | -1.859 |
| NCGC00109125-01 | CCOc1ccccc1NC(=O)CSc1c2c(nc3ccc(Cl)cc13)CCCC2 | NaN | 4 | 0 | -2.862 | NaN | 4 | 0 | 1.749 |
| NCGC00113555-01 | CCN1CCSc2ccc(C(=O)NCc3ccccc3Br)cc21 | NaN | 4 | 0 | -0.682 | NaN | 4 | 0 | -3.573 |
| NCGC00111928-01 | COc1ccc(CCNC(=O)c2ccc3c(c2)N(Cc2ccc(Cl)cc2)CCS3)c(OC)c1 | NaN | 4 | 0 | 1.56 | NaN | 4 | 0 | -0.356 |
| NCGC00114605-01 | O=C(Nc1cc(-c2ccccc2)sc1C(=O)O)c1cccc(F)c1 | 10 | 4 | 14.77 | 12.494 | 14.125375 | 4 | -23.544 | -23.009 |
| NCGC00108908-01 | CCOc1ccc(CCNC(=O)c2nn(-c3ccc(OC)c(Cl)c3)c(=O)c3c2c2ccccc2n3C)cc1OCC | NaN | 4 | 0 | 0.682 | NaN | 4 | 0 | -4.295 |
| NCGC00113050-01 | CCCOc1ccc(CNC(=O)CCCn2ncn3c(cc4sc(CC)cc43)c2=O)cc1 | NaN | 4 | 0 | -0.348 | NaN | 4 | 0 | -0.145 |
| NCGC00114703-02 | Cc1cccc(OCCCn2c(CCNC(=O)C3CCCCC3)nc3ccccc32)c1 | NaN | 4 | 0 | 1.439 | 0.501187 | 4 | 9.353 | -0.075 |
| NCGC00114705-02 | Cc1cc(C)cc(OCCCn2c(CCNC(=O)C3CCCCC3)nc3ccccc32)c1 | NaN | 4 | 0 | 8.708 | NaN | 4 | 0 | 7.593 |
| NCGC00113077-01 | Cc1ccc(CNC(=O)C2CCCN(c3nc4c(C)cc(C)cc4s3)C2)cc1 | NaN | 4 | 0 | -1.469 | NaN | 4 | 0 | -0.667 |
| NCGC00114765-01 | Cc1ccc(Cn2c(CCCNC(=O)C3CCCCC3)nc3ccccc32)cc1 | NaN | 4 | 0 | 0.197 | NaN | 4 | 0 | 1.779 |
| NCGC00113267-02 | Cc1ccc(NCc2cccn2-c2nnc(N3CCC(C(=O)NCc4ccc(OC(C)C)cc4)CC3)s2)cc1 | 14.125375 | 4 | 30.202 | 27.275 | 14.125375 | 4 | -15.313 | -12.344 |
| NCGC00115601-01 | CCCCC12CN3CC(C)(CN(C1)C3c1ccccn1)C2=O | NaN | 4 | 0 | 0.197 | NaN | 4 | 0 | 0.266 |
| NCGC00114737-01 | O=C(NCCCc1nc2ccccc2n1Cc1ccccc1)C1CCCCC1 | NaN | 4 | 0 | -0.772 | NaN | 4 | 0 | -0.837 |
| NCGC00114679-01 | Cc1cccc(C)c1OCCCn1c(CCNC(=O)C2CCCCC2)nc2ccccc21 | 14.125375 | 4 | 14.742 | 10.495 | 14.125375 | 4 | -13.06 | -12.134 |
| NCGC00114369-01 | CCCN1CCN(CCCNC(=O)C(CC)n2nc(C)n3c(cc4occc43)c2=O)CC1 | NaN | 4 | 0 | 0.742 | NaN | 4 | 0 | 1.228 |
| NCGC00115663-01 | Cc1nnc(SCC(=O)Nc2ccc(N(C)C)cc2)nc1O | 0.316228 | 4 | -15.591 | 1.015 | NaN | 4 | 0 | 0.917 |
| NCGC00116382-01 | CCc1cccc2c(-c3ccccc3)nc(SCC(=O)Nc3ccc(OC)cc3)nc12 | NaN | 4 | 0 | 0.409 | NaN | 4 | 0 | -1.238 |
| NCGC00114711-01 | Cc1cccc(C)c1OCCn1c(CCNC(=O)C2CCCCC2)nc2ccccc21 | NaN | 4 | 0 | -3.014 | NaN | 4 | 0 | -8.254 |
| NCGC00117490-01 | Cc1ccc(Cl)cc1NC(=O)C(C)n1nc(-c2ccccc2)ccc1=O | NaN | 4 | 0 | 0.076 | NaN | 4 | 0 | -1.519 |
| NCGC00117518-01 | CCC(C(=O)Nc1cccc(C#N)c1)n1nc(-c2ccccc2)ccc1=O | NaN | 4 | 0 | -2.287 | 3.548134 | 2.4 | 32.388 | 2.22 |
| NCGC00117494-01 | Cc1ccc(C)c(-c2ccc(=O)n(C(C)C(=O)Nc3ccccc3C(F)(F)F)n2)c1 | NaN | 4 | 0 | -2.408 | NaN | 4 | 0 | 1.208 |
| NCGC00117520-01 | CCC(C(=O)Nc1ccccc1)n1nc(-c2cc(C)ccc2C)ccc1=O | NaN | 4 | 0 | -2.529 | NaN | 4 | 0 | 2.32 |
| NCGC00117498-01 | CCC(C(=O)Nc1cccc(Cl)c1C)n1nc(-c2ccc(C)cc2)ccc1=O | NaN | 4 | 0 | 0.379 | NaN | 4 | 0 | -0.737 |
| NCGC00118053-01 | Cn1nnnc1SCCCNCc1ccc(-c2ccc(F)c(Cl)c2)o1 | 14.125375 | 2.4 | 32.657 | 26.549 | NaN | 4 | 0 | -11.733 |
| NCGC00109960-01 | CSc1ccc(CCNC(=O)CCc2c(C)nc3cc(-c4ccccc4)nn3c2C)cc1 | NaN | 4 | 0 | 0.379 | NaN | 4 | 0 | 0.877 |
| NCGC00112089-01 | Cc1ccc(CN2CCSc3ccc(C(=O)Nc4cc(C)cc(C)c4)cc32)cc1 | NaN | 4 | 0 | 0.045 | NaN | 4 | 0 | -4.786 |
| NCGC00114794-01 | CCOc1ccc(N(C)C(CN[S+](=O)([O-])c2ccccc2)c2ccccc2)cc1 | 14.125375 | 2.4 | 30.51 | 24.883 | NaN | 4 | 0 | 4.165 |
| NCGC00114683-02 | O=C(NCCc1nc2ccccc2n1CCCOc1ccc(Cl)cc1)C1CCCCC1 | NaN | 4 | 0 | 2.893 | NaN | 4 | 0 | 2.741 |
| NCGC00114932-01 | Cc1cccc(C)c1NC(=O)N1CCC(C(N)=O)(N2CCCCC2)CC1 | NaN | 4 | 0 | -0.833 | NaN | 4 | 0 | -0.226 |
| NCGC00118037-01 | CCOc1ccc(NC(=O)C(CC)n2nc(-c3ccc(C)c(C)c3)ccc2=O)cc1 | NaN | 4 | 0 | 0.227 | NaN | 4 | 0 | 2.701 |
| NCGC00114879-01 | CCn1cc(/C=C(\NC(=O)c2ccc(C)cc2)C(=O)NCCCn2ccnc2)c2ccccc21 | NaN | 4 | 0 | -2.135 | NaN | 4 | 0 | -0.025 |
| NCGC00118061-01 | CCC(C(=O)Nc1ccc(C(=O)OC)cc1)n1nc(-c2ccc(C)c(C)c2)ccc1=O | NaN | 4 | 0 | 1.408 | NaN | 4 | 0 | 2.32 |
| NCGC00118494-01 | Cc1cccc(OCCSc2nc3ccc(NC(=O)COc4ccccc4)cc3s2)c1 | NaN | 4 | 0 | 0.227 | NaN | 4 | 0 | -0.065 |
| NCGC00118231-01 | COc1ccc(NC(=O)C(C)Sc2cc(-c3ccccc3)nc(C)n2)cc1Cl | NaN | 4 | 0 | 1.318 | NaN | 4 | 0 | -1.659 |
| NCGC00118269-01 | CC(C)(C)c1ccc(C(=O)NCC(c2ccc(Cl)cc2)N2CCOCC2)cc1 | NaN | 4 | 0 | -0.136 | NaN | 4 | 0 | -1.388 |
| NCGC00120008-01 | O=C(Cn1ncn2nc(-c3cccs3)cc2c1=O)NCc1cccc(Br)c1 | NaN | 4 | 0 | -0.409 | NaN | 4 | 0 | -1.418 |
| NCGC00118946-01 | COc1ccccc1OC(C)C(=O)N1CCN(CCc2ccccn2)CC1 | NaN | 4 | 0 | -1.318 | NaN | 4 | 0 | -0.356 |
| NCGC00119297-01 | CCc1ccc(OCCSc2nc3ccccc3n2CC(=O)N2CCCCC2)cc1 | NaN | 4 | 0 | -2.378 | NaN | 4 | 0 | 0.025 |
| NCGC00119508-01 | CCc1noc(-c2ccc(NC(C)c3ccccc3)c([N+](=O)[O-])c2)n1 | 11.220185 | 4 | 16.667 | 16.402 | NaN | 4 | 0 | 2.651 |
| NCGC00119380-01 | COc1ccc(-c2noc(-c3ccc(NC(C)c4ccccc4)c([N+](=O)[O-])c3)n2)cc1OC | NaN | 4 | 0 | 0.348 | NaN | 4 | 0 | -0.907 |
| NCGC00118063-01 | CCc1ccccc1NC(=O)C(C)n1nc(-c2ccc(C)c(C)c2)ccc1=O | NaN | 4 | 0 | -0.379 | NaN | 4 | 0 | 0.486 |
| NCGC00119494-01 | CCOc1ccc(NC(=O)CSc2nnc(-c3ccc(OCC)cc3)o2)cc1 | NaN | 4 | 0 | -1.257 | NaN | 4 | 0 | 0.296 |
| NCGC00117524-01 | CCC(C(=O)Nc1cc(F)ccc1F)n1nc(-c2cc(C)ccc2C)ccc1=O | NaN | 4 | 0 | 1.802 | NaN | 4 | 0 | 5.839 |
| NCGC00120580-01 | Cc1c(NC(=O)Cc2ccc(F)cc2)cccc1-c1nc2cccnc2s1 | NaN | 4 | 0 | 0.076 | NaN | 4 | 0 | -2.371 |
| NCGC00120416-01 | CC(C)CCNC(=O)c1nn(C)c2c1CSc1ccccc1-2 | NaN | 4 | 0 | -1.439 | NaN | 4 | 0 | -1.118 |
| NCGC00121502-01 | Cc1ccc(CNC(=O)C2CCCN(c3nc4ccccc4n4cccc34)C2)cc1 | NaN | 4 | 0 | -0.772 | NaN | 4 | 0 | -1.348 |
| NCGC00120072-01 | CC1CCCCN1CCCNC(=O)c1[nH]c2ccccc2c1Sc1ccc(Cl)cc1 | 5.011872 | 4 | 12.359 | -4.771 | 14.125375 | -3 | -112.769 | -99.358 |
| NCGC00121681-01 | CC1CCCCN1CCCNC(=O)c1ccc2c(c1)sc1nc(-c3ccc(F)cc3)cn12 | 4.466836 | 3 | 112.683 | -4.559 | 12.589254 | -3 | -106.866 | -99.268 |
| NCGC00121695-01 | Cc1ccc(-c2cn3c(n2)sc2cc(C(=O)NCCC4=CCCCC4)ccc23)cc1 | NaN | 4 | 0 | -0.651 | NaN | 4 | 0 | -0.336 |
| NCGC00123901-01 | O=C(Cn1c(-c2cccs2)cc2ccccc21)N1CCN(c2ccccc2F)CC1 | NaN | 4 | 0 | 6.012 | NaN | 4 | 0 | 4.766 |
| NCGC00120070-01 | CCC1CCCCN1CCCNC(=O)c1[nH]c2ccccc2c1Sc1ccc(Cl)cc1 | 4.466836 | 3 | 41.676 | -4.377 | 14.125375 | -3 | -111.749 | -98.617 |
| NCGC00123907-01 | Cc1ccc(Cl)cc1N1CCN(C(=O)Cn2c(-c3cccs3)cc3cc(F)ccc32)CC1 | NaN | 4 | 0 | 1.318 | NaN | 4 | 0 | -0.647 |
| NCGC00126383-01 | CCCCN(C)C(=O)Cn1ncc2c([nH]c3ccc(C)cc32)c1=O | NaN | 4 | 0 | -0.469 | NaN | 4 | 0 | 2.24 |
| NCGC00126660-01 | O=C(NCC1CCN(Cc2cccc(Cl)c2)CC1)c1cc2ccc3cccnc3c2[nH]1 | NaN | 4 | 0 | 0.409 | NaN | 4 | 0 | -0.717 |
| NCGC00123876-01 | Cc1ccccc1CSc1nc2cccnc2n1Cc1ccc(C(=O)NC2CC2)cc1 | NaN | 4 | 0 | -0.772 | NaN | 4 | 0 | -5.418 |
| NCGC00126455-01 | Cc1ccc(N2CCN(C(=O)Cn3ncc4c([nH]c5ccc(C)cc54)c3=O)CC2)c(C)c1 | NaN | 4 | 0 | 0.833 | NaN | 4 | 0 | 0.576 |
| NCGC00118095-01 | CC(Sc1nc(-c2ccccc2)c(C#N)c(=O)[nH]1)C(=O)Nc1ccc(F)cc1 | NaN | 4 | 0 | 7.709 | NaN | 4 | 0 | -2.371 |
| NCGC00130540-01 | CCCCC(=O)Nc1ccc(C(=O)OCc2cc(=O)n3nc(C4CC4)sc3n2)cc1 | NaN | 4 | 0 | -1.439 | NaN | 4 | 0 | -0.947 |
| NCGC00130453-01 | CCCNC(=O)Cn1nc(-c2ccc(F)cc2)c2cnc3ccc(F)cc3c21 | NaN | 4 | 0 | 1.408 | NaN | 4 | 0 | 1.208 |
| NCGC00130341-01 | O=C(C1CCC(Cn2c(=O)c3sccc3n(Cc3ccc(F)cc3)c2=O)CC1)N1CCOCC1 | NaN | 4 | 0 | -0.288 | NaN | 4 | 0 | -0.406 |
| NCGC00128112-01 | COc1ccc(-c2onc(C)c2C)cc1[S+](=O)([O-])N1CCN(c2ccc(Cl)cc2)CC1 | NaN | 4 | 0 | 1.833 | NaN | 4 | 0 | -0.917 |
| NCGC00126820-01 | CC(C)(C)OC(=O)N1CCC(c2c(C(=O)NCCN3CCCC3)cnn2-c2ccc(Cl)cc2)CC1 | 12.589254 | 4 | 13.262 | 10.738 | NaN | 4 | 0 | -1.308 |
| NCGC00130542-01 | CCC(CC)C(=O)Nc1ccc(C(=O)OCc2cc(=O)n3nc(C4CC4)sc3n2)cc1 | NaN | 4 | 0 | -1.499 | NaN | 4 | 0 | 3.383 |
| NCGC00133323-01 | COc1ccc(-c2nnc(SCC(=O)N3CCN(c4ccc(F)cc4)CC3)[nH]2)cc1 | NaN | 4 | 0 | -0.227 | NaN | 4 | 0 | 1.047 |
| NCGC00125259-01 | Cc1nc2cccnc2n1-c1cccc(C(=O)N2CCN(c3cccc(Cl)c3)CC2)c1 | 14.125375 | 4 | 11 | 10.828 | NaN | 4 | 0 | 2.21 |
| NCGC00124142-01 | Cc1cccc2sc(N(CCCN(C)C)C(=O)COc3ccccc3)nc12 | 7.943282 | 4 | 10.568 | 9.041 | NaN | 4 | 0 | -2.721 |
| NCGC00130944-01 | Cc1ccc(-c2nc(CNC(=O)NC3CCCCC3)c(C)o2)cc1 | NaN | 4 | 0 | 3.559 | NaN | 4 | 0 | 0.246 |
| NCGC00133669-01 | COc1ccccc1Cc1nc2ccccc2nc1SCC(=O)NCCc1ccccc1 | NaN | 4 | 0 | -0.348 | NaN | 4 | 0 | 1.228 |
| NCGC00133727-01 | COc1cccc(Cc2nc3ccccc3nc2SCC(=O)NCCc2ccccc2)c1 | NaN | 4 | 0 | -2.014 | NaN | 4 | 0 | 2.28 |
| NCGC00133737-01 | COc1cccc(Cc2nc3ccccc3nc2SCC(=O)Nc2cccc(C)c2)c1 | NaN | 4 | 0 | 0.651 | NaN | 4 | 0 | 0.576 |
| NCGC00125277-01 | Cc1cccc(N2CCN(C(=O)c3cccc(-n4c(C)nc5cccnc54)c3)CC2)c1C | NaN | 4 | 0 | 0.106 | NaN | 4 | 0 | 2.26 |
| NCGC00133671-01 | COc1ccccc1Cc1nc2ccccc2nc1SCC(=O)NCc1ccc(C)cc1 | NaN | 4 | 0 | 1.378 | NaN | 4 | 0 | -7.362 |
| NCGC00133347-01 | COCCn1c(C)cc(O)c(C(c2ccc(Cl)cc2)N2CCN(c3ccccc3)CC2)c1=O | NaN | 4 | 0 | 4.528 | NaN | 4 | 0 | -2.521 |
| NCGC00134485-01 | CCc1ccccc1NC(=O)C1CCN(c2nc3ccccc3[nH]2)CC1 | NaN | 4 | 0 | 4.074 | NaN | 4 | 0 | -1.98 |
| NCGC00134266-01 | CCC(Oc1ccccc1)C(=O)Nc1cc(-c2ccccc2F)nn1-c1ccccc1 | NaN | 4 | 0 | -0.348 | NaN | 4 | 0 | -2.11 |
| NCGC00136986-01 | O=C(CSc1nc(=O)[nH]c2c1CCCC2)Nc1cc(C(F)(F)F)ccc1Cl | NaN | 4 | 0 | 0.591 | NaN | 4 | 0 | -0.887 |
| NCGC00135408-01 | Cc1ccc(-c2cn3cccnc3n2)cc1NC(=O)C(C)(C)Oc1ccc(Cl)cc1 | NaN | 4 | 0 | 1.166 | NaN | 4 | 0 | -0.877 |
| NCGC00135890-01 | N#Cc1ccccc1NC(=O)C(=O)NCCc1csc(-c2ccc(C(F)(F)F)cc2)n1 | NaN | 4 | 0 | 1.287 | NaN | 4 | 0 | -1.418 |
| NCGC00134593-01 | CCOc1cccc(CC(=O)N2CCc3c([nH]c4ccccc34)C2c2ccc(C)cc2)c1OCC | 14.125375 | 4 | 24.819 | 20.612 | NaN | 4 | 0 | 4.235 |
| NCGC00137666-01 | Cc1cccc(Cn2c(CN3CCC(C(=O)NCc4ccccc4Cl)CC3)cc3ccccc32)c1 | NaN | 4 | 0 | 6.891 | NaN | 4 | 0 | -3.383 |
| NCGC00135953-01 | Cc1cccc(-c2nc(C)c(CCNC(=O)C(=O)Nc3c(C)cc(C)cc3C)s2)c1 | NaN | 4 | 0 | -0.712 | NaN | 4 | 0 | 0.185 |
| NCGC00137951-01 | CCc1cccc(C)c1NC(=O)c1nc2ccccc2nc1N1CCCCC1 | NaN | 4 | 0 | 0.379 | NaN | 4 | 0 | -3.894 |
| NCGC00138672-01 | Cc1ccc(CNC(=O)CSc2nc3ccsc3c(=O)n2-c2ccc(C(=O)O)cc2)cc1 | NaN | 4 | 0 | 0.803 | NaN | 4 | 0 | 2.401 |
| NCGC00131763-01 | Cc1ccc(C)c(Cn2c(SCC(=O)NCc3ccccc3)nc3ccccc32)c1 | NaN | 4 | 0 | 0.984 | NaN | 4 | 0 | -2.661 |
| NCGC00138700-01 | Clc1cccc(CSc2nc(NCCc3ccccc3)c3ccccc3n2)c1 | NaN | 4 | 0 | -0.076 | NaN | 4 | 0 | 1.99 |
| NCGC00120136-01 | COc1ccc2c(Sc3ccccc3)c(C(=O)NCc3cccc(OC)c3OC)[nH]c2c1 | NaN | 4 | 0 | -0.984 | NaN | 4 | 0 | 0.216 |
| NCGC00132023-01 | CCn1ccc2cc([S+](=O)([O-])N3CCC(C(=O)NCC(C)c4ccccc4)CC3)ccc21 | NaN | 4 | 0 | 0.53 | NaN | 4 | 0 | 1.96 |
| NCGC00119299-01 | CCN(CC)C(=O)Cn1c(SCCOc2ccc(Cl)cc2)nc2ccccc21 | NaN | 4 | 0 | -1.62 | NaN | 4 | 0 | 0.095 |
| NCGC00138730-01 | COc1ccc(CCNc2nc(SCc3ccc(Cl)cc3)nc3ccccc23)cc1OC | NaN | 4 | 0 | 6.376 | NaN | 4 | 0 | 4.185 |
| NCGC00138728-01 | COc1ccc(CCNc2nc(SCc3ccccc3Cl)nc3ccccc23)cc1OC | NaN | 4 | 0 | -0.651 | NaN | 4 | 0 | 0.647 |
| NCGC00139256-01 | Cc1ccc(-n2cnc3cc(C(=O)NCCC4=CCCCC4)ccc32)cc1Cl | NaN | 4 | 0 | -0.924 | NaN | 4 | 0 | 1.238 |
| NCGC00138704-01 | COc1ccc(C(C)=O)cc1CSc1nc(NCCc2ccccc2)c2ccccc2n1 | NaN | 4 | 0 | -0.439 | NaN | 4 | 0 | 3.593 |
| NCGC00137700-01 | COc1cccc(CNC(=O)C2CCN(Cc3cc4ccccc4n3Cc3ccc(C)cc3)CC2)c1 | 14.125375 | 4 | 23.031 | 18.885 | NaN | 4 | 0 | 1.899 |
| NCGC00139038-01 | COc1cc2c(cc1OC)C(COc1ccc(F)cc1)N(C(=O)Cc1ccccc1)CC2 | NaN | 4 | 0 | 4.044 | NaN | 4 | 0 | -4.165 |
| NCGC00139437-01 | CCc1ccccc1NC(=O)c1ccc(F)c([S+](=O)([O-])N2CCN(c3cccc(Cl)c3)CC2)c1 | NaN | 4 | 0 | -0.379 | NaN | 4 | 0 | 0.847 |
| NCGC00139646-01 | CCOc1ccc(-c2cc(C(=O)NCCc3ccc(Cl)cc3)c3ccccc3n2)cc1 | NaN | 4 | 0 | -0.621 | NaN | 4 | 0 | -2.17 |
| NCGC00139548-01 | Cc1ccc(C)c(N2CCN([S+](=O)([O-])c3cc(C(=O)Nc4cc(Cl)cc(Cl)c4)ccc3F)CC2)c1 | NaN | 4 | 0 | -1.863 | NaN | 4 | 0 | -0.326 |
| NCGC00139664-01 | COc1ccc(CCNC(=O)c2cc(Sc3cccc(Cl)c3)nc3ccccc23)cc1OC | NaN | 4 | 0 | 0.439 | NaN | 4 | 0 | -4.105 |
| NCGC00138907-01 | COc1ccc(CCNc2ncnc3c2c(-c2ccccc2)cn3-c2ccc(Cl)cc2)cc1OC | NaN | 4 | 0 | -0.197 | NaN | 4 | 0 | -8.034 |
| NCGC00140128-01 | CCOCCCNC(=O)c1cc(Sc2cccc(Cl)c2)nc2ccccc12 | NaN | 4 | 0 | -0.742 | NaN | 4 | 0 | -3.664 |
| NCGC00139776-01 | CCOc1ccccc1CNC(=O)c1cc(-c2ccc(C)c(C)c2)nc2ccccc12 | NaN | 4 | 0 | 0.439 | NaN | 4 | 0 | -3.243 |
| NCGC00139679-01 | Cc1ccc(N2CCN([S+](=O)([O-])c3cc(C(=O)Nc4cc(Cl)cc(Cl)c4)ccc3F)CC2C)cc1 | 12.589254 | 4 | 14.959 | 13.524 | NaN | 4 | 0 | -1.759 |
| NCGC00140138-01 | O=C(CSCc1ccc(F)cc1)N1CCN(c2ccc(Cl)cc2)CC1 | NaN | 4 | 0 | 4.771 | NaN | 4 | 0 | -5.819 |
| NCGC00132017-01 | CCn1ccc2cc([S+](=O)([O-])N3CCC(C(=O)NCCc4ccc(C)cc4)CC3)ccc21 | NaN | 4 | 0 | 0.984 | NaN | 4 | 0 | 3.112 |
| NCGC00140186-01 | CCc1ccc(Sc2cc(C(=O)NCCCOC(C)C)c3ccccc3n2)cc1 | NaN | 4 | 0 | -2.166 | NaN | 4 | 0 | -1.499 |
| NCGC00140224-01 | CCOc1ccc(Sc2cc(C(=O)NCCCOC)c3ccccc3n2)cc1 | NaN | 4 | 0 | -2.257 | NaN | 4 | 0 | -4.024 |
| NCGC00140194-01 | CCc1ccc(Sc2cc(C(=O)NCc3cccc(OC)c3)c3ccccc3n2)cc1 | NaN | 4 | 0 | -1.53 | NaN | 4 | 0 | -3.343 |
| NCGC00139962-01 | Cc1ccc(CCNC(=O)c2ccc(CSCc3ccc(C)cc3)o2)cc1 | NaN | 4 | 0 | -0.257 | NaN | 4 | 0 | 2.681 |
| NCGC00140329-02 | COc1ccccc1Nc1cc(C(=O)NCCCN(C)c2ccccc2)c2ccccc2n1 | NaN | 4 | 0 | 5.073 | NaN | 4 | 0 | -2.661 |
| NCGC00141864-01 | Fc1cccc(F)c1-c1nc(-c2ccncc2)no1 | NaN | 4 | 0 | 0.257 | NaN | 4 | 0 | -0.005 |
| NCGC00141619-01 | Cc1ccc(NC(=O)N2CC=C(c3c[nH]c4ccccc34)CC2)cc1Cl | 14.125375 | 4 | 19.954 | 17.613 | NaN | 4 | 0 | -3.423 |
| NCGC00140469-01 | Cc1ccc(CCNC(=O)c2cc(Nc3ccc(C)c(C)c3)nc3ccccc23)cc1 | NaN | 4 | 0 | 2.59 | NaN | 4 | 0 | 3.303 |
| NCGC00375254-01 | O=C(NCc1cccnc1)c1cc(-c2cnccc2Cl)c2cccn2n1 | NaN | 4 | 0 | 0.015 | NaN | 4 | 0 | -0.005 |
| NCGC00164987-01 | O=C(CSc1cn(CCNC(=O)c2c(F)cccc2F)c2ccccc12)Nc1ccc2c(c1)OCCO2 | NaN | 4 | 0 | -0.136 | NaN | 4 | 0 | -2.1 |
| NCGC00164399-01 | CC/C(=C(\c1ccc(O)cc1)c1ccc(OCCN(C)C)cc1)c1ccccc1 | 5.011872 | 4 | -6.797 | -4.831 | 7.943282 | -2.1 | -108.825 | -98.657 |
| NCGC00377235-01 | Cc1ccc2nc(C3CCN(Cc4ccc(-c5cccs5)cc4)CC3)[nH]c(=O)c2c1 | 12.589254 | 4 | 27.202 | 20.279 | NaN | 4 | 0 | -0.035 |
| NCGC00391370-01 | COC(=O)[C@@H]1Cc2ncn(Cc3ccc(Cl)cc3)c2CN1CCCc1ccccc1 | NaN | 4 | 0 | 3.589 | 1.995262 | -1.2 | -48.043 | -43.968 |
| NCGC00393657-01 | Fc1ccc(-c2c[nH]c([C@@H]3COCCN3Cc3c[nH]c4ccccc34)n2)c(F)c1 | NaN | 4 | 0 | 3.408 | NaN | 4 | 0 | 0.496 |
| NCGC00391516-01 | c1coc(-c2ccnc(N[C@H]3CO[C@@H]4[C@@H](NCC5CCCCC5)CO[C@H]34)n2)c1 | NaN | 4 | 0 | 0.53 | NaN | 4 | 0 | -0.326 |
| NCGC00391867-01 | S=C(Nc1ccccc1)N[C@H]1CO[C@@H]2[C@@H](Nc3nccc(-c4ccco4)n3)CO[C@H]12 | NaN | 4 | 0 | 0.167 | NaN | 4 | 0 | 3.233 |
| NCGC00140462-01 | Cc1ccc(C2CCN(C(=O)Nc3ccc(Cl)cc3)C2)cc1 | NaN | 4 | 0 | 4.134 | NaN | 4 | 0 | 0.526 |
| NCGC00398918-01 | O=C(O)COCC(=O)N1C[C@@H]2C[C@H](C1)c1ccc(-c3cc4ccccc4s3)c(=O)n1C2 | NaN | 4 | 0 | -0.5 | NaN | 4 | 0 | 0.256 |
| NCGC00399719-01 | Cc1nc(-c2cccs2)cc([C@H]2CN3CC[C@H]2C[C@@H]3CNC(=O)C2CC2)n1 | NaN | 4 | 0 | 1.287 | NaN | 4 | 0 | -4.285 |
| NCGC00395987-01 | Clc1ccc(-c2c[nH]c([C@@H]3COCCN3C3CCNCC3)n2)s1 | NaN | 4 | 0 | 0.015 | NaN | 4 | 0 | 1.96 |
| NCGC00391410-01 | O=C(NC[C@H]1OC[C@@H](N(Cc2ccccn2)Cc2ccccn2)[C@@H]1O)C1CCCC1 | NaN | 4 | 0 | 8.526 | NaN | 4 | 0 | -2.601 |
| NCGC00141941-02 | CCN(CC)CCNC(=O)c1ccc2c(c1)NC(=O)C(C[S+](=O)([O-])Cc1ccc(F)cc1Cl)N2 | NaN | 4 | 0 | 5.588 | NaN | 4 | 0 | 0.717 |
| NCGC00140273-01 | CC(C)CN(CCCNC(=O)CN1N=C(c2ccc(Cl)cc2)CCC1=O)CC(C)C | NaN | 4 | 0 | 0.833 | NaN | 4 | 0 | -1.579 |
| NCGC00412600-01 | COc1ccc(-c2ccc3ncc(Nc4ccc(F)c(Cl)c4)n3n2)cn1 | NaN | 4 | 0 | 9.132 | NaN | 4 | 0 | -7.453 |
| NCGC00417461-01 | CC(=O)Nc1cccc(-c2nc(-c3c(C)noc3C)cc3cc(CO)oc23)c1 | NaN | 4 | 0 | -1.015 | NaN | 4 | 0 | -5.749 |
| NCGC00411575-01 | Cc1noc2c(-c3cccnc3NC3CCN(C)CC3)cc(-c3ccc(Cl)cc3)nc12 | 5.011872 | 1.4 | 79.033 | 62.865 | 22.387211 | -3 | -66.846 | -51.686 |
| NCGC00422131-01 | Cn1nc(N2CCC(NC(=O)c3ccncn3)CC2)ccc1=O | NaN | 4 | 0 | 1.166 | NaN | 4 | 0 | 2.09 |
| NCGC00423266-01 | OC(c1cc2n(n1)CCN(c1ncc(C(F)(F)F)cc1Cl)C2)C1CC1 | NaN | 4 | 0 | 1.742 | 2.818383 | 4 | 17.2 | 5.187 |
| NCGC00424096-01 | O=C(NC1CCCc2c1[nH]c1ccc(F)cc21)c1n[nH]c2c1CCCC2 | NaN | 4 | 0 | -1.893 | NaN | 4 | 0 | -0.476 |
| NCGC00420940-01 | COC(=O)[C@H](NC(=O)[C@@H]1C[C@H](NC(=O)Nc2cccc(SC)c2)CN1C(=O)CSC)C(C)C | NaN | 4 | 0 | -0.015 | NaN | 4 | 0 | 0.877 |
| NCGC00424049-01 | Cc1ccc(-c2ccc(CCC(=O)N[C@@H]3CCOC[C@H]3OCC(=O)O)o2)cc1 | NaN | 4 | 0 | -0.439 | NaN | 4 | 0 | 0.827 |
| NCGC00417200-01 | Cc1noc(C)c1-c1cc2c(cc(C)n2CCN(C)C)c(-c2ccccc2O)n1 | 5.011872 | 3 | 49.522 | 1.742 | NaN | 4 | 0 | -8.756 |
| NCGC00401203-01 | COc1ccc(C[C@H](NC(=O)[C@]2(O)C[C@H](NC(=O)CC(C)(C)C)[C@@H](O)[C@H](O)C2)C(N)=O)cc1 | NaN | 4 | 0 | -0.621 | NaN | 4 | 0 | 1.589 |
| NCGC00401616-01 | Cc1ccc(-n2nc(C)c([C@@H]3C=C[C@@H](NC(=O)COCC(=O)O)C3)c2C)cc1Cl | NaN | 4 | 0 | 0.348 | NaN | 4 | 0 | 3.132 |
| NCGC00427991-01 | Cc1oc(-c2ccc(C(F)(F)F)cc2)nc1CN(C)C(C)c1cccs1 | NaN | 4 | 0 | -1.106 | NaN | 4 | 0 | -2.812 |
| NCGC00426773-01 | Cc1cccc(CN2CCN(Cc3nc(-c4cccs4)oc3C)CC2CCO)n1 | NaN | 4 | 0 | -0.015 | NaN | 4 | 0 | -1.288 |
| NCGC00429385-01 | CCC(NC(=O)c1cc(C(C)C)nn1C)c1ccc([S+](C)(=O)[O-])cc1 | NaN | 4 | 0 | -0.167 | NaN | 4 | 0 | -1.729 |
| NCGC00431814-01 | CNC(=O)[C@@H]1C[C@@H](n2cc(CN(C)C(=O)Nc3ccc(F)cc3)nn2)CN1 | NaN | 4 | 0 | 1.075 | NaN | 4 | 0 | -0.767 |
| NCGC00431589-01 | COc1cccc(-c2[nH]cnc2-c2ccccc2)c1OCCCN(C)C | NaN | 4 | 0 | -1.287 | NaN | 4 | 0 | -2.431 |
| NCGC00427701-01 | CN(C(=O)c1ccc2cccc(F)c2n1)C1CCCN(CCCc2ccccc2)C1 | NaN | 4 | 0 | 0.712 | NaN | 4 | 0 | -1.308 |
| NCGC00428272-01 | CC(C)(C)c1nnc(NC(=O)NCc2ccncc2)s1 | NaN | 4 | 0 | -0.439 | NaN | 4 | 0 | -1.93 |
| NCGC00433698-01 | CCSc1nnc(-c2ccccc2NC(=O)C(CC)n2cccn2)c(=O)[nH]1 | NaN | 4 | 0 | 0.015 | NaN | 4 | 0 | 1.809 |
| NCGC00435116-01 | O=C(COc1ccc(NC(=O)C2CC2)cn1)NCCc1cccs1 | NaN | 4 | 0 | 1.651 | NaN | 4 | 0 | 0.236 |
| NCGC00433277-01 | COc1ccc(NC(=O)N2CCCC(Cn3c(C4CC4)nc4cccnc43)C2)cc1Cl | NaN | 4 | 0 | 0.288 | NaN | 4 | 0 | 5.177 |
| NCGC00433509-01 | COc1cc(Cl)cc(CN2CCC3(CCCNC3)C2)c1OC | 5.623413 | 4 | 15.101 | 15.129 | NaN | 4 | 0 | 0.667 |
| NCGC00434789-01 | CC(C)(O)C#Cc1ccc(-c2nccn2-c2cccc(-n3cncn3)c2)cc1 | NaN | 4 | 0 | -1.62 | 3.548134 | 4 | 18.027 | -1.208 |
| NCGC00436130-01 | Cc1cccc(C(=O)N2CCCN(c3cc(-c4ccc(Cl)cc4)n[nH]3)CC2)c1 | NaN | 4 | 0 | 3.922 | NaN | 4 | 0 | -5.297 |
| NCGC00437455-01 | COc1ccccc1-n1cc(CNC(=O)C(N)c2c(C)n[nH]c2C)cn1 | NaN | 4 | 0 | 0.833 | NaN | 4 | 0 | -0.035 |
| NCGC00437247-01 | Cc1csc2c(NCCC3CN(c4cnn(C)c(=O)c4)CCO3)ncnc12 | NaN | 4 | 0 | -0.318 | NaN | 4 | 0 | -6.811 |
| NCGC00437347-01 | Cc1c(-c2ccccc2F)[nH]c2ccc(CNC(=O)CCc3cc(O)no3)cc12 | NaN | 4 | 0 | -0.712 | 1.412538 | -1.2 | -49.823 | -49.632 |
| NCGC00435587-01 | CNC(=O)C1(NC(=O)c2ccc(CCC(C)(C)O)cc2)CCCCC1 | NaN | 4 | 0 | 0.348 | NaN | 4 | 0 | 1.879 |
| NCGC00431486-01 | O=C(Cn1ccn2nc(-c3ccc(F)cc3)c(CO)c2c1=O)N1CCN(c2ccccc2F)CC1 | NaN | 4 | 0 | -0.318 | NaN | 4 | 0 | -0.777 |
| NCGC00428095-01 | NCC1CCCN(Cc2ccc(-c3nc(-c4ccncc4)cc(=O)[nH]3)cc2)C1 | NaN | 4 | 0 | 2.196 | NaN | 4 | 0 | 1.027 |
| NCGC00438285-01 | Cc1n[nH]c(C)c1CCCNC(=O)C1CC2(CCNCC2)CN1 | NaN | 4 | 0 | 0.682 | NaN | 4 | 0 | -0.446 |
| NCGC00439147-01 | O=C(c1ncoc1C1CC1)N1CCOCC(O)(CN2CCCC2)C1 | NaN | 4 | 0 | 2.529 | NaN | 4 | 0 | -0.216 |
| NCGC00439810-01 | CN(Cc1nc2ccsc2c(=O)[nH]1)C(=O)CCC1NC(=O)NC1=O | NaN | 4 | 0 | 0.924 | NaN | 4 | 0 | -0.677 |
| NCGC00438711-01 | CC(=O)N[C@@H]1CN(C(=O)c2csc(-c3cccnc3)n2)C[C@H]1C(C)C | NaN | 4 | 0 | 2.923 | NaN | 4 | 0 | 2.361 |
| NCGC00431062-01 | CCc1nc2c(=O)n(CC(=O)NCc3ccc(F)cc3Cl)nc(C)c2s1 | NaN | 4 | 0 | 0.803 | NaN | 4 | 0 | -1.98 |
| NCGC00440502-01 | FC(F)Oc1ccccc1CN1CCCC12CCN(c1ncccn1)CC2 | NaN | 4 | 0 | 0.651 | NaN | 4 | 0 | -0.937 |
| NCGC00440262-01 | COc1ccc(-c2n[nH]cc2CNC(=O)C2CC3(CCNCC3)CN2)cc1 | NaN | 4 | 0 | -1.378 | NaN | 4 | 0 | 3.403 |
| NCGC00440540-01 | N#Cc1cccnc1N1CCC(NC(=O)c2ccc3cccc(F)c3n2)CC1 | NaN | 4 | 0 | 3.438 | NaN | 4 | 0 | -0.837 |
| NCGC00443121-01 | COc1ccc(OC)c2nc(-c3cccc(Cl)c3)c(CN3CCCC(O)C3)cc12 | NaN | 4 | 0 | -1.984 | NaN | 4 | 0 | -0.717 |
| NCGC00446784-01 | O=C1NCCCc2[nH]c(-c3csc(-c4cnccn4)n3)nc21 | NaN | 4 | 0 | 0.076 | NaN | 4 | 0 | -1.468 |
| NCGC00445152-01 | O=C(Cn1cccc1-c1nc(-c2ccccc2)no1)Nc1ccccc1Cl | NaN | 4 | 0 | 3.529 | NaN | 4 | 0 | -0.657 |
| NCGC00427736-01 | O=C(CCC[C@@H]1[C@H]2CCCN3CCC[C@@H](CN1C(=O)NCc1ccc(F)cc1)[C@@H]23)NCC1CC1 | NaN | 4 | 0 | 2.741 | NaN | 4 | 0 | 4.696 |
| NCGC00449073-01 | O=c1c2cc(-c3cccs3)nn2ccn1CC(O)c1cccc(Cl)c1 | NaN | 4 | 0 | -2.105 | NaN | 4 | 0 | 3.193 |
| NCGC00445041-01 | COCCN(Cc1c(C)nn(C)c1C)C[C@@H]1[C@H]2CNC[C@H]21 | NaN | 4 | 0 | 0.197 | NaN | 4 | 0 | 3.994 |
| NCGC00449414-01 | COCCNc1nc2cc(-c3nc(-c4cccnn4)no3)ccc2n1C | NaN | 4 | 0 | -0.439 | NaN | 4 | 0 | -0.747 |
| NCGC00450105-01 | Fc1ccc(-c2noc(-c3nc(-c4ccncc4)no3)n2)cc1F | NaN | 4 | 0 | -1.015 | NaN | 4 | 0 | 2.16 |
| NCGC00439700-01 | Cc1ccnc(C(NC(=O)CCc2cc3n(n2)CCCNC3)C2CC2)c1 | NaN | 4 | 0 | 1.136 | NaN | 4 | 0 | 0.496 |
| NCGC00450768-01 | CCc1ccccc1NC(=O)CN1C(=O)NC2(CCCCC2)C1=O | NaN | 4 | 0 | 0.5 | NaN | 4 | 0 | 0.216 |
| NCGC00449118-01 | Cc1cccc(-n2ncc3c2CCCC3NC(=O)C2(C)CCNCC2)c1C | NaN | 4 | 0 | 1.075 | NaN | 4 | 0 | -0.977 |
| NCGC00456119-01 | COc1ccccc1OCCNC(=O)C1(C)CCCCC(=O)N1 | NaN | 4 | 0 | 1.893 | NaN | 4 | 0 | 1.198 |
| NCGC00456122-01 | COc1cccc(CNC(=O)C2(C)CCCCC(=O)N2)c1OC | NaN | 4 | 0 | -1.53 | NaN | 4 | 0 | -0.576 |
| NCGC00456156-01 | CCC1(C(=O)NCc2ccc(C)cc2)CCCCC(=O)N1 | NaN | 4 | 0 | 0.288 | NaN | 4 | 0 | -1.278 |
| NCGC00460545-01 | CC(C)N1CC[C@H](CO)[C@H](NC(=O)C2CC2)C1 | NaN | 4 | 0 | -1.408 | 4.466836 | 4 | 14.876 | 0.085 |
| NCGC00464151-01 | Cc1ncsc1CN1CCC(Cc2ncc(C)n2CC2CC2)C1 | 10 | 4 | 10.141 | 8.799 | NaN | 4 | 0 | -0.777 |
| NCGC00418019-01 | CNC(=O)CCn1ncc2cc(OCc3ccccc3-c3cccc(NC(C)=O)c3)ccc21 | NaN | 4 | 0 | 0.924 | NaN | 4 | 0 | -2.701 |
| NCGC00449000-01 | Cc1ncc(-c2ccc(F)c3cccnc23)nc1C | NaN | 4 | 0 | 0.5 | NaN | 4 | 0 | 0.135 |
| NCGC00464002-01 | CC(C)(C)NC(=O)N1CCC2(CC1)CC(Cc1nc(-c3cnccn3)no1)CCO2 | NaN | 4 | 0 | -0.409 | NaN | 4 | 0 | -2.26 |
| NCGC00448942-01 | Cc1ccc(F)c(C(=O)NC(C)c2cnn(-c3ccccc3)c2C)c1Cl | NaN | 4 | 0 | -1.106 | NaN | 4 | 0 | -1.468 |
| NCGC00444561-01 | Cc1c(C(C)NC(=O)c2cc(CC(C)C)nn2C)cnn1-c1ccccc1 | 14.125375 | 4 | 13.137 | 11.707 | 6.309573 | 4 | -15.543 | -14.619 |
| NCGC00465972-01 | COc1cccc(CN2CCCC(c3nc(-c4cnccn4)no3)CCNC(=O)CC2)c1OC | NaN | 4 | 0 | -0.712 | NaN | 4 | 0 | 1.679 |
| NCGC00469713-01 | Cc1cnc(CC2CCCN(C(=O)C3(c4ccccc4)CC3)C2)n1Cc1ccc(F)cc1 | NaN | 4 | 0 | 2.257 | NaN | 4 | 0 | -3.613 |
| NCGC00469744-01 | Cc1cnc(CC2CCN(C(=O)C(C)c3ccccc3)CC2)n1Cc1ccc(F)cc1 | NaN | 4 | 0 | 1.954 | NaN | 4 | 0 | -2.992 |
| NCGC00474800-01 | COc1ccc(C(=O)NC(C(=O)NC2CC2)C2CCN(C(C)=O)CC2)cc1 | NaN | 4 | 0 | 2.075 | NaN | 4 | 0 | 3.553 |
| NCGC00470369-01 | CCN1CCN(CCCNC(=O)c2cn3ccnc(-c4ccc(C)cc4)c3n2)CC1 | NaN | 4 | 0 | 2.045 | NaN | 4 | 0 | -3.914 |
| NCGC00462915-01 | CN(C)C[C@@H]1CN(C2CCOCC2)C[C@H]1NC(=O)c1cc2ccccc2[nH]1 | NaN | 4 | 0 | 1.56 | NaN | 4 | 0 | -3.062 |
| NCGC00470533-01 | O=C(c1ccc2ccccc2n1)N1CCC2(CC1)CC(Cc1cccc(F)c1)CCO2 | NaN | 4 | 0 | -0.833 | NaN | 4 | 0 | 5.378 |
| NCGC00466921-01 | O=C1CCN(C(=O)[C@H](Cc2ccccc2)NC(=O)c2cccs2)CCCOCCN1 | NaN | 4 | 0 | 0.621 | NaN | 4 | 0 | -2.681 |
| NCGC00466138-01 | O=C1NCCCCCCN[C@H]2C[C@H](c3nc(-c4cnccn4)no3)C[C@@H]12 | NaN | 4 | 0 | -0.045 | NaN | 4 | 0 | -1.098 |
| NCGC00482025-01 | O=C(NCCc1ccccc1F)c1cc(-c2cccc(C(F)(F)F)c2)c2nncn2n1 | NaN | 4 | 0 | -2.105 | NaN | 4 | 0 | -0.496 |
| NCGC00476968-01 | COc1cccc([S+](=O)([O-])N2CCc3nc(C4CC4)nc(-c4ccc(Cl)cc4)c3CC2)c1 | NaN | 4 | 0 | -2.014 | NaN | 4 | 0 | 1.559 |
| NCGC00489670-01 | COc1ccc(NC(=O)c2cn3cc(-c4ccc(F)cc4)sc3n2)cc1 | NaN | 4 | 0 | 2.862 | NaN | 4 | 0 | 1.96 |
| NCGC00478196-01 | CCCN1CCC2(CCN(C(=O)c3ccc(C)cc3)CC2)Oc2ccccc21 | NaN | 4 | 0 | -1.378 | NaN | 4 | 0 | -1.108 |
| NCGC00478356-02 | CC(C)CCN1CCC2(CC1)CC(=O)N(c1ccccc1)c1ccccc1O2 | NaN | 4 | 0 | -2.075 | NaN | 4 | 0 | 2.862 |
| NCGC00490602-01 | Cc1ccc(-c2ccccc2CN2CCN(C(=O)Cc3ccc(F)cc3)CC2)cc1Cl | NaN | 4 | 0 | 4.256 | NaN | 4 | 0 | -0.025 |
| NCGC00465936-01 | COc1cccc(CN2CCCC(c3nc(-c4cccnc4)no3)CCNC(=O)CC2)n1 | NaN | 4 | 0 | -0.954 | NaN | 4 | 0 | 0.767 |
| NCGC00492821-01 | O=C(Nc1ccc(Cl)cc1)N1CCN(Cc2ccccc2-c2ccccc2F)CC1 | NaN | 4 | 0 | -0.712 | NaN | 4 | 0 | 2.12 |
| NCGC00490610-01 | Cc1ccc(-c2ccccc2CN2CCN(C(=O)Nc3ccc(Cl)cc3)CC2)cc1Cl | NaN | 4 | 0 | 2.075 | NaN | 4 | 0 | -3.904 |
| NCGC00490613-01 | Cc1ccc([S+](=O)([O-])N2CCN(Cc3ccccc3-c3ccc(C)c(Cl)c3)CC2)cc1 | NaN | 4 | 0 | 0.772 | 0.316228 | 4 | -28.591 | -1.128 |
| NCGC00492929-01 | O=C(NC1CC1)C1CCCCN1Cc1ccc(-c2cccs2)cc1 | NaN | 4 | 0 | -0.591 | NaN | 4 | 0 | -8.124 |
| NCGC00492822-01 | Cc1ccc([S+](=O)([O-])N2CCN(Cc3ccccc3-c3ccccc3F)CC2)cc1 | NaN | 4 | 0 | 5.164 | NaN | 4 | 0 | -4.576 |
| NCGC00493022-01 | O=C(NCCNC(=O)c1cccc(-c2ccsc2)c1)Nc1ccc(Cl)cc1 | NaN | 4 | 0 | 2.075 | NaN | 4 | 0 | 0.496 |
| NCGC00492964-01 | O=C(Cc1ccc(F)cc1)NC1CCN(Cc2cccc(-c3cccs3)c2)CC1 | NaN | 4 | 0 | 1.863 | NaN | 4 | 0 | -0.897 |
| NCGC00492943-01 | O=C(Nc1ccc(F)cc1)C1CCCCN1Cc1ccc(-c2cccs2)cc1 | NaN | 4 | 0 | 2.741 | NaN | 4 | 0 | -4.866 |
| NCGC00493064-01 | COc1cccc(-c2cccc(C(=O)N3CCCC3C(=O)NCCc3ccccc3)c2)c1 | NaN | 4 | 0 | -1.136 | NaN | 4 | 0 | 2.531 |
| NCGC00492974-01 | COc1ccc(CC(=O)NC2CCN(Cc3cccc(-c4cccs4)c3)CC2)cc1 | NaN | 4 | 0 | -2.105 | NaN | 4 | 0 | 0.326 |
| NCGC00493133-01 | O=C(NCc1ccc(F)cc1)C1CCN(Cc2ccccc2-c2cccnc2)CC1 | NaN | 4 | 0 | -0.742 | NaN | 4 | 0 | -0.967 |
| NCGC00493138-01 | O=C(NCc1ccccc1Cl)C1CCN(Cc2ccccc2-c2cccnc2)CC1 | NaN | 4 | 0 | 4.71 | NaN | 4 | 0 | -2.11 |
| NCGC00493137-01 | O=C(NCCc1ccc(Cl)cc1)C1CCN(Cc2ccccc2-c2cccnc2)CC1 | 14.125375 | 4 | 11.959 | 8.284 | NaN | 4 | 0 | -4.656 |
| NCGC00498632-01 | O=C(NCc1cccc(F)c1)C1CCCN(Cc2cccc(-c3ccc(F)cc3F)c2)C1 | NaN | 4 | 0 | 5.679 | 10 | -2.2 | -52.392 | -53.992 |
| NCGC00498631-01 | O=C(NCc1ccc(F)cc1)C1CCCN(Cc2cccc(-c3ccc(F)cc3F)c2)C1 | 12.589254 | 3 | 54.007 | 47.872 | 12.589254 | -3 | -35.909 | -35.077 |
| NCGC00433090-01 | Cc1nnc(CN2CCOc3c(O)cc(-c4ccccc4C)cc3C2)s1 | NaN | 4 | 0 | 0.106 | NaN | 4 | 0 | -2.05 |
| NCGC00498679-01 | COc1ccccc1-c1cccc(CN2CCCC(C(=O)Nc3ccc(F)cc3)C2)c1 | 7.079458 | 2.2 | 58.741 | 51.598 | 12.589254 | -3 | -44.29 | -44.64 |
| NCGC00498681-01 | COc1ccccc1-c1cccc(CN2CCCC(C(=O)Nc3cccc(Cl)c3)C2)c1 | 11.220185 | 4 | 25 | 23.671 | 14.125375 | -3 | -46.667 | -44.52 |
| NCGC00498684-01 | COc1ccccc1-c1cccc(CN2CCCC(C(=O)NCc3cccc(F)c3)C2)c1 | NaN | 4 | 0 | -0.742 | 8.912509 | -2.1 | -85.613 | -86.849 |
| NCGC00498909-01 | O=C(Cc1ccc(F)cc1)NC1CCN(Cc2ccc(-c3ccsc3)cc2)C1 | NaN | 4 | 0 | 5.346 | NaN | 4 | 0 | -4.977 |
| NCGC00498763-01 | Cc1cc(C)cc(-c2ccc(CN3CCCC3C(=O)NCCc3ccccc3)cc2)c1 | 14.125375 | 4 | 15.613 | 13.312 | 14.125375 | 4 | -21.54 | -19.2 |
| NCGC00498972-01 | Cc1ccc(-c2cccc(CN3CCCC(NC(=O)c4ccc(Cl)nc4)C3)c2)cc1Cl | NaN | 4 | 0 | 0.257 | NaN | 4 | 0 | -1.468 |
| NCGC00469712-01 | Cc1cnc(CC2CCCN(C(=O)COc3ccccc3)C2)n1Cc1ccc(F)cc1 | NaN | 4 | 0 | -0.439 | NaN | 4 | 0 | -6.54 |
| NCGC00498779-01 | O=C(NCc1ccc(F)cc1F)C1CCCCN1Cc1ccc(-c2ccsc2)cc1 | NaN | 4 | 0 | -1.802 | NaN | 4 | 0 | -8.074 |
| NCGC00498975-01 | Cc1ccc(-c2cccc(CN3CCCC(NC(=O)c4ccc(C(F)(F)F)cc4)C3)c2)cc1Cl | NaN | 4 | 0 | 0.197 | NaN | 4 | 0 | 1.158 |
| NCGC00498971-01 | Cc1ccc(C(=O)NC2CCCN(Cc3cccc(-c4ccc(C)c(Cl)c4)c3)C2)cc1 | NaN | 4 | 0 | -0.984 | NaN | 4 | 0 | 0.296 |
| NCGC00498982-01 | COc1cccc(C(=O)NC2CCCN(Cc3cccc(-c4ccc(C)c(Cl)c4)c3)C2)c1 | NaN | 4 | 0 | -0.621 | NaN | 4 | 0 | -1.448 |
| NCGC00498988-01 | CCOC(=O)NC1CCCN(Cc2cccc(-c3ccc(C)c(Cl)c3)c2)C1 | NaN | 4 | 0 | 2.378 | NaN | 4 | 0 | -5.307 |
| NCGC00017063-12 | CCN(CC)Cc1cc(Nc2ccnc3cc(Cl)ccc23)ccc1O | 5.623413 | 2.1 | 83.412 | 89.007 | 11.220185 | 4 | -10.768 | -11.211 |
| NCGC00098551-01 | CCn1c2ccccc2c2cc(CN3CCN(Cc4cccc(Cl)c4)CC3)ccc21 | 12.589254 | 3 | 105.909 | 97.042 | NaN | 4 | 0 | 0.058 |
| NCGC00099289-01 | CCOc1ccc(CN2CCN(Cc3ccc4c(c3)c3ccccc3n4CC)CC2)cc1 | 11.220185 | 3 | 68.7 | 65.778 | 14.125375 | 4 | -25.834 | -23.612 |
| NCGC00100643-02 | COc1ccc(Nc2nc(Nc3ccc(OC)cc3OC)nc(N3CCOCC3)n2)c(OC)c1 | 3.162278 | 1.1 | 87.52 | 90.175 | NaN | 4 | 0 | -12.885 |
| NCGC00105925-01 | CCCN(CCC)CCNC(=O)c1cc(-c2ccc(Br)s2)nc2ccccc12 | 10 | 2.2 | 77.801 | 67.093 | NaN | 4 | 0 | -5.383 |
| NCGC00105953-02 | CCCCN(CCCC)CCCNC(=O)c1cc(-c2ccc(Cl)s2)nc2ccccc12 | 6.309573 | 2.2 | 81.257 | 72.462 | NaN | 4 | 0 | -10.165 |
| NCGC00108138-01 | CCCCN(CCCNC(=O)c1cc(Nc2ccc(OC)c(OC)c2)nc2ccccc12)Cc1ccccc1 | 10 | 2.2 | 69.245 | 60.044 | NaN | 4 | 0 | -5.237 |
| NCGC00108196-02 | COc1ccc(Nc2cc(C(=O)NCCCN3CCN(c4ccccc4F)CC3)c3ccccc3n2)c(OC)c1 | 5.623413 | 2.2 | 40.356 | 42.768 | 12.589254 | 4 | -31.476 | -29.566 |
| NCGC00108262-01 | Cc1ccc(C)c(Nc2cc(C(=O)NCCCN(C)C3CCCCC3)c3ccccc3n2)c1 | 6.309573 | 2.2 | 83.393 | 74.58 | 12.589254 | 4 | -29.828 | -27.794 |
| NCGC00108321-02 | CCCCN(CCCNC(=O)c1cc(Nc2cccc(SC)c2)nc2ccccc12)Cc1ccccc1 | 3.548134 | 1.4 | 37.903 | 33.126 | 6.309573 | 4 | -10.269 | -8.974 |
| NCGC00109487-02 | COc1ccc(-c2nc(CN3CCC(C(=O)NCCCN4CCN(c5ccccc5F)CC4)CC3)c(C)o2)cc1OC | 8.912509 | 2.2 | 79.446 | 73.667 | 11.220185 | 4 | -24.31 | -22.76 |
| NCGC00109549-01 | CCCCSCCCNC(=O)C1CCN(Cc2nc(-c3ccc(CC)cc3)oc2C)CC1 | 12.589254 | 3 | 99.013 | 89.847 | 5.623413 | 4 | -8.262 | -5.218 |
| NCGC00109615-01 | CCCCNC(=O)C1CCCN(Cc2nc(-c3ccc(Cl)cc3)oc2C)C1 | 12.589254 | 3 | 51.889 | 48.21 | 7.943282 | 4 | -28.622 | -26.352 |
| NCGC00109633-01 | CCCN(CCC)CCCNC(=O)C1CCCN(Cc2nc(-c3ccc(Cl)cc3)oc2C)C1 | 3.981072 | 1.1 | 87.273 | 85.062 | 11.220185 | 4 | -18.959 | -15.799 |
| NCGC00110541-02 | CCCN1CCN(CCCNC(=O)c2cc3c(Cl)nc4ccccc4c3s2)CC1 | 3.981072 | 1.2 | 81.995 | 78.853 | 5.623413 | 4 | -17.134 | -15.945 |
| NCGC00110841-01 | Cc1ccc2nc(Cl)c3cc(C(=O)NCCCN4CCC(C)CC4)sc3c2c1 | 3.548134 | 1.2 | 70.716 | 61.322 | 10 | 4 | -14.903 | -16.651 |
| NCGC00110901-02 | CCN(CC)CCCNC(=O)CSc1c2c(nc3cc(Cl)ccc13)CCCC2 | 14.125375 | 3 | 95.547 | 84.039 | 2.818383 | 4 | 7.946 | 0.6 |
| NCGC00113431-01 | CCCCN(CCCC)CCCNC(=O)c1ccc2nc(N3CCCCC3)sc2c1 | 11.220185 | 3 | 46.106 | 42.513 | NaN | 4 | 0 | -4.821 |
| NCGC00113433-01 | CCCCN(C)CCCNC(=O)c1ccc2nc(N3CCCCC3)sc2c1 | 10 | 3 | 69.91 | 66.472 | NaN | 4 | 0 | -8.877 |
| NCGC00120066-01 | CCN(CC)CCNC(=O)c1[nH]c2ccccc2c1Sc1ccc(Cl)cc1 | 12.589254 | 3 | 114.77 | 104.529 | NaN | 4 | 0 | -3.175 |
| NCGC00120134-01 | COc1ccc2c(Sc3ccccc3)c(C(=O)NCCCN3CCCCC3)[nH]c2c1 | 12.589254 | 3 | 91.809 | 83.346 | 12.589254 | 4 | -17.112 | -14.26 |
| NCGC00120194-01 | COc1ccc2c(Sc3ccc(Cl)cc3)c(C(=O)NCCCN3CCCCC3)[nH]c2c1 | 12.589254 | 3 | 109.684 | 97.443 | 3.981072 | 4 | 11.507 | -2.004 |
| NCGC00121825-01 | COc1ccc(-c2cn3c(n2)sc2cc(C(=O)NCCCN4CCCCCC4)ccc23)cc1 | 14.125375 | 3 | 78.545 | 68.517 | 10 | 4 | -28.321 | -25.267 |
| NCGC00121953-01 | O=C(NCCCN1CCCCCC1)c1ccc2c(c1)sc1nc(-c3ccccc3)cn12 | 12.589254 | 3 | 86.696 | 76.552 | 7.943282 | -2.4 | -33.262 | -26.052 |
| NCGC00132814-01 | CCN(CC)CCCNC(=O)c1ccc(-c2nc(CN3CCc4ccccc43)c(C)o2)cc1 | 12.589254 | 3 | 127.332 | 116.034 | NaN | 4 | 0 | -3.059 |
| NCGC00132818-01 | CCN(CC)CCNC(=O)c1ccc(-c2nc(CN3CCc4ccccc43)c(C)o2)cc1 | 12.589254 | 3 | 108.22 | 98.101 | NaN | 4 | 0 | -2.856 |
| NCGC00137630-01 | COCCCNC(=O)C1CCN(Cc2cc3ccccc3n2Cc2cccc(Cl)c2)CC1 | 12.589254 | 3 | 50.073 | 44.923 | NaN | 4 | 0 | -9.913 |
| NCGC00137604-01 | COc1cccc(CNC(=O)C2CCN(Cc3cc4ccccc4n3Cc3ccccc3)CC2)c1OC | 12.589254 | 3 | 46.828 | 41.782 | 14.125375 | 4 | -10.285 | -11.453 |
| NCGC00139688-01 | O=C(NCCCN1CCCCC1)c1cc(Sc2ccc(Cl)cc2)nc2ccccc12 | 11.220185 | 2.1 | 108.286 | 98.904 | 0.354813 | 4 | -16.703 | -6.002 |
| NCGC00139692-01 | CC1CCN(CCCNC(=O)c2cc(Sc3ccc(F)cc3)nc3ccccc23)CC1 | 11.220185 | 3 | 97.629 | 92.768 | NaN | 4 | 0 | -9.265 |
| NCGC00140152-01 | CC1CCCCN1CCCNC(=O)c1cc(Sc2cccc(Cl)c2)nc2ccccc12 | 10 | 2.1 | 108.557 | 97.991 | 12.589254 | 4 | -11.597 | -11.782 |
| NCGC00140328-01 | COc1ccc(Nc2cc(C(=O)NCCCN(C)Cc3ccccc3)c3ccccc3n2)cc1 | 11.220185 | 3 | 73.855 | 65.888 | 8.912509 | 4 | -9.518 | -14.105 |
| NCGC00377200-01 | Cc1ccc2nc(C3CCCN(Cc4cccc(-c5cccnc5)c4)C3)[nH]c(=O)c2c1 | 12.589254 | 3 | 82.666 | 74.872 | NaN | 4 | 0 | -1.413 |
| NCGC00379304-01 | CC(Nc1nc(N2CCN(C)CC2)nc2ccccc12)c1ccccc1 | 12.589254 | 3 | 119.723 | 109.277 | NaN | 4 | 0 | 0.252 |
| NCGC00384291-01 | CNCCN(C)c1nc(NC(C)c2ccccc2)c2ccccc2n1 | 10 | 2.1 | 116.742 | 101.899 | NaN | 4 | 0 | -1.927 |
| NCGC00178968-01 | CCN(CC)Cc1ccc(Nc2ccnc3cc(Cl)ccc23)cc1O | 8.912509 | 2.1 | 97.152 | 92.221 | NaN | 4 | 0 | -9.139 |
| NCGC00140589-02 | COc1ccc(N2CCN(CCCNC(=O)c3cc(Nc4ccc(OC)cc4OC)nc4ccccc34)CC2)cc1 | 11.220185 | 3 | 80.089 | 73.63 | 12.589254 | 4 | -14.696 | -13.631 |
| NCGC00377241-01 | Cc1ccc(-c2cccc(CN3CCCC(c4nc5ccc(C)cc5c(=O)[nH]4)C3)c2)c(C)c1 | 10 | 2.1 | 111.277 | 100.913 | NaN | 4 | 0 | -8.471 |
| NCGC00377240-01 | COc1cccc(-c2ccc(CN3CCCC(c4nc5ccc(C)cc5c(=O)[nH]4)C3)cc2)c1 | 4.466836 | 1.1 | 99.414 | 100.256 | NaN | 4 | 0 | 6.157 |
| NCGC00140661-02 | COc1ccc(Nc2cc(C(=O)NCCCN3CCN(c4ccc(OC)cc4)CC3)c3ccccc3n2)cc1 | 12.589254 | 3 | 46.812 | 46.457 | 10 | 4 | -20.596 | -21.24 |
| NCGC00390036-01 | C[C@@H](Nc1nc(N2CCNCC2)nc2ccccc12)c1ccccc1 | 11.220185 | 3 | 101.554 | 97.589 | NaN | 4 | 0 | -2.565 |
| NCGC00390104-01 | Cc1ccc(C(C)Nc2nc(N3CCNCC3)nc3ccccc23)cc1 | 5.011872 | 1.1 | 90.132 | 89.408 | NaN | 4 | 0 | -3.94 |
| NCGC00389098-01 | CC(Nc1nc(N2CCNCC2C)nc2ccccc12)c1ccccc1 | 8.912509 | 2.1 | 112.168 | 98.393 | NaN | 4 | 0 | -9.081 |
| NCGC00400702-01 | O=C(C[C@@H]1CCNC[C@@H]1Cc1cc(-c2ccc(F)cc2)on1)N1CCC(Cc2ccccc2)CC1 | 12.589254 | 3 | 100.317 | 91.709 | NaN | 4 | 0 | -1.559 |
| NCGC00419475-01 | CN(C)CCNC(=O)c1ccc2c(c1)C1(CCN(CC3CC3)C1)CN2Cc1ccccc1 | 5.011872 | 1.1 | 96.941 | 94.302 | NaN | 4 | 0 | -3.398 |
| NCGC00419419-01 | CN1CCCC2(C1)CN(Cc1cccnc1)c1ccc(C(=O)NCC3CCCCC3)cc12 | 14.125375 | 3 | 89.028 | 91.673 | 19.952623 | 4 | 18.683 | 10.552 |
| NCGC00427036-01 | O=C(NCCN1CCCCC1)c1ccc(-c2ccc(N3CCSCC3)nc2)cc1 | 12.589254 | 3 | 57.448 | 52.739 | 14.125375 | 4 | -24.459 | -24.067 |
| NCGC00437792-01 | CN(C)c1cccc(C(=O)NCC2CCN(c3cc(-c4ccc(Cl)cc4)n[nH]3)CC2)c1 | 12.589254 | 3 | 74.363 | 68.736 | 14.125375 | 4 | -27.995 | -25.413 |
| NCGC00449240-01 | Clc1ccc2nc(N3CCC(NCCc4cccnc4)CC3)sc2c1 | 14.125375 | 3 | 101.112 | 89.554 | 8.912509 | 4 | -15.257 | -14.018 |
| NCGC00479266-01 | Cc1ccc([C@@H](C)Nc2nc(N3CCNCC3)nc3ccccc23)cc1 | 4.466836 | 1.1 | 91.278 | 89.883 | NaN | 4 | 0 | -9.362 |
| NCGC00480765-01 | CC(C)(Nc1nc(N2CCNCC2)nc2ccccc12)c1ccccc1 | 11.220185 | 3 | 52.546 | 50.584 | 5.623413 | 4 | -10.856 | -4.047 |
| NCGC00487028-01 | CC(Nc1nc(N2CCN(CCN)CC2)nc2ccccc12)c1ccccc1 | 5.623413 | 2.2 | 52.154 | 56.465 | 14.125375 | 4 | -12.702 | -12.372 |
| NCGC00479261-01 | C[C@@H](Nc1nc(N2CCNCC2)nc2ccccc12)c1ccc(F)cc1 | 10 | 2.1 | 93.718 | 87.217 | 3.981072 | 4 | -8.942 | -9.952 |
| NCGC00489837-01 | Cc1ncccc1-c1nc2cc(C(=O)NCCCc3ccccc3)ccc2o1 | 15.848932 | 3 | 45.211 | 48.21 | NaN | 4 | 0 | -10.785 |
| NCGC00492858-01 | Cc1ccc(Nc2ccnc(N3CCC(C(=O)NCCCc4ccccc4)CC3)n2)cc1 | 7.079458 | 2.2 | 53.961 | 49.963 | 11.220185 | -2.4 | -30.352 | -29.111 |
| NCGC00496752-01 | Cc1ccc2c(NC(C)c3ccccc3)nc(N3CCNCC3)nc2c1 | 3.981072 | 1.1 | 92.845 | 92.549 | NaN | 4 | 0 | -9.458 |
| NCGC00496256-01 | O=C(CCc1ccc(CC2CCN(Cc3c[nH]c4ccccc34)CC2)cc1)N1CCCC1 | 12.589254 | 3 | 36.194 | 31.848 | NaN | 4 | 0 | -11.685 |
| NCGC00522637-01 | CC(C)(C)NCc1ccc(Nc2ccnc3cc(Cl)ccc23)cc1O | 3.162278 | 1.3 | 85.147 | 75.53 | 11.220185 | 4 | -17.765 | -19.507 |
| NCGC00098575-02 | CCn1c2ccccc2c2cc(CN3CCN(Cc4ccc(Cl)cc4)CC3)ccc21 | 4.466836 | 3 | 60.607 | 24.507 | 14.125375 | -3 | -79.476 | -72.453 |
| NCGC00101043-02 | CCn1cc(CNC(=O)c2cc(-c3ccc(C)cc3)nc3ccc(Br)cc23)c(C)n1 | NaN | 4 | 0 | 0.037 | NaN | 4 | 0 | -1.046 |
| NCGC00103969-01 | O=C(Nc1ccc(F)c(Cl)c1)c1ccc2c(Cl)c3c(nc2c1)CCCC3 | NaN | 4 | 0 | -0.621 | NaN | 4 | 0 | -1.084 |
| NCGC00094585-01 | CCN(CC)Cc1ccc(Nc2ccnc3cc(Cl)ccc23)cc1O | NaN | 4 | 0 | 0.073 | 10 | 4 | 13.178 | 11.278 |
| NCGC00102900-01 | CCOc1ccc(Cn2c(-c3ccc(OCC)c(OC)c3)nc3ccccc32)cc1OC | NaN | 4 | 0 | 2.447 | NaN | 4 | 0 | -0.639 |
| NCGC00104256-01 | CCN(CC)c1cc(C)c2cc(NC(=O)c3cccc(OC)c3OC)ccc2n1 | NaN | 4 | 0 | 0.146 | NaN | 4 | 0 | -4.55 |
| NCGC00104045-02 | COc1ccc(Nc2c3c(nc4ccc(NC(=O)/C=C/c5cccc(OC)c5OC)cc24)CCCC3)cc1OC | 3.981072 | 3 | 78.787 | -2.666 | 10 | -2.1 | -100.256 | -98.601 |
| NCGC00104429-01 | Cc1cc(N2CCN(C)CC2)nc2ccc(NC(=O)c3ccc(C(C)(C)C)cc3)cc12 | 4.466836 | 3 | 107.959 | -3.214 | 14.125375 | -3 | -113.042 | -99.598 |
| NCGC00104306-01 | CCCCN(CC)CCCNC(=O)C1CCN(c2nc3ccc(CC)cc3s2)CC1 | 4.466836 | 3 | 119.918 | -2.995 | 12.589254 | -3 | -106.139 | -99.53 |
| NCGC00104314-01 | CCCCN(C)CCCNC(=O)C1CCN(c2nc3ccc(CC)cc3s2)CC1 | NaN | 4 | 0 | 6.903 | 11.220185 | -3 | -60.806 | -58.164 |
| NCGC00104473-01 | CCN(CC)CC(C)CNC(=O)c1cc(-c2ccc(Br)s2)nc2ccccc12 | 4.466836 | 3 | 83.088 | -3.178 | 12.589254 | -3 | -105.916 | -99.763 |
| NCGC00104469-01 | CC(C)CN(CCNC(=O)c1cc(-c2cccs2)nc2ccccc12)CC(C)C | NaN | 4 | 0 | 0 | 4.466836 | 4 | 16.714 | -1.994 |
| NCGC00104467-01 | CC(C)CN(CCCNC(=O)c1cc(-c2cccs2)nc2ccccc12)CC(C)C | NaN | 4 | 0 | 1.278 | NaN | 4 | 0 | -10.678 |
| NCGC00104475-01 | CC(C)CN(CCNC(=O)c1cc(-c2ccc(Cl)s2)nc2ccccc12)CC(C)C | NaN | 4 | 0 | 1.169 | NaN | 4 | 0 | -3.301 |
| NCGC00105929-01 | CC(C)CN(CCCNC(=O)c1cc(-c2ccc(Br)s2)nc2ccccc12)CC(C)C | NaN | 4 | 0 | -0.11 | NaN | 4 | 0 | -4.163 |
| NCGC00105957-01 | CCc1ccc(CNC(=O)c2cc(-c3ccc(Cl)s3)nc3ccccc23)cc1 | NaN | 4 | 0 | 0.511 | NaN | 4 | 0 | -4.114 |
| NCGC00105965-01 | Cc1ccc(NC(=O)c2cc(-c3ccc(Br)s3)nc3ccccc23)cc1F | NaN | 4 | 0 | -0.913 | 4.466836 | 4 | 16.777 | -2.42 |
| NCGC00105989-01 | COc1cccc(CNC(=O)c2cc(-c3ccc(Cl)s3)nc3ccccc23)c1 | NaN | 4 | 0 | -1.023 | NaN | 4 | 0 | -2.498 |
| NCGC00105959-01 | CCOc1ccc(CNC(=O)c2cc(-c3ccc(Cl)s3)nc3ccccc23)cc1 | NaN | 4 | 0 | 0.548 | NaN | 4 | 0 | -6.351 |
| NCGC00107851-01 | CCC1=Nc2cc(C(=O)NCCN(CC)CC)ccc2Sc2ccccc21 | 14.125375 | 4 | 25.331 | 21.768 | NaN | 4 | 0 | -9.265 |
| NCGC00108136-01 | COc1ccc(Nc2cc(C(=O)NCc3ccccc3)c3ccccc3n2)cc1OC | NaN | 4 | 0 | -0.804 | NaN | 4 | 0 | -1.462 |
| NCGC00108132-01 | COc1ccc(Nc2cc(C(=O)NC3CCC(C)CC3)c3ccccc3n2)cc1OC | NaN | 4 | 0 | 4.419 | NaN | 4 | 0 | 0.745 |
| NCGC00108164-01 | CCN(CCCNC(=O)c1cc(Nc2ccc(OC)cc2)nc2ccccc12)c1cccc(C)c1 | NaN | 4 | 0 | -1.023 | NaN | 4 | 0 | -7.754 |
| NCGC00108168-01 | CCCCN(CCCNC(=O)c1cc(Nc2ccc(OC)cc2)nc2ccccc12)Cc1ccccc1 | NaN | 4 | 0 | -0.037 | NaN | 4 | 0 | 0.978 |
| NCGC00108162-01 | COc1ccc(Nc2cc(C(=O)NCc3ccccc3)c3ccccc3n2)cc1 | NaN | 4 | 0 | -1.023 | NaN | 4 | 0 | -2.401 |
| NCGC00108208-01 | CCc1ccccc1Nc1cc(C(=O)NCCc2ccccc2)c2ccccc2n1 | NaN | 4 | 0 | 0.438 | 5.011872 | 4 | -17.584 | -19.943 |
| NCGC00108204-01 | CCc1ccccc1Nc1cc(C(=O)NC(C)c2ccccc2)c2ccccc2n1 | NaN | 4 | 0 | -0.292 | NaN | 4 | 0 | -12.188 |
| NCGC00108258-01 | Cc1ccc(C)c(Nc2cc(C(=O)NC3CCCC3)c3ccccc3n2)c1 | NaN | 4 | 0 | -0.986 | NaN | 4 | 0 | -3.698 |
| NCGC00108260-01 | Cc1ccc(C)c(Nc2cc(C(=O)NC3CCCCC3)c3ccccc3n2)c1 | NaN | 4 | 0 | -0.584 | NaN | 4 | 0 | 0.426 |
| NCGC00108234-01 | COc1ccc(CNC(=O)c2cc(Nc3ccccc3C)nc3ccccc23)cc1 | NaN | 4 | 0 | 0.146 | NaN | 4 | 0 | 0.542 |
| NCGC00108206-01 | CCc1ccccc1Nc1cc(C(=O)NC2CCC(C)CC2)c2ccccc2n1 | NaN | 4 | 0 | 6.465 | NaN | 4 | 0 | -3.543 |
| NCGC00108295-01 | Cc1cc(C)cc(Nc2cc(C(=O)NC3CCCCC3)c3ccccc3n2)c1 | NaN | 4 | 0 | 0.402 | 4.466836 | 4 | 31.007 | -1.007 |
| NCGC00108299-01 | Cc1cc(C)cc(Nc2cc(C(=O)NCCCN(C)C3CCCCC3)c3ccccc3n2)c1 | 4.466836 | 3 | 108.329 | -3.214 | 14.125375 | -3 | -109.037 | -99.724 |
| NCGC00108323-01 | CC(C)c1ccc(Nc2cc(C(=O)NCc3cccs3)c3ccccc3n2)cc1 | NaN | 4 | 0 | -0.146 | NaN | 4 | 0 | -1.733 |
| NCGC00108944-01 | COc1ccc(OC)c(CNC(=O)c2nn(-c3ccc(OC)c(Cl)c3)c(=O)c3c2c2ccccc2n3C)c1 | NaN | 4 | 0 | -1.023 | NaN | 4 | 0 | -1.81 |
| NCGC00108995-02 | CCCCN(CC)CCNC(=O)c1cc2c(-c3ccc(Cl)cc3)nn(C)c2s1 | 4.466836 | 3 | 78.292 | -3.178 | 10 | -2.1 | -96.437 | -99.724 |
| NCGC00108997-01 | CCCN(CCC)CCNC(=O)c1cc2c(-c3ccc(Cl)cc3)nn(C)c2s1 | 3.981072 | 3 | 79.435 | -3.214 | 11.220185 | -2.1 | -103.917 | -99.647 |
| NCGC00108999-01 | CCCN(CCC)CCCNC(=O)c1cc2c(-c3ccc(Cl)cc3)nn(C)c2s1 | 4.466836 | 3 | 34.241 | -2.995 | 14.125375 | -3 | -114.343 | -99.424 |
| NCGC00109469-01 | CCC(C)NC(=O)CSCc1nc(-c2ccc(C)cc2)oc1C | 14.125375 | 4 | 12.402 | 10.628 | NaN | 4 | 0 | -2.953 |
| NCGC00109473-01 | COc1ccccc1CNC(=O)C1CCN(Cc2nc(-c3ccc(OC)c(OC)c3)oc2C)CC1 | NaN | 4 | 0 | -1.424 | NaN | 4 | 0 | -5.325 |
| NCGC00109483-01 | CCCCNC(=O)C1CCN(Cc2nc(-c3ccc(OC)c(OC)c3)oc2C)CC1 | NaN | 4 | 0 | 0.402 | NaN | 4 | 0 | -1.743 |
| NCGC00109477-01 | COCCCNC(=O)C1CCN(Cc2nc(-c3ccc(OC)c(OC)c3)oc2C)CC1 | 2.818383 | 4 | 13.264 | -1.169 | NaN | 4 | 0 | -3.582 |
| NCGC00109537-01 | Cc1ccc(CCNC(=O)C2CCN(Cc3nc(-c4ccccc4C)oc3C)CC2)cc1 | NaN | 4 | 0 | 8.181 | NaN | 4 | 0 | -11.027 |
| NCGC00109513-02 | Cc1oc(-c2ccc(Cl)cc2)nc1CN1CCC(C(=O)NCCCN2CCN(c3cccc(Cl)c3)CC2)CC1 | NaN | 4 | 0 | -3.104 | 3.981072 | -1.1 | -101.149 | -99.085 |
| NCGC00109541-01 | Cc1ccccc1-c1nc(CN2CCC(C(=O)NCCc3ccccc3)CC2)c(C)o1 | 14.125375 | 4 | 20.947 | 18.371 | NaN | 4 | 0 | -9.265 |
| NCGC00109551-01 | CCCCSCCCNC(=O)C1CCN(Cc2nc(-c3cccc(Br)c3)oc2C)CC1 | 11.220185 | 4 | 20.897 | 18.7 | NaN | 4 | 0 | -2.662 |
| NCGC00109545-01 | CCc1ccc(-c2nc(CN3CCC(C(=O)NCCc4ccc(Cl)cc4)CC3)c(C)o2)cc1 | NaN | 4 | 0 | -0.329 | NaN | 4 | 0 | -5.228 |
| NCGC00109573-01 | Cc1oc(-c2cccc(Cl)c2)nc1CN1CCC(C(=O)NCCc2ccc(Cl)cc2)CC1 | NaN | 4 | 0 | 2.739 | 12.589254 | 4 | -12.971 | -12.024 |
| NCGC00109547-01 | CCc1ccc(-c2nc(CN3CCC(C(=O)NCCc4ccc(C)cc4)CC3)c(C)o2)cc1 | NaN | 4 | 0 | 6.245 | NaN | 4 | 0 | -6.312 |
| NCGC00109569-01 | Cc1ccc(C)c(N2CCN(C(=O)C3CCN(Cc4nc(-c5cccc(Cl)c5)oc4C)CC3)CC2)c1 | 4.466836 | 3 | 61.179 | -3.178 | 12.589254 | -3 | -108.53 | -98.843 |
| NCGC00109579-01 | Cc1oc(-c2cccc(Cl)c2)nc1CN1CCC(C(=O)NCc2ccc(F)cc2)CC1 | NaN | 4 | 0 | 9.24 | 8.912509 | 4 | -13.206 | -13.766 |
| NCGC00109583-01 | Cc1oc(-c2cccc(Cl)c2)nc1CN1CCC(C(=O)NCCCN2CCN(c3ccccc3F)CC2)CC1 | 4.466836 | 4 | 15.884 | -2.885 | 12.589254 | -3 | -106.249 | -98.921 |
| NCGC00109639-01 | CCc1ccc(-c2nc(CN3CCCC(C(=O)NCc4ccccc4Cl)C3)c(C)o2)cc1 | NaN | 4 | 0 | -2.922 | 11.220185 | -3 | -91.118 | -89.065 |
| NCGC00110535-02 | O=C(NCCCN1CCN(c2ccc(F)cc2)CC1)c1cc2c(Cl)nc3ccccc3c2s1 | NaN | 4 | 0 | -1.863 | NaN | 4 | 0 | -4.908 |
| NCGC00109581-01 | Cc1oc(-c2cccc(Cl)c2)nc1CN1CCC(C(=O)NCCCN2CCN(c3cccc(Cl)c3)CC2)CC1 | 4.466836 | 3 | 120.203 | -3.068 | 12.589254 | -3 | -107.551 | -99.317 |
| NCGC00110592-01 | COc1ccc(CNC(=O)c2ccc3c(Cl)c4c(nc3c2)CCCC4)c(OC)c1 | NaN | 4 | 0 | 3.944 | NaN | 4 | 0 | 0.687 |
| NCGC00110903-01 | COc1ccc(OC)c(NC(=O)CSc2c3c(nc4cc(Cl)ccc24)CCCC3)c1 | NaN | 4 | 0 | 0.657 | NaN | 4 | 0 | 2.595 |
| NCGC00110843-01 | CCC1CCCCN1CCCNC(=O)c1cc2c(Cl)nc3ccc(C)cc3c2s1 | 4.466836 | 3 | 101.362 | -3.104 | 12.589254 | -3 | -107.363 | -99.608 |
| NCGC00110567-01 | CC1CCCN(CCCNC(=O)c2cc3c(Cl)nc4ccccc4c3s2)C1 | 4.466836 | 3 | 42.689 | -0.329 | 12.589254 | -3 | -79.548 | -74.728 |
| NCGC00112164-01 | CCC1CCCCN1CCCNC(=O)c1c(O)nc2c(C)cccn2c1=O | NaN | 4 | 0 | -0.767 | NaN | 4 | 0 | -4.753 |
| NCGC00113395-01 | CCOc1ccccc1CNC(=O)c1ccc2nc(N3CCC(C)CC3)sc2c1 | NaN | 4 | 0 | 0.767 | NaN | 4 | 0 | -1.346 |
| NCGC00113265-02 | COc1ccc(Br)cc1CNC(=O)C1CCN(c2nnc(-n3cccc3CNc3ccc(C)cc3)s2)CC1 | NaN | 4 | 0 | 5.442 | NaN | 4 | 0 | -3.698 |
| NCGC00111254-01 | Cc1cccc(CN2CCC(CNc3ncnc4onc(-c5ccc(Cl)cc5)c34)CC2)c1 | 4.466836 | 4 | 13 | 5.442 | 14.125375 | -3 | -98.348 | -87.042 |
| NCGC00113653-01 | O=C(NCc1cccc(Cl)c1)c1ccc(NC(=O)N2CCSc3ccccc32)cc1 | NaN | 4 | 0 | -0.183 | NaN | 4 | 0 | -2.749 |
| NCGC00119149-01 | CC(Oc1ccccc1F)C(=O)Nc1ccc(-c2nc3ccccc3s2)cc1 | NaN | 4 | 0 | 0.475 | NaN | 4 | 0 | -5.663 |
| NCGC00113606-01 | CCN(CC)CCCNC(=O)c1ccc2nc(N3CCCCC3)sc2c1 | NaN | 4 | 0 | 0.548 | 0.316228 | 4 | -26.442 | 0.91 |
| NCGC00121677-01 | O=C(NCCCN1CCCCC1)c1ccc2c(c1)sc1nc(-c3ccc(F)cc3)cn12 | 4.466836 | 3 | 47.037 | -3.068 | 11.220185 | -2.1 | -110.876 | -99.453 |
| NCGC00121679-01 | CCC1CCCCN1CCCNC(=O)c1ccc2c(c1)sc1nc(-c3ccc(F)cc3)cn12 | 4.466836 | 3 | 98.984 | -3.214 | 5.011872 | -1.1 | -95.459 | -99.618 |
| NCGC00121651-01 | CCN(CC)CCNC(=O)c1ccc(-n2c(-c3ccccc3)cc3c2CCCC3)cc1 | 4.466836 | 3 | 93.482 | -3.141 | 12.589254 | -3 | -109.04 | -99.308 |
| NCGC00121699-01 | Cc1ccc(-c2cn3c(n2)sc2cc(C(=O)NCCCN4CCCCC4)ccc23)cc1 | 4.466836 | 3 | 90.255 | -3.068 | 11.220185 | -2.1 | -110.229 | -99.163 |
| NCGC00121711-01 | CCOc1ccc(-c2cn3c(n2)sc2cc(C(=O)NCCCN4CCCCC4)ccc23)cc1 | 4.466836 | 3 | 74.129 | -2.885 | 10 | -2.1 | -100.106 | -99.569 |
| NCGC00121701-01 | Cc1ccc(-c2cn3c(n2)sc2cc(C(=O)NCCCN4CCCCCC4)ccc23)cc1 | 4.466836 | 3 | 91.775 | -2.776 | 11.220185 | -2.1 | -111.208 | -99.095 |
| NCGC00121793-01 | COc1ccc(-c2cn3c(n2)sc2cc(C(=O)NCc4ccccc4OC)ccc23)cc1 | NaN | 4 | 0 | 4.748 | NaN | 4 | 0 | -1.607 |
| NCGC00121705-01 | CCCN1CCN(CCCNC(=O)c2ccc3c(c2)sc2nc(-c4ccc(C)cc4)cn23)CC1 | 4.466836 | 3 | 68.66 | -2.228 | 11.220185 | -2.1 | -112.267 | -98.64 |
| NCGC00121713-01 | CCOc1ccc(-c2cn3c(n2)sc2cc(C(=O)NCCCN4CCCCCC4)ccc23)cc1 | NaN | 4 | 0 | -3.178 | 12.589254 | -3 | -107.625 | -99.385 |
| NCGC00121987-01 | COc1ccccc1CNC(=O)c1ccc2c(c1)sc1nc(-c3ccc(F)cc3)cn12 | NaN | 4 | 0 | 0.292 | NaN | 4 | 0 | -1.655 |
| NCGC00121823-01 | COc1ccc(-c2cn3c(n2)sc2cc(C(=O)NCCCN4CCCCC4)ccc23)cc1 | 4.466836 | 3 | 68.349 | -2.995 | 11.220185 | -2.1 | -109.91 | -98.475 |
| NCGC00126039-01 | CC(=O)c1ccc(NC(=O)c2cccc(NC(=O)N3CCSc4ncccc43)c2)cc1 | NaN | 4 | 0 | 1.242 | NaN | 4 | 0 | -0.581 |
| NCGC00124677-01 | O=C(c1nc(-c2ccc(Cl)cc2)n2c1CCCCC2)N1CCN(c2ccccc2)CC1 | NaN | 4 | 0 | 1.753 | NaN | 4 | 0 | -3.437 |
| NCGC00122847-01 | COc1ccccc1CNC(=O)CSc1ccc(-c2sc(-c3ccccc3)nc2C)nn1 | 14.125375 | 4 | 14.791 | 12.454 | 14.125375 | 4 | -13.857 | -14.047 |
| NCGC00123977-01 | CCN(CC)CCCNC(=O)Cn1c(-c2cccs2)cc2cc(Cl)ccc21 | 11.220185 | 3 | 83.633 | 77.831 | 14.125375 | -3 | -34.161 | -33.051 |
| NCGC00128787-01 | COc1ccccc1CNc1nccn(-c2ccc(F)cc2)c1=O | NaN | 4 | 0 | 3.908 | NaN | 4 | 0 | -3.834 |
| NCGC00126147-01 | CCOc1ccccc1CNC(=O)C1CCCN(c2nnc(C)c3c(C)n(-c4ccccc4)nc23)C1 | NaN | 4 | 0 | -0.073 | NaN | 4 | 0 | -4.976 |
| NCGC00126165-01 | CCOc1ccc(CNC(=O)Cc2ccc(NC(=O)N3CCSc4ncccc43)cc2)cc1 | NaN | 4 | 0 | 0.621 | NaN | 4 | 0 | 1.094 |
| NCGC00128825-01 | COc1ccc(-n2ccnc(NCc3ccccc3OC)c2=O)cc1 | NaN | 4 | 0 | -0.438 | NaN | 4 | 0 | 3.311 |
| NCGC00126043-01 | Cc1cc(Cl)ccc1NC(=O)c1cccc(NC(=O)N2CCSc3ncccc32)c1 | NaN | 4 | 0 | 2.447 | NaN | 4 | 0 | -6.254 |
| NCGC00126163-01 | COc1cccc(CNC(=O)Cc2ccc(NC(=O)N3CCSc4ncccc43)cc2)c1OC | NaN | 4 | 0 | -0.365 | NaN | 4 | 0 | -0.252 |
| NCGC00129367-01 | CCCNC(=O)c1c2c(nc3ccccc13)C(=O)N(C1CCCCC1)C2 | NaN | 4 | 0 | 0.146 | NaN | 4 | 0 | -3.601 |
| NCGC00132442-01 | CCN(CC)CCCNC(=O)Cc1csc2nc(-c3ccc(Cl)cc3)cn12 | 11.220185 | 4 | 26.384 | 24.47 | NaN | 4 | 0 | -4.773 |
| NCGC00133606-01 | CCCNC(=O)c1ccc2c(c1)nc1n2CCCCC1 | NaN | 4 | 0 | 2.009 | NaN | 4 | 0 | -3.998 |
| NCGC00132824-01 | COc1ccccc1CNC(=O)c1ccc(-c2nc(CN3CCc4ccccc43)c(C)o2)cc1 | NaN | 4 | 0 | 6.355 | NaN | 4 | 0 | -3.263 |
| NCGC00130952-01 | COc1ccccc1-c1nc(CNC(=O)NC2CCCCC2)c(C)o1 | NaN | 4 | 0 | -1.315 | NaN | 4 | 0 | 1.249 |
| NCGC00129233-01 | CCN(CC)CCNC(=O)c1c2c(nc3ccccc13)C(=O)N(C1CCCCC1)C2 | NaN | 4 | 0 | -1.059 | NaN | 4 | 0 | -4.318 |
| NCGC00133707-01 | Cc1cccn2c(=O)cc(COc3ccc(NC(=O)c4cccc(F)c4)cc3)nc12 | NaN | 4 | 0 | 7.706 | NaN | 4 | 0 | -3.166 |
| NCGC00134320-01 | COc1cccc(OC)c1C(=O)Nc1cccc(-c2cn3ccsc3n2)c1 | NaN | 4 | 0 | -1.753 | NaN | 4 | 0 | -5.45 |
| NCGC00134322-01 | COc1cccc(C(=O)Nc2cccc(-c3cn4ccsc4n3)c2)c1OC | NaN | 4 | 0 | -1.607 | NaN | 4 | 0 | -4.618 |
| NCGC00134412-01 | COc1ccc(OC)c(CNC(=O)c2ccc(NC(=O)c3nsc4ccccc34)cc2)c1 | NaN | 4 | 0 | 4.456 | NaN | 4 | 0 | -3.563 |
| NCGC00135406-01 | CCOc1ccc(C(=O)Nc2cc(-c3cn4cccnc4n3)ccc2C)cc1OCC | NaN | 4 | 0 | -1.023 | NaN | 4 | 0 | 0.029 |
| NCGC00133999-01 | COc1cccc(C(=O)Nc2cccc(-c3cn4c(C)csc4n3)c2)c1OC | NaN | 4 | 0 | -0.329 | 12.589254 | 4 | -12.031 | -14.231 |
| NCGC00137596-01 | CCCNC(=O)C1CCN(Cc2cc3ccccc3n2Cc2ccccc2)CC1 | NaN | 4 | 0 | 7.012 | NaN | 4 | 0 | -10.349 |
| NCGC00139452-01 | Cc1oc(-c2cccc(Br)c2)nc1CN1CCC(C(=O)NCCc2ccc(Cl)cc2)CC1 | NaN | 4 | 0 | -0.183 | NaN | 4 | 0 | -5.237 |
| NCGC00137594-01 | CN(C)CCNC(=O)C1CCN(Cc2cc3ccccc3n2Cc2ccccc2)CC1 | 4.466836 | 3 | 49.876 | 9.752 | 14.125375 | -3 | -50.626 | -45.791 |
| NCGC00137698-01 | COCCNC(=O)C1CCN(Cc2cc3ccccc3n2Cc2ccc(C)cc2)CC1 | 14.125375 | 4 | 15.484 | 13.367 | 1 | 4 | 6.236 | 1.375 |
| NCGC00137690-01 | COCCCNC(=O)C1CCN(Cc2cc3ccccc3n2Cc2cccc(C)c2)CC1 | NaN | 4 | 0 | 3.981 | NaN | 4 | 0 | -5.615 |
| NCGC00137658-01 | COCCCNC(=O)C1CCN(Cc2cc3ccccc3n2Cc2ccccc2C)CC1 | NaN | 4 | 0 | 5.625 | NaN | 4 | 0 | -4.657 |
| NCGC00137638-01 | Cc1ccccc1Cn1c(CN2CCC(C(=O)NCCN(C)C)CC2)cc2ccccc21 | 3.981072 | 3 | 48.594 | -3.141 | 12.589254 | -3 | -108.577 | -99.085 |
| NCGC00139640-01 | CCCCN(C)CCCNC(=O)c1cc(-c2cccnc2)nc2ccccc12 | NaN | 4 | 0 | -0.95 | NaN | 4 | 0 | -4.114 |
| NCGC00140130-01 | O=C(NCCc1ccccc1)c1cc(Sc2cccc(Cl)c2)nc2ccccc12 | NaN | 4 | 0 | 0.621 | NaN | 4 | 0 | -0.281 |
| NCGC00139666-01 | O=C(NCc1ccccc1)c1cc(Sc2cccc(Cl)c2)nc2ccccc12 | NaN | 4 | 0 | -1.169 | NaN | 4 | 0 | -0.378 |
| NCGC00139766-01 | CCC(C)NC(=O)c1cc(Sc2cccc(Cl)c2)nc2ccccc12 | NaN | 4 | 0 | -1.461 | NaN | 4 | 0 | -4.134 |
| NCGC00140220-01 | CCCCNC(=O)c1cc(Sc2ccc(OCC)cc2)nc2ccccc12 | NaN | 4 | 0 | -0.329 | NaN | 4 | 0 | -4.018 |
| NCGC00140216-01 | CCOc1ccc(Sc2cc(C(=O)NCc3ccc(C)cc3)c3ccccc3n2)cc1 | NaN | 4 | 0 | -1.205 | NaN | 4 | 0 | -7.164 |
| NCGC00139820-01 | CCc1c(-c2ccccc2)nc2ccccc2c1C(=O)NCCc1ccccc1 | NaN | 4 | 0 | -1.899 | NaN | 4 | 0 | -6.738 |
| NCGC00140188-01 | CCc1ccc(Sc2cc(C(=O)NC(C)CC)c3ccccc3n2)cc1 | NaN | 4 | 0 | -1.169 | NaN | 4 | 0 | 3.127 |
| NCGC00140331-02 | COc1ccc(N2CCN(CCCNC(=O)c3cc(Nc4ccccc4OC)nc4ccccc34)CC2)cc1 | 11.220185 | 3 | 61.816 | 59.24 | 11.220185 | -3 | -37.198 | -39.876 |
| NCGC00140332-01 | COc1ccc(Nc2cc(C(=O)NCCCN(C)Cc3ccccc3)c3ccccc3n2)cc1OC | 10 | 3 | 40.43 | 37.655 | 12.589254 | -3 | -59.096 | -56.789 |
| NCGC00140218-01 | CCOc1ccc(Sc2cc(C(=O)NCCc3ccccc3)c3ccccc3n2)cc1 | NaN | 4 | 0 | -0.292 | 1.995262 | 4 | -13.157 | -5.547 |
| NCGC00140493-02 | Cc1ccccc1Nc1cc(C(=O)NCCCN2CCN(c3cccc(Cl)c3)CC2)c2ccccc2n1 | 4.466836 | 3 | 64.004 | 13.148 | 5.011872 | 4 | -11.022 | -15.238 |
| NCGC00141370-01 | COc1ccccc1CNc1nccn(-c2ccccc2)c1=O | NaN | 4 | 0 | -0.767 | NaN | 4 | 0 | -2.169 |
| NCGC00140230-01 | CCOc1ccc(Sc2cc(C(=O)NCc3ccc(F)cc3)c3ccccc3n2)cc1 | NaN | 4 | 0 | 0.584 | NaN | 4 | 0 | -1.791 |
| NCGC00140438-01 | Cc1cccc(-c2nc(CSCC(=O)N3CCN(c4cccc(C)c4C)CC3)c(C)o2)c1 | 14.125375 | 4 | 16.766 | 15.522 | NaN | 4 | 0 | -3.524 |
| NCGC00140342-01 | CC(C)c1ccc(Nc2cc(C(=O)NCCCN(C)Cc3ccccc3)c3ccccc3n2)cc1 | NaN | 4 | 0 | 0.11 | NaN | 4 | 0 | 0.194 |
| NCGC00141939-01 | O=C(NC(=O)c1c(F)cccc1F)Nc1ccc(-c2cc(C(F)(F)F)nn2-c2ccccc2)cc1 | NaN | 4 | 0 | -1.023 | NaN | 4 | 0 | -2.943 |
| NCGC00393906-01 | COC[C@H]1C[C@@H](NC(=O)Nc2cccc(-c3noc(C)n3)c2)[C@H](O)[C@@H]1O | NaN | 4 | 0 | 0.037 | NaN | 4 | 0 | 0.871 |
| NCGC00396549-01 | CN1C(=O)CO[C@H](c2ccc(NC(=O)c3cccc(F)c3)cc2)[C@H]1CO | NaN | 4 | 0 | -1.096 | NaN | 4 | 0 | -1.52 |
| NCGC00394129-01 | COc1ccccc1CNC(=O)[C@@H]1Cc2c[nH]c3cccc(c23)[C@H](CC(C)C)N1 | NaN | 4 | 0 | -0.183 | NaN | 4 | 0 | -1.191 |
| NCGC00411161-01 | Fc1ccc(Nc2cc3c(cn2)CN(Cc2ccccc2)CCN3)cc1 | 19.952623 | 3 | 72.804 | 53.798 | 11.220185 | -2.2 | -34.185 | -34.29 |
| NCGC00411920-01 | COc1ccc(Nc2n[nH]c(C(O)c3ccccc3)n2)cc1OC | NaN | 4 | 0 | 1.132 | NaN | 4 | 0 | 0.852 |
| NCGC00391713-01 | CN(C)c1ccc(C(=O)N[C@H]2C[C@@H](c3nc(-c4ccc5ccccc5c4)no3)N(C)C2)cc1 | NaN | 4 | 0 | 7.487 | NaN | 4 | 0 | -8.413 |
| NCGC00411730-01 | COc1ccccc1-c1cc2c(ccn2C(C)C)c(C(=O)NCc2c(C)cc(C)[nH]c2=O)n1 | NaN | 4 | 0 | -0.219 | 10 | 4 | 9.009 | 5.412 |
| NCGC00416332-01 | CC1(CNCc2ccccc2)COc2ccc(C(=O)Nc3cccnc3)cc2N1 | NaN | 4 | 0 | 0.584 | NaN | 4 | 0 | -5.886 |
| NCGC00411607-01 | COc1ccccc1-c1cc2c(ccn2C)c(C(=O)NCc2c(C)cc(C)[nH]c2=O)n1 | NaN | 4 | 0 | -0.511 | 7.079458 | 4 | -9.754 | -14.483 |
| NCGC00400548-01 | CC(C)(C)c1ccc(-c2cc(C[C@H]3CNCC[C@H]3CC(=O)N3CCCCC3)no2)cc1 | 7.943282 | 3 | 66.054 | -2.849 | 22.387211 | -3 | -133.331 | -99.114 |
| NCGC00421798-01 | CCCN(C)C(=O)c1c(-c2ccccc2)noc1[C@H](C)O | NaN | 4 | 0 | 5.004 | NaN | 4 | 0 | -1.191 |
| NCGC00423521-01 | O=C(N[C@H]1C[C@@H]2OCC[C@H]12)c1cccnc1SCCc1ccccc1 | NaN | 4 | 0 | -1.68 | NaN | 4 | 0 | -3.359 |
| NCGC00423648-01 | CCCOc1ccccc1CCCNC(=O)[C@H]1CCC[C@@H](O)C1 | NaN | 4 | 0 | -1.132 | NaN | 4 | 0 | 1.2 |
| NCGC00423695-01 | COc1cccc2c1ccn2CCNC(=O)C1CC(=O)N(c2n[nH]c3ccccc23)C1 | NaN | 4 | 0 | -0.767 | NaN | 4 | 0 | -4.143 |
| NCGC00419541-01 | CC(C)(Oc1nn(Cc2cccc(Cl)c2)c2ccccc12)C(=O)Nc1cccnc1 | 15.848932 | 4 | 14.305 | 14.207 | 15.848932 | 4 | -29.692 | -33.157 |
| NCGC00425617-01 | CCCCNC(=O)c1cc(NC(=O)CC)cc2nc(C)ccc12 | NaN | 4 | 0 | -1.315 | NaN | 4 | 0 | 0.707 |
| NCGC00420204-01 | Cc1cc(F)ccc1NC(=O)c1cnc2c(c1)N([S+](=O)([O-])c1cccnc1)CCC2 | NaN | 4 | 0 | -1.388 | NaN | 4 | 0 | -4.085 |
| NCGC00423091-01 | CCO[C@H]1CN(C(=O)Cc2ccccc2OCc2ccccc2)C[C@@H]1O | NaN | 4 | 0 | 1.68 | NaN | 4 | 0 | 3.33 |
| NCGC00424151-01 | COCCn1cc(C(=O)NC2CCCc3c2[nH]c2ccccc32)c2ccccc2c1=O | NaN | 4 | 0 | -0.329 | NaN | 4 | 0 | -7.28 |
| NCGC00426532-01 | O=C(Nc1ccccc1)c1cnc(-c2ccc(F)cc2)nc1-c1ccncc1 | NaN | 4 | 0 | -0.329 | NaN | 4 | 0 | -2.449 |
| NCGC00425621-01 | CCC(=O)Nc1cc(C(=O)NCc2ccc(N3CCOCC3)cc2)c2ccc(C)nc2c1 | NaN | 4 | 0 | 1.534 | NaN | 4 | 0 | -1.878 |
| NCGC00425619-01 | CCC(=O)Nc1cc(C(=O)NCc2ccc(N3CCN(C)CC3)cc2)c2ccc(C)nc2c1 | NaN | 4 | 0 | 1.79 | NaN | 4 | 0 | -2.585 |
| NCGC00424661-01 | CCCC1=Nc2cc(C(=O)NCCCN3CCCCC3CC)ccc2Sc2ccc(C)cc21 | 4.466836 | 3 | 112.095 | -3.141 | 12.589254 | -3 | -108.211 | -99.53 |
| NCGC00427016-01 | CC(C)CCNC(=O)c1ccc(-c2ccc(N3CCSCC3)nc2)cc1 | NaN | 4 | 0 | -0.584 | NaN | 4 | 0 | 3.592 |
| NCGC00426068-01 | O=C(CCC1CCN(C(=O)c2ccccc2)CC1)NCCCN1CCCc2ccccc21 | NaN | 4 | 0 | 0.256 | NaN | 4 | 0 | -5.857 |
| NCGC00425603-01 | Cc1ccc2c(C(=O)NCc3ccccc3C(F)(F)F)cc(NC(=O)C(C)C)cc2n1 | NaN | 4 | 0 | -0.804 | NaN | 4 | 0 | -1.075 |
| NCGC00427142-01 | CCn1cc(CN2CCCC(COc3ccccc3C(=O)N3CCCC3)C2)c2ccccc21 | 14.125375 | 4 | 14.428 | 11.651 | NaN | 4 | 0 | -7.919 |
| NCGC00425364-01 | O=C(CCC1CCN(Cc2cc3ccccc3o2)CC1)NC1CCN(Cc2ccccc2)C1 | NaN | 4 | 0 | 6.099 | NaN | 4 | 0 | -6.147 |
| NCGC00427266-01 | CCN1CCCC1CNC(=O)CCC1CCN(Cc2cn(CC)c3ccccc23)CC1 | NaN | 4 | 0 | -0.73 | NaN | 4 | 0 | -0.416 |
| NCGC00432871-01 | O=C(Nc1cnc(-c2ccccc2)nc1)c1ccc(F)c(F)c1 | NaN | 4 | 0 | 1.607 | NaN | 4 | 0 | -1.142 |
| NCGC00433004-01 | CC(C)(CN1CCOCC1)NC(=O)c1csc2nc(-c3ccc(F)cc3)cn12 | NaN | 4 | 0 | 1.169 | NaN | 4 | 0 | -7.212 |
| NCGC00435072-01 | COc1ccc(OC)c(CNC(=O)COc2ccc(NC(C)=O)cn2)c1 | NaN | 4 | 0 | 0.475 | NaN | 4 | 0 | 1.317 |
| NCGC00426966-01 | O=C(NCCN1CCCCC1)c1ccc(-c2ccc(N3CCCCC3)nc2)cc1 | 12.589254 | 2.4 | 31.078 | 24.507 | 11.220185 | 4 | -14.383 | -11.946 |
| NCGC00432291-01 | Cc1cc(=O)n(CC(=O)Nc2ccc(Cl)cc2F)c(N(C)Cc2ccccc2)n1 | NaN | 4 | 0 | -0.767 | 5.011872 | 4 | -23.512 | -10.426 |
| NCGC00436982-01 | COc1cccc(C(=O)NC2CCN(c3cc(-c4ccccc4)n[nH]3)CC2)c1OC | NaN | 4 | 0 | -1.278 | NaN | 4 | 0 | -1.965 |
| NCGC00428202-01 | CSc1ncccc1C(=O)NC1CCCN(CCCc2ccccc2)C1 | NaN | 4 | 0 | -1.899 | NaN | 4 | 0 | -2.691 |
| NCGC00426964-01 | O=C(NCCCN1CCCC1)c1ccc(-c2ccc(N3CCCCC3)nc2)cc1 | NaN | 4 | 0 | 3.47 | 14.125375 | 4 | 27.42 | 19.914 |
| NCGC00428525-01 | COc1cccc(NC(=O)CCC[C@H]2NC[C@@H]3C[C@H]2CN(Cc2ccccc2)C3)c1 | NaN | 4 | 0 | -0.365 | NaN | 4 | 0 | -0.91 |
| NCGC00432913-01 | O=C(Nc1cnc(-c2ccc(Cl)cc2)nc1)c1cccc(F)c1 | NaN | 4 | 0 | 2.228 | NaN | 4 | 0 | -2.062 |
| NCGC00437395-01 | CCc1ccc2nc(C)cc(C(=O)N3CCN(c4ccccc4)C(=O)C3)c2c1 | NaN | 4 | 0 | -0.073 | NaN | 4 | 0 | 7.183 |
| NCGC00437190-01 | COc1cccc(C(=O)NC2CCN(c3cc(-c4cccs4)n[nH]3)CC2)c1OC | NaN | 4 | 0 | -0.95 | NaN | 4 | 0 | -7.125 |
| NCGC00437936-01 | O=C(NC1CCN(c2cc(-c3ccc(Cl)cc3)n[nH]2)C1)c1ccc(F)c(F)c1 | NaN | 4 | 0 | 2.703 | NaN | 4 | 0 | -4.54 |
| NCGC00437970-01 | O=C(NC1CCN(c2cc(-c3ccc(F)cc3)n[nH]2)C1)c1cccc(F)c1 | NaN | 4 | 0 | 0 | 1.122018 | 4 | 8.566 | -0.242 |
| NCGC00437486-01 | CCOc1ccc(C(=O)NC2CCCN(c3cc(-c4ccc(F)cc4)n[nH]3)C2)cc1OCC | NaN | 4 | 0 | 0.95 | NaN | 4 | 0 | -3.185 |
| NCGC00437951-01 | COc1ccc(OC)c2nc(-c3cccc(Cl)c3)c(CN3CCC(O)CC3)cc12 | NaN | 4 | 0 | -1.936 | NaN | 4 | 0 | -0.668 |
| NCGC00437148-01 | O=C(NC1CCN(c2cc(-c3cccs3)n[nH]2)CC1)c1cccc(F)c1 | NaN | 4 | 0 | -1.059 | NaN | 4 | 0 | -3.388 |
| NCGC00446181-01 | O=C(c1cc(Cl)ccc1F)N1CCC[C@H]1c1nc(-c2ccccc2)no1 | NaN | 4 | 0 | -0.877 | 10 | 4 | 12.997 | 6.709 |
| NCGC00447205-01 | O=C(NCc1ccccc1F)c1noc(-c2ccc3c(c2)OCO3)n1 | NaN | 4 | 0 | -1.242 | NaN | 4 | 0 | -13.011 |
| NCGC00437066-01 | COc1cccc(C(=O)NC2CCN(c3cc(-c4ccc(Cl)cc4)n[nH]3)CC2)c1OC | NaN | 4 | 0 | -1.497 | 4.466836 | 4 | 20.323 | -7.135 |
| NCGC00437794-01 | CN(C)c1ccc(C(=O)NCC2CCN(c3cc(-c4ccc(Cl)cc4)n[nH]3)CC2)cc1 | NaN | 4 | 0 | 2.812 | NaN | 4 | 0 | 5.354 |
| NCGC00437975-01 | COc1cccc(CN2CCCC(NC(=O)Nc3ccccc3OC)C2)c1 | NaN | 4 | 0 | 0 | NaN | 4 | 0 | -6.312 |
| NCGC00448221-01 | O=C(Nc1ccc(-c2nnc3n2CCCC3)cc1)c1ccc(F)c(F)c1 | NaN | 4 | 0 | -1.351 | NaN | 4 | 0 | -1.307 |
| NCGC00447560-01 | O=C(NC[C@H]1CCc2ccccc2O1)C1CCN(C(=O)C2CC2)CC1 | NaN | 4 | 0 | -1.644 | 1.258925 | 4 | 23.606 | -1.423 |
| NCGC00447755-01 | CCOc1ccc(-c2nc(C3CC(=O)N(c4ccc(F)c(Cl)c4)C3)no2)cc1OCC | NaN | 4 | 0 | -0.548 | NaN | 4 | 0 | -5.596 |
| NCGC00451887-01 | O=C(NCCc1noc(-c2ccccc2)n1)c1ccc(Cl)s1 | NaN | 4 | 0 | -0.365 | NaN | 4 | 0 | 5.499 |
| NCGC00438172-01 | COc1ccc(-c2cc(N3CCC(NC(=O)c4cccc(F)c4)C3)[nH]n2)cc1 | NaN | 4 | 0 | 0.584 | NaN | 4 | 0 | 2.411 |
| NCGC00440656-01 | Cc1cccc(-c2nc3cc(C)ccc3cc2CN2CCN(CCCO)CC2)c1 | NaN | 4 | 0 | 3.944 | NaN | 4 | 0 | -1.307 |
| NCGC00451900-01 | COc1ccc(C(=O)NCCc2noc(-c3ccncc3)n2)cc1 | NaN | 4 | 0 | -0.329 | NaN | 4 | 0 | 0.232 |
| NCGC00437620-01 | CCOc1ccc(C(=O)NC2CCCN(c3cc(-c4ccc(OC)cc4)n[nH]3)C2)cc1OCC | NaN | 4 | 0 | 1.169 | NaN | 4 | 0 | -2.43 |
| NCGC00443101-01 | CN(C)CC1(O)CCCN(Cc2cccc3ncccc23)C1 | NaN | 4 | 0 | 1.169 | NaN | 4 | 0 | -4.734 |
| NCGC00447763-01 | COc1ccccc1-c1nc(C2CC(=O)N(c3ccc(F)c(Cl)c3)C2)no1 | NaN | 4 | 0 | -1.351 | NaN | 4 | 0 | -3.592 |
| NCGC00457390-01 | CCCc1noc(-c2cccn3c(CNC(=O)C4CC4)nnc23)n1 | NaN | 4 | 0 | 0.694 | NaN | 4 | 0 | 1.036 |
| NCGC00454588-01 | CCOc1ccccc1CNC(=O)c1ccc(CN2C(=O)NC3(CCOCC3)C2=O)cc1 | NaN | 4 | 0 | -0.84 | NaN | 4 | 0 | -1.83 |
| NCGC00457391-01 | CCCc1noc(-c2cccn3c(CNC(=O)C4CCC4)nnc23)n1 | NaN | 4 | 0 | -1.278 | NaN | 4 | 0 | -1.597 |
| NCGC00452463-01 | CCN1c2cc(C(=O)NCc3cc(F)ccc3F)ccc2C(=O)N2CCCCC21 | NaN | 4 | 0 | 3.543 | NaN | 4 | 0 | -3.65 |
| NCGC00453731-01 | O=C(NCC1CC1)c1cccc(OC2CN(C3CCN(Cc4ccccc4)CC3)C2)c1 | NaN | 4 | 0 | 6.063 | NaN | 4 | 0 | -2.73 |
| NCGC00447759-01 | COc1ccc(-c2nc(C3CC(=O)N(c4ccc(F)c(Cl)c4)C3)no2)cc1OC | NaN | 4 | 0 | -1.278 | NaN | 4 | 0 | -1.007 |
| NCGC00457416-01 | CCCc1noc(-c2cccn3c(CNC(=O)C4CCCCC4)nnc23)n1 | NaN | 4 | 0 | 0.329 | NaN | 4 | 0 | -0.397 |
| NCGC00457474-01 | CC(C)c1noc(-c2cccn3c(CNC(=O)C4CCC4)nnc23)n1 | NaN | 4 | 0 | -1.242 | NaN | 4 | 0 | -0.039 |
| NCGC00453300-01 | Cc1ccc(C)c(NC(=O)N2CCCC(c3noc(COc4ccccc4)n3)C2)c1 | NaN | 4 | 0 | -1.169 | NaN | 4 | 0 | -5.083 |
| NCGC00457476-01 | CC(C)c1noc(-c2cccn3c(CNC(=O)C4CCCC4)nnc23)n1 | NaN | 4 | 0 | -1.169 | NaN | 4 | 0 | -4.85 |
| NCGC00457570-01 | CCc1noc(-c2cccn3c(CNC(=O)C4CCCCC4)nnc23)n1 | NaN | 4 | 0 | 0.256 | NaN | 4 | 0 | -1.501 |
| NCGC00457389-01 | CCCc1noc(-c2cccn3c(CNC(=O)C4CCCC4)nnc23)n1 | NaN | 4 | 0 | 2.447 | NaN | 4 | 0 | -3.572 |
| NCGC00458090-01 | O=C(NCc1ccccc1)c1ccccc1OC1CN(Cc2ccc(O)c(F)c2)C1 | NaN | 4 | 0 | -1.972 | NaN | 4 | 0 | -5.199 |
| NCGC00470450-01 | CCc1nc2c(NC(=O)NCCc3ccccc3)cccn2n1 | NaN | 4 | 0 | -0.548 | NaN | 4 | 0 | -0.765 |
| NCGC00469966-01 | CCn1ncc(N2CCC(NC(=O)c3ccc(C)c(F)c3)CC2)cc1=O | NaN | 4 | 0 | 2.447 | NaN | 4 | 0 | -5.77 |
| NCGC00457427-01 | CCCc1noc(-c2cccn3c(CNC(=O)c4cccc(Cl)c4)nnc23)n1 | NaN | 4 | 0 | -2.264 | NaN | 4 | 0 | -6.525 |
| NCGC00471045-01 | Cc1ccc(Cn2c(-c3cccc(Cl)c3)nc(C)c(CCO)c2=O)cc1 | NaN | 4 | 0 | -1.132 | NaN | 4 | 0 | -3.514 |
| NCGC00470997-01 | Cc1nc(-c2ccc(F)cc2)n(Cc2nc(-c3ccccc3)oc2C)c(=O)c1CCO | NaN | 4 | 0 | 1.57 | 14.125375 | 3 | 43.125 | 35.491 |
| NCGC00457380-01 | CC(C)c1noc(-c2cccn3c(CNC(=O)C4CCCCC4)nnc23)n1 | NaN | 4 | 0 | -0.292 | NaN | 4 | 0 | -0.91 |
| NCGC00471428-01 | COc1cc(Cl)ccc1C(=O)N1[C@H]2CC[C@@H]1CC(NC(=O)c1ccncc1)C2 | NaN | 4 | 0 | -0.219 | NaN | 4 | 0 | 0.465 |
| NCGC00457758-01 | Cc1ccc(C(=O)NCc2nnc3cc(-c4nc(C5CC5)no4)ccn23)cc1F | NaN | 4 | 0 | -1.388 | NaN | 4 | 0 | -0.01 |
| NCGC00474627-01 | Cc1cccc(CNC(=O)c2c(C)noc2-c2cccs2)c1 | NaN | 4 | 0 | 0.511 | 0.707946 | 4 | 8.416 | 1.936 |
| NCGC00473777-01 | Cc1noc(-c2ccc3c(c2)OCO3)c1C(=O)NCCCc1ccccc1 | NaN | 4 | 0 | -1.132 | NaN | 4 | 0 | -3.669 |
| NCGC00457469-01 | CC(C)c1noc(-c2cccn3c(CNC(=O)C4CC4)nnc23)n1 | NaN | 4 | 0 | -0.329 | NaN | 4 | 0 | -0.774 |
| NCGC00474629-01 | Cc1cccc(CNC(=O)c2c(C)noc2-c2ccc3c(c2)OCO3)c1 | NaN | 4 | 0 | 0.11 | NaN | 4 | 0 | 6.738 |
| NCGC00469917-01 | Cc1ccc(C(=O)NC2CCN(c3cnn(C)c(=O)c3)CC2)cc1F | NaN | 4 | 0 | -0.438 | NaN | 4 | 0 | -3.766 |
| NCGC00474746-01 | Cc1cccc(CNC(=O)c2c(C)noc2-c2ccccc2Cl)c1 | NaN | 4 | 0 | -1.863 | NaN | 4 | 0 | -2.459 |
| NCGC00477132-01 | Cc1ccc2c(C(=O)Nc3ccc(F)cc3)cnn2c1 | NaN | 4 | 0 | -0.292 | NaN | 4 | 0 | -2.527 |
| NCGC00470957-01 | Cc1ccc(Cn2c(-c3ccc(F)cc3)nc(C)c(CCO)c2=O)cc1 | NaN | 4 | 0 | -0.292 | NaN | 4 | 0 | -3.098 |
| NCGC00476356-01 | COCCNC(=O)c1nc(-c2cccc(C(F)(F)F)c2)c2ccccn12 | NaN | 4 | 0 | 0.804 | 12.589254 | 4 | -15.15 | -11.375 |
| NCGC00477171-01 | Cc1nnn2c(C(=O)NC3CC3)cccc12 | NaN | 4 | 0 | -1.023 | NaN | 4 | 0 | -3.263 |
| NCGC00474538-01 | COc1ccc([C@H](C)NC(=O)c2c(C)noc2-c2cccs2)cc1 | NaN | 4 | 0 | 3.506 | NaN | 4 | 0 | -1.152 |
| NCGC00483777-01 | COc1ccccc1-c1nnc2ccc(C(=O)NCc3ccc(F)cc3)cn12 | NaN | 4 | 0 | -1.096 | NaN | 4 | 0 | -3.137 |
| NCGC00486164-01 | COc1cccc(C(=O)Nc2cc(-c3ccncc3)nn2C)c1 | NaN | 4 | 0 | -1.461 | NaN | 4 | 0 | -2.14 |
| NCGC00484362-01 | CCOCCNC(=O)c1nc2cccnc2n(C2CC2)c1=O | NaN | 4 | 0 | -0.219 | NaN | 4 | 0 | -2.856 |
| NCGC00486418-01 | O=C(NCc1ccccc1F)c1ccc2oc(-c3cccnc3)nc2c1 | NaN | 4 | 0 | 1.644 | NaN | 4 | 0 | -3.456 |
| NCGC00474637-01 | COc1cc(OC)c(CNC(=O)c2c(C)noc2-c2cccs2)c(OC)c1 | NaN | 4 | 0 | 0.292 | 14.125375 | 4 | 14.921 | 9.729 |
| NCGC00486451-01 | COCCn1c(=O)c(C(=O)NC2CC2)nc2cccnc21 | NaN | 4 | 0 | 0.767 | NaN | 4 | 0 | 0.891 |
| NCGC00486123-01 | CC(Nc1nc(N2CCNCCNCC2)nc2ccccc12)c1ccccc1 | NaN | 4 | 0 | 1.644 | NaN | 4 | 0 | 5.412 |
| NCGC00487589-01 | COc1cccc(C(=O)NC2CCN(c3ccnc4ccnn34)CC2)c1 | NaN | 4 | 0 | 5.077 | NaN | 4 | 0 | -2.778 |
| NCGC00486840-01 | CC(Nc1nc(N2CCNCC2)nc2ccccc12)c1ccc(-c2ccccc2)cc1 | 3.548134 | 3 | 79.796 | 3.178 | 12.589254 | -3 | -107.012 | -99.782 |
| NCGC00480767-01 | C[C@@H](Nc1nc(N2CCNCC2)nc2ccccc12)c1ccc2ccccc2c1 | 3.981072 | 2.1 | 109.306 | -3.104 | 14.125375 | -3 | -115.722 | -99.317 |
| NCGC00487758-01 | Cc1ccc(-c2c(C(=O)NCc3ccccc3)nc3sccn23)cc1 | NaN | 4 | 0 | -1.79 | 3.981072 | 4 | -10.012 | -2.701 |
| NCGC00487573-01 | COc1ccc(Nc2ccnc(N3CCN(C(=O)c4ccc(C)cc4)CC3)n2)cc1 | NaN | 4 | 0 | -1.644 | 8.912509 | 4 | -20.007 | -20.998 |
| NCGC00488166-01 | COc1ccccc1-c1cc(C(=O)Nc2ccc(F)cc2)nn2cnnc12 | NaN | 4 | 0 | -0.511 | NaN | 4 | 0 | 5.354 |
| NCGC00489513-01 | COc1ccc(CCNC(=O)c2nc3sccn3c2-c2cccc(C(F)(F)F)c2)cc1 | NaN | 4 | 0 | -1.169 | NaN | 4 | 0 | -7.106 |
| NCGC00489400-01 | COc1cccc(C(=O)NC2CCN(c3nccc(Nc4ccc(C)cc4)n3)CC2)c1 | 7.943282 | 4 | 29.614 | 25.676 | 14.125375 | -3 | -34.755 | -33.303 |
| NCGC00487736-01 | O=C(NCc1ccccc1)c1nc2sccn2c1-c1cccc(C(F)(F)F)c1 | NaN | 4 | 0 | 1.315 | NaN | 4 | 0 | -3.156 |
| NCGC00489839-01 | CCOC(=O)C1CCCN(C(=O)c2ccc3oc(-c4cccnc4C)nc3c2)C1 | NaN | 4 | 0 | -0.584 | NaN | 4 | 0 | -1.336 |
| NCGC00490399-01 | CCCCCNC(=O)C1CCCN1c1nccc(Nc2ccc(C)cc2)n1 | NaN | 4 | 0 | 1.79 | 3.981072 | 4 | -19.11 | -21.976 |
| NCGC00490335-01 | COc1ccc(Nc2ccnc(N3CCC(NC(=O)c4cccc(Cl)c4)CC3)n2)cc1 | 7.943282 | 2.2 | 66.485 | 59.058 | 14.125375 | -3 | -57.595 | -50.961 |
| NCGC00490338-01 | COc1ccc(Nc2ccnc(N3CCC(NC(=O)c4ccc(Cl)cc4)CC3)n2)cc1 | NaN | 4 | 0 | 0.292 | NaN | 4 | 0 | 1.636 |
| NCGC00490529-01 | COCCNC(=O)C1CCCN1c1nccc(Nc2ccc(C)cc2)n1 | NaN | 4 | 0 | 0.548 | NaN | 4 | 0 | -3.301 |
| NCGC00490397-01 | CSCCNC(=O)C1CCCN1c1nccc(Nc2ccc(C)cc2)n1 | NaN | 4 | 0 | -0.73 | NaN | 4 | 0 | -3.96 |
| NCGC00489844-01 | CCc1ccc(C(C)Nc2nc(N3CCNCC3)nc3ccccc23)cc1 | 3.981072 | 3 | 74.234 | 0 | 11.220185 | 4 | -28.366 | -26.555 |
| NCGC00490398-01 | Cc1ccc(Nc2ccnc(N3CCCC3C(=O)NC3CCCC3)n2)cc1 | NaN | 4 | 0 | 5.88 | NaN | 4 | 0 | -11.046 |
| NCGC00490340-01 | COc1ccc(Nc2ccnc(N3CCC(NC(=O)c4ccc(C)cc4)CC3)n2)cc1 | NaN | 4 | 0 | 0.73 | NaN | 4 | 0 | -2.856 |
| NCGC00492526-01 | Cc1ccc(Nc2ccnc(N3CCCC(C(=O)NCCc4ccc(F)cc4)C3)n2)cc1 | NaN | 4 | 0 | -1.68 | NaN | 4 | 0 | -0.571 |
| NCGC00492511-01 | O=C(NCc1ccccc1)c1nc2cccnc2n(CCc2ccccc2)c1=O | NaN | 4 | 0 | -1.315 | NaN | 4 | 0 | -1.452 |
| NCGC00490608-01 | Cc1ccc(-c2ccccc2CN2CCN(C(=O)NC3CCCC3)CC2)cc1Cl | NaN | 4 | 0 | -2.484 | NaN | 4 | 0 | -7.474 |
| NCGC00490403-01 | Cc1ccc(Nc2ccnc(N3CCCC3C(=O)N3CCC(c4ccccc4)CC3)n2)cc1 | 4.466836 | 4 | 23.5 | -1.68 | 12.589254 | -3 | -103.378 | -95.271 |
| NCGC00492538-01 | Cc1ccc(Nc2ccnc(N3CCCC(C(=O)NCCc4ccccc4)C3)n2)cc1 | NaN | 4 | 0 | -1.169 | NaN | 4 | 0 | -4.492 |
| NCGC00490532-01 | Cc1ccc(Nc2ccnc(N3CCCC3C(=O)N3CCC(Cc4ccccc4)CC3)n2)cc1 | 10 | 2.2 | 69.447 | 60.08 | 12.589254 | -3 | -105.437 | -95.571 |
| NCGC00492889-01 | COc1cccc(C(=O)NCCNC(=O)c2cccc(-c3ccc(C(F)(F)F)cc3)c2)c1 | NaN | 4 | 0 | 0.11 | NaN | 4 | 0 | -2.556 |
| NCGC00492546-01 | Cc1ccc(Nc2ccnc(N3CCCC3C(=O)NCCc3ccccc3)n2)cc1 | NaN | 4 | 0 | -1.753 | 8.912509 | -2.2 | -33.515 | -33.622 |
| NCGC00492710-01 | COc1ccc(C(=O)NCCNC(=O)c2cccc(-c3ccc(SC)cc3)c2)cc1 | NaN | 4 | 0 | -0.11 | NaN | 4 | 0 | -2.323 |
| NCGC00492924-01 | O=C(CNC(=O)c1cccc(-c2ccc(C(F)(F)F)cc2)c1)NCc1ccccc1F | NaN | 4 | 0 | -1.497 | NaN | 4 | 0 | -0.571 |
| NCGC00493002-01 | Cc1ccc(Nc2ccnc(N3CCCC(C(=O)NCCCc4ccccc4)C3)n2)cc1 | NaN | 4 | 0 | -0.511 | 12.589254 | -3 | -93.954 | -88.426 |
| NCGC00493128-01 | O=C(NCc1ccc(F)cc1F)C1CCN(Cc2ccccc2-c2cccnc2)CC1 | NaN | 4 | 0 | 1.023 | NaN | 4 | 0 | -4.715 |
| NCGC00493025-01 | COc1ccc(CC(=O)NCCNC(=O)c2cccc(-c3ccsc3)c2)cc1 | NaN | 4 | 0 | -1.205 | NaN | 4 | 0 | 0.252 |
| NCGC00494501-01 | O=C(CC1c2cccnc2C(=O)N1c1c(Cl)cccc1Cl)NCCN1CCCC1 | NaN | 4 | 0 | 0.329 | NaN | 4 | 0 | -0.561 |
| NCGC00493156-01 | Cc1cc(C)cc(-c2ccc(CN3CCCC(NC(=O)c4ccc(Cl)nc4)C3)cc2)c1 | NaN | 4 | 0 | -0.037 | NaN | 4 | 0 | -1.297 |
| NCGC00494693-01 | CC(Nc1nc(N2CCNCC2)nc2cccc(Cl)c12)c1ccccc1 | 4.466836 | 3 | 87.614 | -2.52 | 12.589254 | -3 | -114.117 | -99.395 |
| NCGC00496026-01 | O=C(NC1CC1)[C@@H]1C[C@H](Oc2ccccc2)CN1C(=O)CCCc1c[nH]c2ccccc12 | NaN | 4 | 0 | -1.57 | NaN | 4 | 0 | -7.203 |
| NCGC00498635-01 | O=C(NCc1cccc(C(F)(F)F)c1)C1CCCN(Cc2cccc(-c3ccc(F)cc3F)c2)C1 | 11.220185 | 4 | 15.884 | 12.966 | 10 | 4 | -21.121 | -23.476 |
| NCGC00498695-01 | COc1ccccc1-c1cccc(CN2CCCC(C(=O)NCc3ccc(C(F)(F)F)cc3)C2)c1 | 0.316228 | 4 | -19.725 | -3.104 | 12.589254 | 4 | -17.635 | -16.971 |
| NCGC00594513-01 | CCNC(=O)c1cc(-c2ccc(Cl)s2)nc2ccccc12 | NaN | 4 | 0 | -1.132 | 0.316228 | 4 | 12.584 | -2.972 |
| NCGC00498744-01 | COc1ccccc1CNC(=O)C1CCCN1Cc1ccc(-c2cc(C)cc(C)c2)cc1 | 14.125375 | 3 | 39.089 | 33.199 | 10 | -3 | -40.34 | -41.009 |

|  | testing results in the cytophatic effect (CPE) assay |
| --- | --- |
|  | testing results in the cytotoxicity Counter-Assay |

**Supplementary Table 8.** Experimental results for all ‘cherry-picked’ compounds retested in CPE and counter-screen assay (8-point assay and in duplicates).

| **Sample ID** | **Smiles** | **AC50 (uM)** | **CC-v2** | **Efficacy** | **Max response** | **AC50 (uM)_counter-screen** | **CC-v2_counter-screen** | **Efficacy_counter-screen** | **Max response_counter-screen** |
| --- | --- | --- | --- | --- | --- | --- | --- | --- | --- |
| NCGC00100643-02 | COc1ccc(Nc2nc(Nc3ccc(OC)cc3OC)nc(N3CCOCC3)n2)c(OC)c1 | 5.011872 | 1.1 | 89.86 | 86.359 | 12.589254 | 4 | -17.148 | -18.203 |
| NCGC00384291-01 | CNCCN(C)c1nc(NC(C)c2ccccc2)c2ccccc2n1 | 6.309573 | 1.2 | 81.769 | 67.935 | NaN | 4 | 0 | -4.007 |
| NCGC00487028-01 | CC(Nc1nc(N2CCN(CCN)CC2)nc2ccccc12)c1ccccc1 | 7.943282 | 2.1 | 123.165 | 78.285 | NaN | 4 | 0 | -11.491 |
| NCGC00419475-01 | CN(C)CCNC(=O)c1ccc2c(c1)C1(CCN(CC3CC3)C1)CN2Cc1ccccc1 | 7.943282 | 1.1 | 98.008 | 91.994 | NaN | 4 | 0 | -7.362 |
| NCGC00104471-02 | CCN(CC)CCNC(=O)c1cc(-c2ccc(Br)s2)nc2ccccc12 | 7.943282 | 2.2 | 95.22 | -0.296 | 14.125375 | -1.1 | -86.512 | -73.121 |
| NCGC00140192-01 | CCc1ccc(Sc2cc(C(=O)NCCCN3CCC(C)CC3)c3ccccc3n2)cc1 | 7.943282 | 1.1 | 89.226 | 85.496 | 14.125375 | -1.2 | -31.425 | -31.58 |
| NCGC00104445-01 | CCCCN(C)CCNC(=O)c1cc(-c2ccc(Cl)s2)nc2ccccc12 | 7.943282 | 1.2 | 54.555 | 40.96 | 14.125375 | -1.2 | -55.72 | -55.326 |
| NCGC00492858-01 | Cc1ccc(Nc2ccnc(N3CCC(C(=O)NCCCc4ccccc4)CC3)n2)cc1 | 8.912509 | 1.2 | 62.015 | 59.328 | 14.125375 | -2.2 | -33.416 | -28.101 |
| NCGC00437792-01 | CN(C)c1cccc(C(=O)NCC2CCN(c3cc(-c4ccc(Cl)cc4)n[nH]3)CC2)c1 | 8.912509 | 4 | 19.477 | 35.887 | 14.125375 | 4 | -22.267 | -21.725 |
| NCGC00496752-01 | Cc1ccc2c(NC(C)c3ccccc3)nc(N3CCNCC3)nc2c1 | 10 | 1.1 | 89.313 | 83.843 | 11.220185 | 4 | -14.377 | -17.361 |
| NCGC00110541-02 | CCCN1CCN(CCCNC(=O)c2cc3c(Cl)nc4ccccc4c3s2)CC1 | 10 | 1.2 | 75.143 | 74.075 | 14.125375 | 4 | -29.648 | -21.157 |
| NCGC00139688-01 | O=C(NCCCN1CCCCC1)c1cc(Sc2ccc(Cl)cc2)nc2ccccc12 | 11.220185 | 1.1 | 99.262 | 94.328 | NaN | 4 | 0 | -7.176 |
| NCGC00110841-01 | Cc1ccc2nc(Cl)c3cc(C(=O)NCCCN4CCC(C)CC4)sc3c2c1 | 11.220185 | 1.1 | 91.075 | 88.716 | NaN | 4 | 0 | -5.302 |
| NCGC00105925-01 | CCCN(CCC)CCNC(=O)c1cc(-c2ccc(Br)s2)nc2ccccc12 | 11.220185 | 1.2 | 81.95 | 76.585 | 14.125375 | 4 | -17.703 | -17.47 |
| NCGC00121825-01 | COc1ccc(-c2cn3c(n2)sc2cc(C(=O)NCCCN4CCCCCC4)ccc23)cc1 | 11.220185 | 1.2 | 70.567 | 68.514 | 10 | -1.4 | -32.335 | -26.281 |
| NCGC00108138-01 | CCCCN(CCCNC(=O)c1cc(Nc2ccc(OC)c(OC)c2)nc2ccccc12)Cc1ccccc1 | 11.220185 | 4 | 11.852 | 10.174 | NaN | 4 | 0 | -3.926 |
| NCGC00377241-01 | Cc1ccc(-c2cccc(CN3CCCC(c4nc5ccc(C)cc5c(=O)[nH]4)C3)c2)c(C)c1 | 12.589254 | 1.1 | 97.954 | 96.745 | NaN | 4 | 0 | -4.117 |
| NCGC00390104-01 | Cc1ccc(C(C)Nc2nc(N3CCNCC3)nc3ccccc23)cc1 | 12.589254 | 1.1 | 92.392 | 90.822 | NaN | 4 | 0 | -11.135 |
| NCGC00120066-01 | CCN(CC)CCNC(=O)c1[nH]c2ccccc2c1Sc1ccc(Cl)cc1 | 12.589254 | 1.1 | 91.961 | 91.174 | NaN | 4 | 0 | -6.95 |
| NCGC00139690-01 | CC1CCCN(CCCNC(=O)c2cc(Sc3ccc(F)cc3)nc3ccccc23)C1 | 12.589254 | 1.1 | 90.753 | 92.851 | NaN | 4 | 0 | -1.863 |
| NCGC00109633-01 | CCCN(CCC)CCCNC(=O)C1CCCN(Cc2nc(-c3ccc(Cl)cc3)oc2C)C1 | 12.589254 | 1.2 | 87.64 | 78.705 | 14.125375 | 4 | -17.699 | -16.48 |
| NCGC00140190-01 | CCc1ccc(Sc2cc(C(=O)NCCCN3CCCC(C)C3)c3ccccc3n2)cc1 | 12.589254 | 1.1 | 86.56 | 84.131 | 14.125375 | 4 | -18.694 | -20.145 |
| NCGC00132818-01 | CCN(CC)CCNC(=O)c1ccc(-c2nc(CN3CCc4ccccc43)c(C)o2)cc1 | 12.589254 | 2.1 | 82.794 | 91.896 | 14.125375 | 4 | -10.025 | -13.235 |
| NCGC00377240-01 | COc1cccc(-c2ccc(CN3CCCC(c4nc5ccc(C)cc5c(=O)[nH]4)C3)cc2)c1 | 12.589254 | 1.4 | 45.428 | 40.349 | 28.183829 | -2.4 | -51.573 | -11.727 |
| NCGC00498993-01 | COc1ccccc1C(=O)NC1CCCN(Cc2cccc(-c3ccc(C)c(Cl)c3)c2)C1 | 12.589254 | 1.2 | 37.56 | 41.818 | 31.622777 | 4 | -32.311 | -12.16 |
| NCGC00113431-01 | CCCCN(CCCC)CCCNC(=O)c1ccc2nc(N3CCCCC3)sc2c1 | 14.125375 | 1.1 | 83.675 | 86.482 | NaN | 4 | 0 | -7.193 |
| NCGC00132814-01 | CCN(CC)CCCNC(=O)c1ccc(-c2nc(CN3CCc4ccccc43)c(C)o2)cc1 | 14.125375 | 1.1 | 81.819 | 97.248 | NaN | 4 | 0 | -0.173 |
| NCGC00120194-01 | COc1ccc2c(Sc3ccc(Cl)cc3)c(C(=O)NCCCN3CCCCC3)[nH]c2c1 | 14.125375 | 1.1 | 81.531 | 90.118 | 14.125375 | -3 | -75.639 | -1.675 |
| NCGC00479261-01 | C[C@@H](Nc1nc(N2CCNCC2)nc2ccccc12)c1ccc(F)cc1 | 14.125375 | 1.1 | 81.301 | 89.118 | 14.125375 | 4 | -12.942 | -15.443 |
| NCGC00427036-01 | O=C(NCCN1CCCCC1)c1ccc(-c2ccc(N3CCSCC3)nc2)cc1 | 14.125375 | 1.1 | 81.074 | 83.378 | NaN | 4 | 0 | -5.878 |
| NCGC00098551-01 | CCn1c2ccccc2c2cc(CN3CCN(Cc4cccc(Cl)c4)CC3)ccc21 | 14.125375 | 1.2 | 79.455 | 88.273 | NaN | 4 | 0 | -3.35 |
| NCGC00389098-01 | CC(Nc1nc(N2CCNCC2C)nc2ccccc12)c1ccccc1 | 14.125375 | 1.2 | 75.562 | 84.36 | NaN | 4 | 0 | -10.058 |
| NCGC00120134-01 | COc1ccc2c(Sc3ccccc3)c(C(=O)NCCCN3CCCCC3)[nH]c2c1 | 14.125375 | 1.2 | 73.449 | 76.007 | 15.848932 | 4 | -18.421 | -19.112 |
| NCGC00109549-01 | CCCCSCCCNC(=O)C1CCN(Cc2nc(-c3ccc(CC)cc3)oc2C)CC1 | 14.125375 | 1.2 | 73.005 | 80.038 | 14.125375 | 4 | -21.335 | -14.601 |
| NCGC00400702-01 | O=C(C[C@@H]1CCNC[C@@H]1Cc1cc(-c2ccc(F)cc2)on1)N1CCC(Cc2ccccc2)CC1 | 14.125375 | 1.2 | 72.596 | 85.233 | 3.981072 | 4 | 9.626 | -3.872 |
| NCGC00440956-01 | Clc1ccccc1C1(c2ccnc(-c3ccccc3)n2)CCNCC1 | 14.125375 | 1.2 | 69.584 | 82.291 | 28.183829 | 4 | -16.355 | -10.185 |
| NCGC00498682-01 | COc1ccccc1-c1cccc(CN2CCCC(C(=O)NCc3ccc(F)cc3)C2)c1 | 14.125375 | 1.2 | 67.172 | 74.104 | 12.589254 | 4 | -29.924 | -12.039 |
| NCGC00139692-01 | CC1CCN(CCCNC(=O)c2cc(Sc3ccc(F)cc3)nc3ccccc23)CC1 | 14.125375 | 1.2 | 65.913 | 84.436 | NaN | 4 | 0 | -8.01 |
| NCGC00449240-01 | Clc1ccc2nc(N3CCC(NCCc4cccnc4)CC3)sc2c1 | 14.125375 | 1.2 | 64.034 | 76.234 | 14.125375 | 4 | -18.146 | -17.213 |
| NCGC00105953-02 | CCCCN(CCCC)CCCNC(=O)c1cc(-c2ccc(Cl)s2)nc2ccccc12 | 14.125375 | 1.2 | 63.824 | 75.768 | 17.782794 | 4 | -16.131 | -12.51 |
| NCGC00121953-01 | O=C(NCCCN1CCCCCC1)c1ccc2c(c1)sc1nc(-c3ccccc3)cn12 | 14.125375 | 1.2 | 63.781 | 70.598 | 14.125375 | -1.2 | -30.183 | -32.509 |
| NCGC00377200-01 | Cc1ccc2nc(C3CCCN(Cc4cccc(-c5cccnc5)c4)C3)[nH]c(=O)c2c1 | 14.125375 | 2.2 | 58.439 | 69.7 | NaN | 4 | 0 | -8.213 |
| NCGC00426433-01 | COc1cccc(-c2ccccc2NC(=O)C2CCN(Cc3ccccc3F)CC2)c1 | 14.125375 | 1.2 | 55.79 | 61.319 | NaN | 4 | 0 | -0.611 |
| NCGC00522637-01 | CC(C)(C)NCc1ccc(Nc2ccnc3cc(Cl)ccc23)cc1O | 14.125375 | 1.2 | 54.851 | 70.062 | 14.125375 | 4 | -29.465 | -33.984 |
| NCGC00140589-02 | COc1ccc(N2CCN(CCCNC(=O)c3cc(Nc4ccc(OC)cc4OC)nc4ccccc34)CC2)cc1 | 14.125375 | 1.2 | 46.734 | 51.725 | 14.125375 | 4 | -22.595 | -22.537 |
| NCGC00493148-01 | COc1ccc(CC(=O)NC2CCN(Cc3ccc(-c4ccccc4OC)cc3)C2)cc1 | 14.125375 | 1.2 | 44.222 | 48.262 | 19.952623 | 4 | -11.413 | -7.42 |
| NCGC00116815-01 | CN(C)c1ccc(-c2nc(-c3cccs3)c(-c3cccs3)[nH]2)cc1 | 14.125375 | 2.2 | 40.59 | 56.407 | NaN | 4 | 0 | -20.003 |
| NCGC00498661-01 | O=C(NCc1ccccc1Cl)C1CCCN(Cc2cccc(-c3ccc(F)cc3F)c2)C1 | 14.125375 | 1.2 | 37.261 | 44.175 | 14.125375 | -2.2 | -33.487 | -18.1 |
| NCGC00120248-01 | CCC1CCCCN1CCCNC(=O)c1sc2nc(C)cc(C)c2c1-n1cccc1 | 14.125375 | 2.2 | 35.65 | 53.233 | NaN | 4 | 0 | -8.835 |
| NCGC00140413-01 | Cc1ccccc1CN1C(=O)c2ccccc2Sc2ccc(C(=O)NCCCN3CCOCC3)cc21 | 14.125375 | 1.2 | 32.206 | 40.818 | 14.125375 | 4 | -14.136 | -15.628 |
| NCGC00126272-01 | CCOc1ccc(NC(=O)c2cnn(-c3ccc(C)c(Cl)c3)c2C2CCNCC2)cc1 | 14.125375 | 1.2 | 30.062 | 33.913 | 19.952623 | 4 | -11.432 | -9.03 |
| NCGC00480765-01 | CC(C)(Nc1nc(N2CCNCC2)nc2ccccc12)c1ccccc1 | 14.125375 | 4 | 23.533 | 39.734 | NaN | 4 | 0 | -4.437 |
| NCGC00510147-03 | CC(C)(C)N1CCC(c2ccccc2)(c2ccccc2)CC1 | 14.125375 | 4 | 20.165 | 23.622 | NaN | 4 | 0 | -4.277 |
| NCGC00109615-01 | CCCCNC(=O)C1CCCN(Cc2nc(-c3ccc(Cl)cc3)oc2C)C1 | 14.125375 | 4 | 19.996 | 24.874 | 14.125375 | 4 | -27.647 | -28.809 |
| NCGC00498749-01 | Cc1cc(C)cc(-c2ccc(CN3CCCC3C(=O)NCc3cccc(F)c3)cc2)c1 | 14.125375 | 4 | 15.122 | 18.111 | 14.125375 | 4 | -24.311 | -26.713 |
| NCGC00140464-01 | Cc1cccc(N2CCN(C(=O)C3CCN(Cc4nc(-c5ccc(Cl)cc5)oc4C)CC3)CC2)c1C | 14.125375 | 4 | 12.144 | 13.329 | NaN | 4 | 0 | 2.266 |
| NCGC00427056-01 | Cn1cc(CN2CCCC(COc3ccccc3C(=O)N3CCCCC3)C2)c2ccccc21 | 14.125375 | 4 | 9.96 | 11.925 | NaN | 4 | 0 | -11.185 |
| NCGC00419419-01 | CN1CCCC2(C1)CN(Cc1cccnc1)c1ccc(C(=O)NCC3CCCCC3)cc12 | 25.118864 | 2.1 | 124.124 | 97.373 | NaN | 4 | 0 | -3.806 |
| NCGC00416827-01 | COc1cc2c(cc1OCC1CC1)NC(=O)C21CCN(c2cc(NC3CC3)ncn2)CC1 | 25.118864 | 2.2 | 73.039 | 57.54 | NaN | 4 | 0 | -10.135 |
| NCGC00489837-01 | Cc1ncccc1-c1nc2cc(C(=O)NCCCc3ccccc3)ccc2o1 | 25.118864 | 2.2 | 50.97 | 38.177 | NaN | 4 | 0 | -8.085 |
| NCGC00109669-01 | Cc1oc(-c2cccc(Cl)c2)nc1CN1CCCC(C(=O)NCCc2ccc(Cl)cc2)C1 | 25.118864 | 3 | 43.391 | 34.045 | 14.125375 | -1.2 | -44.328 | -45.623 |
| NCGC00411503-01 | Cc1noc2c(NC3CCOCC3)cc(-c3cccc(C(F)(F)F)c3)nc12 | 25.118864 | 4 | 22.792 | 17.282 | 25.118864 | -2.2 | -58.79 | -44.012 |
| NCGC00116825-01 | Oc1ccccc1-c1nc(-c2cccs2)c(-c2cccs2)[nH]1 | 28.183829 | 4 | 15.447 | 11.516 | 14.125375 | 4 | -23.827 | -7.093 |
| NCGC00108262-01 | Cc1ccc(C)c(Nc2cc(C(=O)NCCCN(C)C3CCCCC3)c3ccccc3n2)c1 | NaN | 4 | 0 | 58.938 | 14.125375 | -1.2 | -41.828 | -45.15 |
| NCGC00113433-01 | CCCCN(C)CCCNC(=O)c1ccc2nc(N3CCCCC3)sc2c1 | NaN | 4 | 0 | 6.989 | NaN | 4 | 0 | -8.01 |
| NCGC00137630-01 | COCCCNC(=O)C1CCN(Cc2cc3ccccc3n2Cc2cccc(Cl)c2)CC1 | NaN | 4 | 0 | 1.642 | NaN | 4 | 0 | -9.06 |
| NCGC00137604-01 | COc1cccc(CNC(=O)C2CCN(Cc3cc4ccccc4n3Cc3ccccc3)CC2)c1OC | NaN | 4 | 0 | 4.752 | NaN | 4 | 0 | -4.814 |
| NCGC00140311-01 | COc1ccc(CCNC(=O)c2cc(Nc3ccccc3)nc3ccccc23)cc1OC | NaN | 4 | 0 | 2.557 | 8.912509 | 4 | -11.789 | -10.943 |
| NCGC00498641-01 | O=C(Nc1cccc(Cl)c1)C1CCCN(Cc2cccc(-c3ccc(F)cc3F)c2)C1 | NaN | 4 | 0 | 7.041 | NaN | 4 | 0 | 2.165 |

|  | testing results in the cytophatic effect (CPE) assay |
| --- | --- |
|  | testing results in the cytotoxicity Counter-Assay |
